# Estimating the effectiveness of control actions on African swine fever transmission in commercial swine populations in the United States

**DOI:** 10.1101/2022.09.04.506538

**Authors:** Abagael L. Sykes, Jason A. Galvis, Kathleen C. O’Hara, Cesar Corzo, Gustavo Machado

## Abstract

Given the proximity of African swine fever (ASF) to the U.S., there is an urgent need to better understand the possible dissemination pathways of the virus within the U.S. swine industry and to evaluate mitigation strategies. Here, we extended *PigSpread*, a farm-level spatially-explicit stochastic compartmental transmission model incorporating six transmission routes including between-farm swine movements, vehicle movements, and local spread, to model the dissemination of ASF. We then examined the effectiveness of control actions similar to the ASF national response plan. The average number of secondary infections during the first 60 days of the outbreak was 49 finisher farms, 17 nursery farms, 5 sow farms, and less than one farm in other production types. The between-farm movements of swine were the predominant route of ASF transmission with an average contribution of 71.1%, while local spread and movement of vehicles were less critical with average contributions of 14.6% and 14.4%. We demonstrated that the combination of quarantine, depopulation, movement restrictions, contact tracing, and enhanced surveillance, was the most effective mitigation strategy, resulting in an average reduction of 79.0% of secondary cases by day 140 of the outbreak. Implementing these control actions led to a median of 495,619 depopulated animals, 357,789 diagnostic tests, and 54,522 movement permits. Our results suggest that the successful elimination of an ASF outbreak is likely to require the deployment of all control actions listed in the ASF national response plan for more than 140 days, as well as estimating the resources needed for depopulation, testing, and movement permits under these controls.

## 1. Introduction

In July 2021, African swine fever (ASF) was identified in the North American continent for the first time since 1984 (Costard et al., 2009), with outbreaks now reported in domestic swine farms of both the Dominican Republic (Gonzales et al., 2021) and Haiti (OIE, 2021). This occurred alongside ongoing outbreaks across Europe and Asia in both domestic and feral swine (Mighell and Ward, 2021; You et al., 2021; de la Torre et al., 2022; Iscaro et al., 2022; Penrith and Kivaria, 2022) which have been characterized, in most cases, by high mortality and morbidity (EFSA Panel on Animal Health and Welfare (AHAW), 2010, 2014; Gallardo et al., 2017). The transmission of ASF between domestic swine farms is primarily driven by direct transmission via contact with infected swine, often introduced to commercial farms via swine movements, while close contact with feral swine has been associated with spillover events (Vergne et al., 2015; Miller et al., 2017; Podgórski and Śmietanka, 2018; Ferdousi et al., 2019; Gebhardt et al., 2021; Yoo et al., 2021). ASF dissemination is also positively correlated with the spatial proximity between farms, which facilitates the local spread of ASF over short distances and the possible formation of spatial clusters (Vergne et al., 2015; Yoo et al., 2021). In particular, the movement of contaminated personnel and/or equipment between farms at short distances has been previously associated with local spread (Bellini et al., 2016; Zani et al., 2019; Gao et al., 2021; Chuchard et al., 2022). Additionally, the potential for ASF to survive on contaminated surfaces, such as transportation vehicles, for several days could lead to transmission via indirect contact (Y. Li et al., 2020; Yoo et al., 2021; Adedeji et al., 2022; Galli et al., 2022), another important route in ASF dissemination (Kim et al., 2021; Yoo et al., 2021; Cheng and Ward, 2022; Galli et al., 2022).

With heavy economical losses associated with ASF evident across the globe (Mason-D’Croz et al., 2020; Niemi, 2020; Weaver and Habib, 2020; Nguyen-Thi et al., 2021; You et al., 2021), mathematical-based modeling approaches have been utilized to estimate and forecast ASF epidemic trajectories, as well as to assess the effectiveness and cost of control actions (Nigsch et al., 2013; Korennoy et al., 2014; Halasa, Bøtner et al., 2016; Mur et al., 2018; Andraud et al., 2021; EFSA et al., 2021; Hayes et al., 2021; Machado et al., 2021; Yoo et al., 2021; Ezanno et al., 2022). In countries with previous ASF outbreaks, transmission models can be directly calibrated with outbreak data, allowing for the estimation of transmission parameters (e.g., the transmission coefficient for between-farm animal movements) (Mur et al., 2018; Akhmetzhanov et al., 2020; Hayes et al., 2021; Yoo et al., 2021) however, ASF-free countries lack the disease-related data necessary for model calibration. In such cases, simulation-based approaches can be used to examine the possible transmission and dissemination dynamics of an ASF introduction.

Epidemiological parameters and knowledge regarding the structure of the swine industry are pivotal in the characterization of disease dissemination and the evaluation of control actions (Halasa et al., 2020; Hayes et al., 2021). In this study, we have collected detailed data on domestic swine populations including farm-level demographics, and between-farm swine and vehicle movement data to develop a stochastic, farm-level, compartmental ASF transmission model for the southeastern U.S.. To address the lack of epidemiological parameters regarding ASF dissemination dynamics and outbreaks in the U.S., we calibrated the transmission rates and surveillance effectiveness parameters using historical porcine epidemic diarrhea virus (PEDV) data from the first recorded outbreak in the study population (Machado et al., 2019). From our model, we estimated the volume of secondary infections, quantified the contribution of six transmission routes, measured the spatial distance from seeded to secondary infections, and assessed the impact of between-company dissemination of ASF. Importantly, we also examined the effectiveness of control actions based on the response strategy detailed in the U.S. Department of Agriculture (USDA) ASF response plan (The Red Book) (USDA, 2020), but with key differences, such as stricter movement controls and depopulation strategies, and estimated the resources required under the most effective control strategy, including the number and relative compensation cost of depopulated animals, the quantity of diagnostic tests performed, and the volume of movement permits required for swine and vehicle movements. With the outputs of this model we address the lack of real-data driven transmission models of ASF spread within the U.S..

## 2. Material and Methods

In the following sections we describe the farm-level population data, the construction of the between-farm swine and vehicle movement networks, the modification of our previous stochastic transmission model *PigSpread* (Galvis, Corzo and Machado, 2021; Galvis, Corzo et al., 2022; Galvis, Jones et al., 2022) to accommodate ASF epidemiological parameters, and the evaluation of ASF control actions listed in The Red Book (USDA, 2020).

### 2.1 Data source

The data used in this model was collected from the Morrison Swine Health Monitoring Project (MSHMP) (MSHMP, 2022), and included a total of 2,294 farms from three swine production companies in the southeastern U.S., identified as A (n = 1,745), B (n = 228), and C (n = 321) for confidentiality purposes. The data consisted of: i) geolocation; ii) the unique premises identification number; iii) production type (sow^1^, nursery, finisher^2^, gilt isolation unit^3^, and boar stud); and iv) swine capacity, which accounted for animals at the nursery age and older. In total the data included 1,459 (63.6%) finisher farms, 468 (20.4%) nursery farms, 328 (14.3%) sow farms, 33 (1.4%) gilt isolation units and six (0.3%) boar studs. The median swine capacity across all production types was 3,200 (IQR: 2,201-5,135). In addition, between-farm swine movement records from January 1st, 2020, to December 31st, 2020, were provided for each company and included information on: i) movement dates, ii) origin and destination farms, iii) the number of swine moved, and iv) the movement purpose (e.g., weaning, replacement gilts). Company A also provided global positioning system (GPS) data of four types of transportation vehicles from January 1st, 2020, to December 31st, 2020, described in more detail in section 2.2.

### 2.2 Swine movement and transportation vehicle networks

Using the between-farm swine movement data, we created a daily contact network with directed edges from the origin farm(s) to the destination farm(s). In total 731 (1.7%) movements were removed from the network due to missing data regarding either of the following: i) the farm location, ii) premises identification number, iii) the number of swine moved, or iv) the movement purpose.

Using the GPS data of 398 individual vehicles from company A, which included 76% of the farms in the study area, we constructed four daily directed contact networks to capture the potential for indirect transmission via vehicles moving swine between farms (swine vehicles, n=118), vehicles moving swine from farms to slaughterhouses (market vehicles, n= 89), vehicles delivering feed to farms (feed vehicles, n=159), and vehicles moving loading crews between farms (crew vehicles, n=32). Here we used an approach proposed by Galvis et al., 2022 to reconstruct daily networks. A more detailed description is provided in Supplementary Material Section A. In total there were 3,182,144 edges in the feed vehicle network, 386,730 edges in the pig vehicle network, 35,854 edges in the market vehicle network, and 109,675 edges in the crew vehicle networks (Galvis, Corzo et al., 2022). Further summary metrics for the networks are provided in Galvis, Corzo et al., (2022).

### 2.3 Model calibration

As the U.S. remains free of ASF at the time of writing, there was no outbreak data available to calibrate ASF transmission dynamic parameters. Instead, we used PEDV outbreak data from the first introduction of the virus in the study region in 2013, as a proxy (USDA, 2014; Machado et al., 2019), which allowed for the evaluation and reproduction of the transmission dynamics of a novel pathogen introduced into a susceptible population.

Using an Approximate Bayesian Computation (ABC) rejection algorithm (Hartig *et al*., 2011; Minter and Retkute, 2019), we identified particles (sets of transmission parameters produced by the model) that reproduced the temporal distribution of PEDV cases according to a tolerance acceptance error (Supplementary Material Section B, Table S1) (Sisson et al., 2007). We drew posterior probabilities for selected parameters from these particles, including transmission rate parameters (*β_n_*, *β_s_*, *β_p_*, *β_m_*, *β_f_*, *β_c_* and *β_l_*); and the effectiveness of passive surveillance for each production type (*L_sow_*, *L_nursery_* and *L_finisher_*), the results of which are presented in Supplementary Material Section B Figures S1-S5 and Table S3. An in-depth description of the PEDV calibration is presented in Supplementary Material Section B.

### 2.4 Baseline transmission model (No control scenario)

Here we modified *PigSpread* (Galvis, Corzo et al., 2022) to accommodate an ASF compartmental model which included the following four mutually exclusive farm-level health states, Susceptible (S), Exposed (E), Infected (I), and Detected (D) (SEID) (Figure 1) and six transmission routes: i) swine movements; ii) swine vehicles; iii) market vehicles; iv) feed vehicles; v) crew vehicles; and vi) local spread, which accounts for mechanical transmission via personnel, equipment and wildlife/pests. The swine and vehicle movements were incorporated as directed temporal networks while the local spread was incorporated using a gravity model with a varied cutoff from one km to 25 km. Further details about the transmission routes including local spread are available in Supplementary Material Section C.

**Figure 1.**
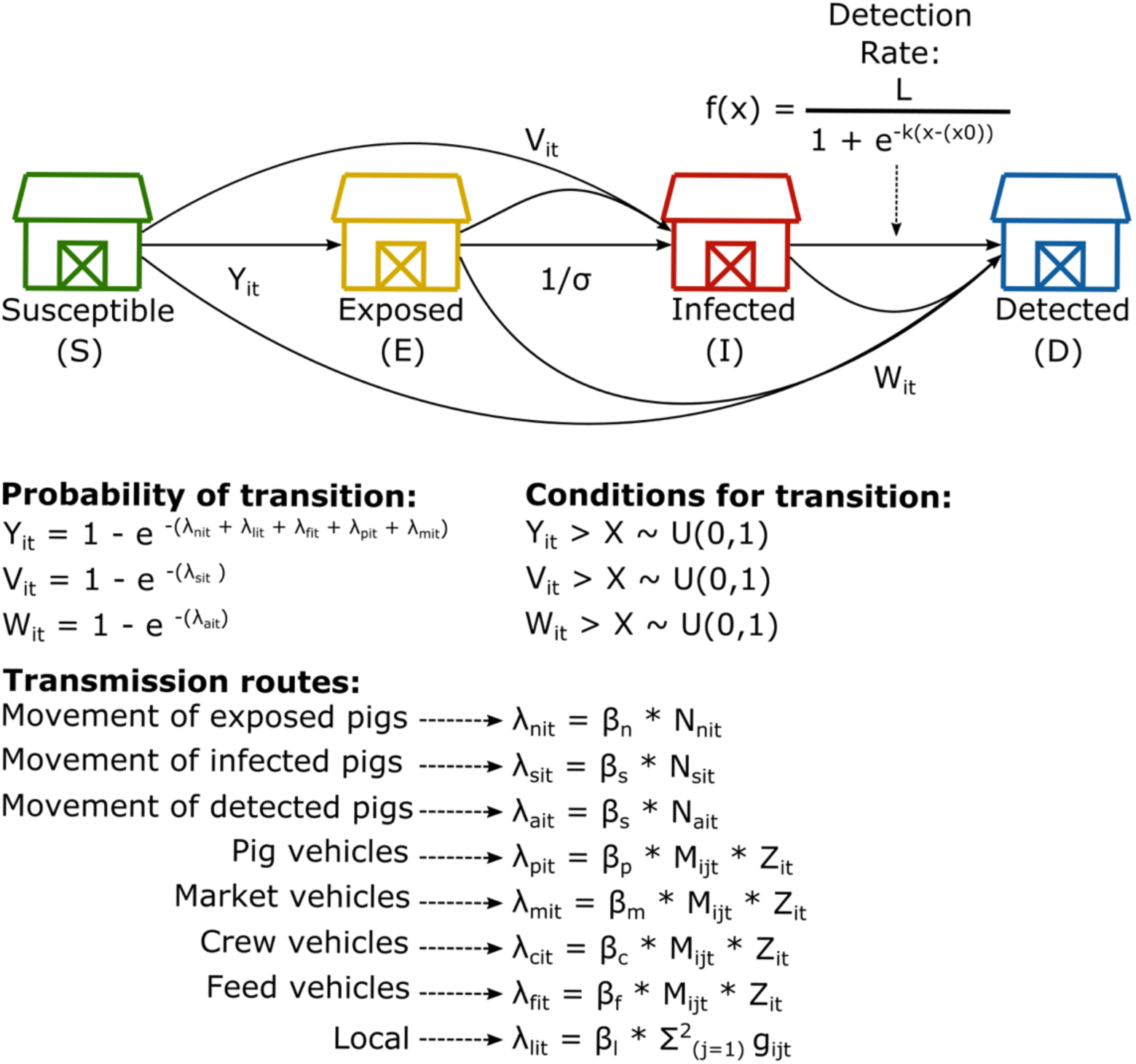
Schematic representation of the SEID ASF model. The schematic demonstrates the transition of farms through the four compartments: Susceptible (S), Exposed (E), Infected (I), and Detected (D). *Y_it_* represents the probability of a farm moving from the susceptible compartment into the exposed compartment; *V_it_* represents the probability of a farm moving from the susceptible or exposed to the infected compartment, and *W_it_* represents the probability of a farm moving from either the susceptible, exposed or infected compartment to the detected compartment. *λ* represents the force of infection for each transmission route and *β* represents the transmission rate for each route.

To account for the variation of between-farm movements we varied the start date and initial seed infection for each simulation. Start dates were sampled between January 1st, 2020, to December 31st, 2020, while each farm in the study population was the initial seeded infection for 100 simulations, resulting in 229,400 repeats in total. To mimic long-duration outbreaks, we ran each simulation for 140 days and collected information at each time step *t,* where *t* → *t +* 1 is one day (Supplementary Material Section C, Figure S6). As our model was implemented at the farm level, we assumed that animal populations within farms remained constant unless purposefully depopulated within the control scenarios (section 2.5). Additionally, as no control actions were implemented in the no control scenario, it was assumed that farms in this scenario could move swine freely regardless of the compartment in which the farm resides. Further details of the model dynamics and sensitivity analyses, including the comparison of sensitivity results are available in Supplementary Material Section C.

### 2.5 Control scenarios

We assessed the impact of five ASF control scenarios (Table 1), which incorporated combinations of the control actions based on the USDA’s ASF response plan, The Red Book (USDA, 2020), detailed below. These control actions were implemented once the first ASF case was detected in a domestic swine farm (USDA, 2020). Below we describe each strategy and note differences between the strategy modeled and the current USDA response plan. Additionally, Supplementary Material Section D Table S4 describes the interaction between the control actions and the transmission routes.

**Table 1.** Control scenarios applied to an ASF outbreak in the U.S.

| **Scenario** | **Depopulation**<br>**of infected**<br>**farms** | **Standstill** | **Contact tracing** | **Depopulation**<br>**of direct**<br>**contacts** | **Control**<br>**areas and**<br>**surveillance**<br>**zones** |
| --- | --- | --- | --- | --- | --- |
| **1** | Yes | No | No | No | No |
| **2** | Yes | Yes | No | No | No |
| **3** | Yes | Yes | Yes | No | No |
| **4** | Yes | Yes | Yes | Yes | No |
| **5** | Yes | Yes | Yes | Yes | Yes |

#### Standstill

The 72-hour movement standstill was defined as a “stop” in all between-farm swine and vehicle movements^4^ (USDA, 2020).

#### Depopulation

Once a domestic swine farm was detected, the farm, including all swine present on the premises, was scheduled for depopulation immediately. To reflect the limited resources available during an outbreak, it was assumed that a maximum of four farms could be depopulated per day (state animal health official, personal communication, 2022), introducing a delay in depopulation whose length was relative to the number of farms awaiting depopulation. Once depopulated, farms were moved to a Removed (R) compartment for 30 days (USDA-APHIS-VS 2022, personal communication, 2022) to simulate a necessary period of cleaning and disinfection prior to repopulation of the farm. After the end of the 30 days, we assumed that all farms had been repopulated and moved back to the S compartment. In control scenarios four and five (Table 1), depopulation was also implemented on direct contacts of the detected farms^5^ regardless of their infection status and these direct contacts were subject to the same depopulation delay as the detected farms.

#### Contact tracing

Contact tracing was defined as identifying farms that were in direct contact with a detected farm in the previous 30 days and indirect contact with the detected farm in the previous 15 days. Both swine and vehicle networks were traced back one step in the contact chain. Farms identified as contacts were then tested three times at six-day intervals (USDA, 2020) starting the day after they were identified as contacts. As direct contact farms are depopulated in control scenarios four and five (Table 1) only the indirect contacts were subjected to testing under the contact tracing control action.

#### Control area and surveillance zones

The control areas (CA) and surveillance zones were defined as areas of increased surveillance and movement permitting around detected farms. The CA consisted of a 3 km infected zone (IZ) and a 2 km buffer zone (BZ), which was then surrounded by a 5 km surveillance zone (SZ) (USDA, 2020). Both the CA and SZ were in place for 30 days after the detection of an infected domestic swine farm. All farms within the IZ were tested every three days for the first two tests, and then every six days, while all farms within the BZ were tested every six days. In contrast, only a sample of the farms in the SZ were tested; see Supplementary Material D for the sampling design utilized to sample farms in the SZ (Cannon, 2001), and the selected farms were tested every 15 days (USDA, 2020). In our model assumptions, it was possible for farms to be within two or more zones (i.e., IZ, BZ or SZ) during the same period. These farms would be tested under the testing schedule for the highest priority zone they were situated within. From highest to lowest, the priority of testing was as follows: IZ, BZ, indirect contacts, and SZ. Farms in multiple zones would only be released from restrictions once the last CA or SZ they were in was lifted. Movement permits were required for any movement of swine or vehicles to and from farms within the IZ and BZ, as well as farms identified as indirect contacts, for 30 days and 15 days, respectively.

It is pertinent to note that the start date and particle sampled for each simulation in the no control scenario were also used for the respective simulation number in the control scenarios to ensure the difference in cases observed was as a result of the control action rather than differences in animal movements by day or transmission parameters.

### 2.6 Model outputs

To capture the dynamics of an ASF outbreak, our model outputs reflect only the simulations which resulted in secondary infections. All results are shown for up to day 60 and day 140 post-ASF introduction, providing a short-term and a long-term perspective of ASF dissemination and control. Model outputs were generated to characterize ASF outbreak dynamics including: i) the number of secondary cases; ii) the contribution of the six transmission routes to ASF dissemination; iii) between-company dissemination of ASF; and iv) the distance between initially seeded farms and secondary infections. The accumulated number of secondary cases was calculated using the time point at which farms transitioned out of the S compartment, regardless of which compartment they transitioned to. The between-company dissemination and distance between seeded and secondary cases were built upon this result. To assess the contribution of the different transmission routes to the dissemination of ASF, we calculated the daily proportional force of infection of the transmission routes for each farm that became exposed, infected, or detected (i.e., moved into the E, I, or D compartment after exposure).

The effectiveness of the five control scenarios was presented via the following outputs: i) number of secondary cases; and ii) the number of animals depopulated and the associated compensation cost under each control scenario. The number of diagnostic tests and volume of movement permits required under the most effective control scenario was also presented. Briefly, the compensation cost of depopulation was calculated using the number of depopulated animals in each production stage and their respective current market-value (USDA, 2022a). The number of diagnostic tests required was calculated assuming a within-farm sampling methodology detailed in The Red Book (USDA, 2020) and Cannon (2001), and pooling the samples by groups of five. We additionally calculated the number of tests required if animals were tested individually. Lastly, the number of permits required was calculated assuming that each movement to or from farms in the IZ and BZ, or movements to or from indirect contact farms, required an individual single-day permit.

## 3. Results

### 3.1 ASF dissemination

From the 229,400 simulations generated by the no control scenario, 91.9% (210,727) resulted in secondary infections. 133,629 of these simulations were seeded in finisher farms, 46,297 in nursery farms, 29,056 in sow farms, and 1,745 were seeded in other production types (gilt isolation and boar stud). In 60 days of ASF dissemination, the average number of secondary infections was 72 (±100, with a lower bound of zero) farms of which 49 (±69, with a lower bound of zero) were in finisher farms, 17 (±24, with a lower bound of zero) were in nursery farms, 5 (±9, with a lower bound of zero) were in sow farms, and 0.07 (±0.57, with a lower bound of zero) were in other production types. The number of secondarily infected farms over 140 days of ASF dissemination was, on average, 810 (±523), which included 543 (±347) infected finisher farms, 191 (±123) infected nursery farms, 74 (±53) infected sow farms, 2 (±3) infected gilt isolation units and 0.2 (±0.5) infected boar studs (Supplementary Material Section E, Figure S8). Sensitivity analyses comparing the mean number of secondary cases were particularly influenced by the date of initial infection; the length of the ASF latent period; the maximum distance for local spread; and the transmission rates for local spread, swine vehicles, feed vehicles and crew vehicles. Results of the sensitivity analyses can be found in Supplementary Material Section E, Figures S9 to S12.

### 3.2 Contribution of transmission routes for baseline model (no control scenario)

Over the first 60 days of ASF dissemination, the dominant transmission route was the between-farm movement of infected swine, whose contribution to dissemination varied from 65.5% on day two of the outbreak to 37.5% on day 60 (Figure 3), with an overall average of 47.7%. The movement of exposed but undetected swine was the second most relevant transmission route, going from no contribution on day two to 23.8% on day 60, with an overall average of 23.3%. The local spread was particularly important during the first days of the outbreak, contributing to 31.7% of ASF dissemination on day two, which decreased to 9.8% by day ten, but increased to 17.3% by day 60, with overall average of 14.6%. Individually, transportation vehicles had limited contributions to ASF dissemination, but the combined contributions varied from 3.3% on day two to 21.3% on day 60 (Figure 3), with a combined overall average of 14.4%. Lastly, the lowest contribution was from the movement of detected swine, which had no contribution on day two and a contribution of 0.1% on day 60, with an overall average of 0.1%.

**Figure 2.**
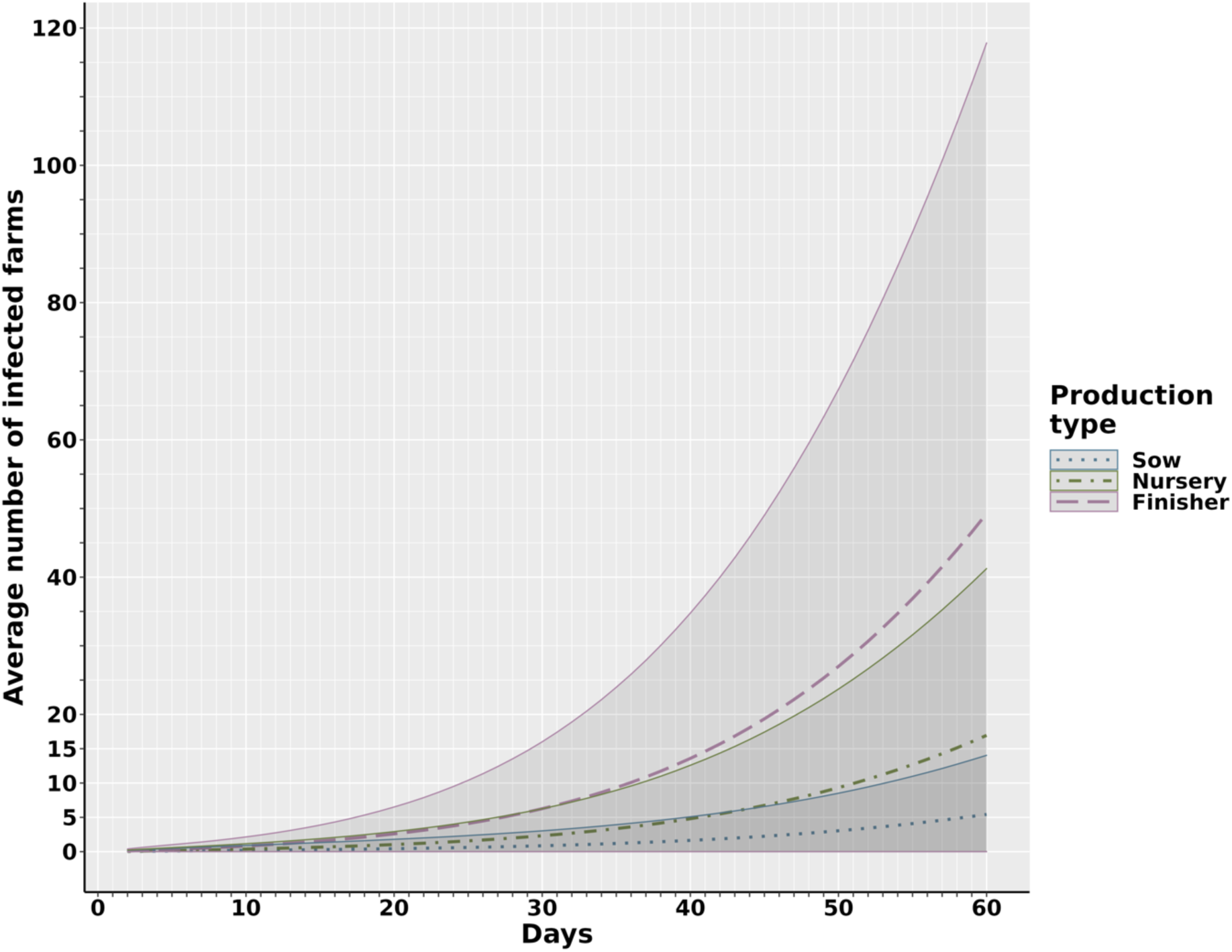
The average number of secondary ASF infections by farm type, accumulated over 60 days of simulated outbreaks. The y-axis represents the average number of secondary infections (farms). The dashed and dotted lines represent the mean at time *t* (days) for each swine production type while the solid lines and shaded areas represent the standard deviation at time *t*.

**Figure 3.**
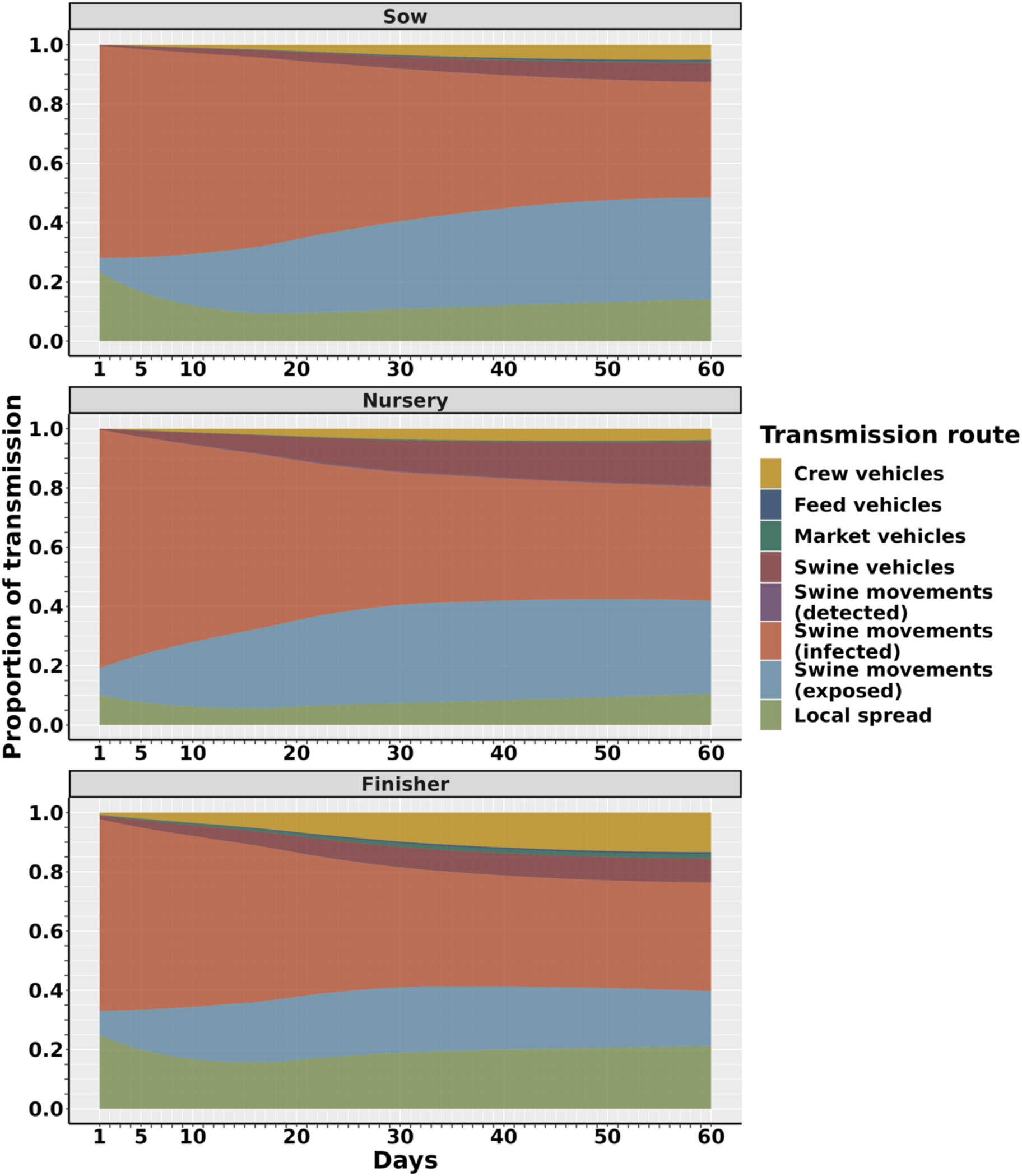
The average contribution of six transmission routes of ASF dissemination in each farm type over 60 days of outbreaks. The y-axis represents the mean proportion of each transmission route on a given day for each production type, which was averaged at each time point and smoothed using a generalized linear model. Transmission of ASF through the movement of swine has been separated into the movement of exposed, infected, and detected swine to reflect the influence of these compartments of ASF transmission. Transmission starts on day two as day one was the time step at which the initial seeded infections were introduced.

Over 140 days, the movements of infected swine, the movements of exposed swine, and local spread continued to lead to transmission with average contributions of 44.7%, 21.2%, and 16.4% at day 140, respectively, while the combined average contribution of the four vehicle movement types was 18.2% (Supplementary Material Section E Figure S13). Details about the average contribution by production type for both 60 and 140 days are available in Supplementary Material Table Section E S5 and S6.

Additional results concerning the average distance between seeded and secondary infections, and the between-company dissemination of ASF are presented in Supplementary Material Section E Figures S14 and S17.

### 3.5 Control scenarios

#### 3.5.1 Effectiveness of control scenarios

To assess the effectiveness of the control actions, the results from each control scenario were benchmarked against the baseline model where no control actions were applied, also referred to as the no control scenario (Figure 4 and Supplementary Material Section E, Table S7). The results of scenario one, which considers quarantine and depopulation of detected farms, demonstrate a median overall reduction in secondary cases of 11.1% by day 60, halting 20% of secondary infections in sow farms, 16.1% of infections in nursery farms and 10.0% of secondary cases in finisher farms. When the 72-hour standstill was added to the quarantine and depopulation of detected farms in scenario two, a median of 14.5% of secondary infections were reduced when compared with the no control scenario, with reductions of 23.5% in sow farms, 20.0% in nursery farms, and 13.6% in finisher farms (Supplementary Material Section E, Table S7).

**Figure 4.**
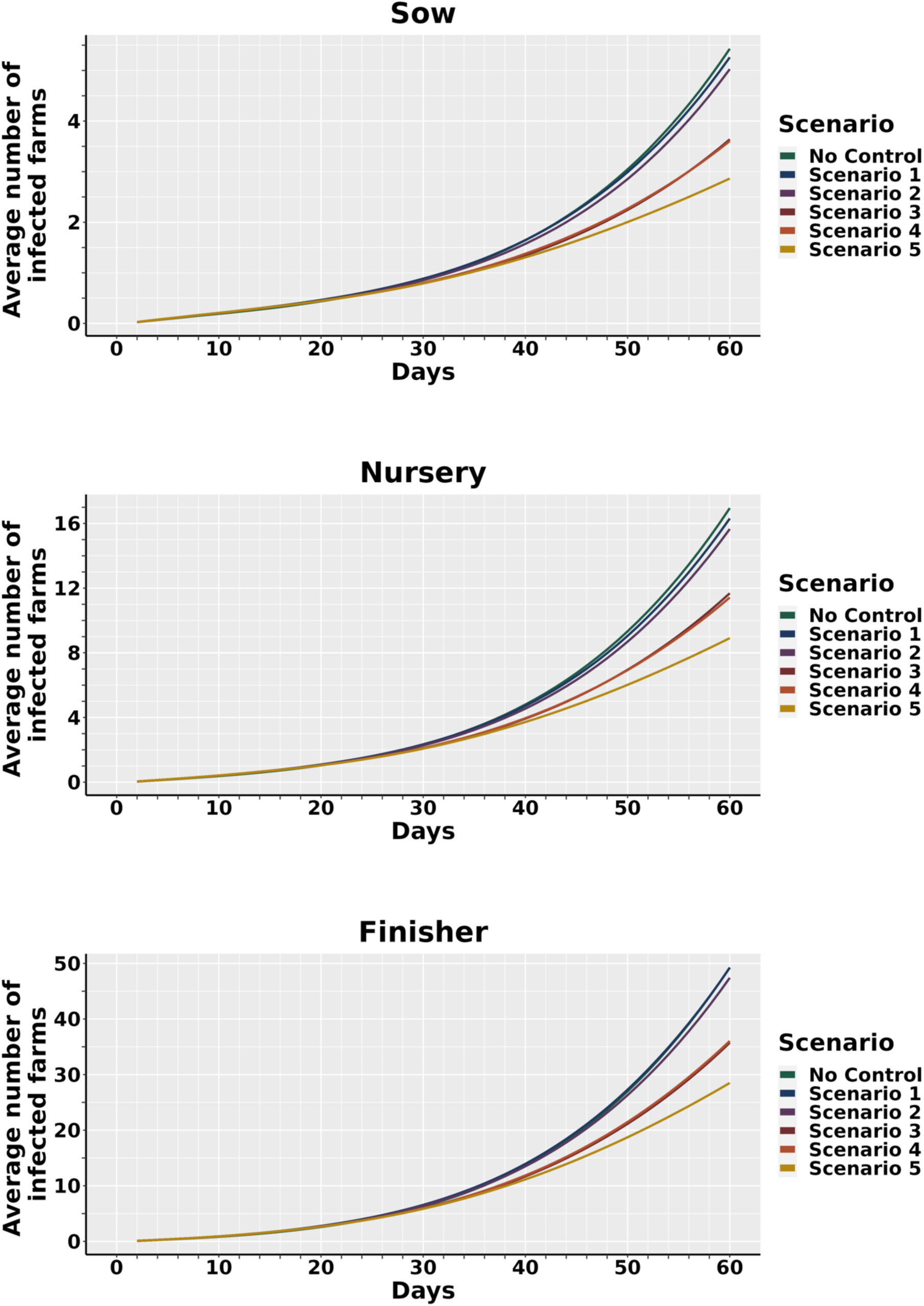
The average number of cumulative infections over 60 days of an outbreak under five different control scenarios. The colored lines represent the mean prevalence after *t* days, under each control scenario. Control scenarios were implemented upon detection of the first ASF case and are described in detail in Table 1.

Scenarios three to five showed a substantial reduction in secondary infections compared to scenarios one and two (Figure 5 and Supplementary Material Section E, Table S7). In scenario three, which included contact tracing of direct contacts (from swine movements) and indirect contacts (from vehicle movements) in addition to the actions described in scenarios one and two, the number of secondary infections was reduced by a median of 34.4%. By farm type, scenario three reduced sow infections by a median of 43.2%, nursery infections by a median of 39.4%, and finisher infections by a median of 33.3%. Scenario four, in which the depopulation of direct contacts is considered, showed very similar overall performance as scenario three with a median decrease in the number of secondary cases of 33.5%, compared to the no control scenario. Here the number of cases in sow farms decreased by a median of 43.8%, in nursery farms decreased by a median of 40.0%, and in finisher farms decreased by a median of 32.1%. Scenario five, where all the control actions including control areas and surveillance zones were implemented, showed the best performance compared to all previous scenarios, with a median decrease of 47.4% in the number of secondary cases compared to the no control scenario. The number of cases in sow farms decreased by a median of 50.0%, nursery farms decreased by a median of 50.0%, and finisher farms decreased by a median of 45.1% (Table S7). In Supplementary Material Figure S18 we show the reduction in the number of secondary cases for each scenario at 140 days. Briefly, for scenario five, the median decrease in total secondary infections at 140 days was 79.0% compared to the no control scenario, with a median decrease in infected sow farms of 82.1%, a median decrease in infected nursery farms of 79.3%, and a median decrease in infected finisher farms of 76.6%. However, while there was a substantial decrease in the number of secondary cases under scenario five, only 29.0% of simulations were controlled by day 140 (i.e. no cases after day 135). Visualization of the total secondary cases at 60 and 140 days across the simulations for each control scenario is available in Supplementary Material Section E, Figures S19 and S20.

**Figure 5.**
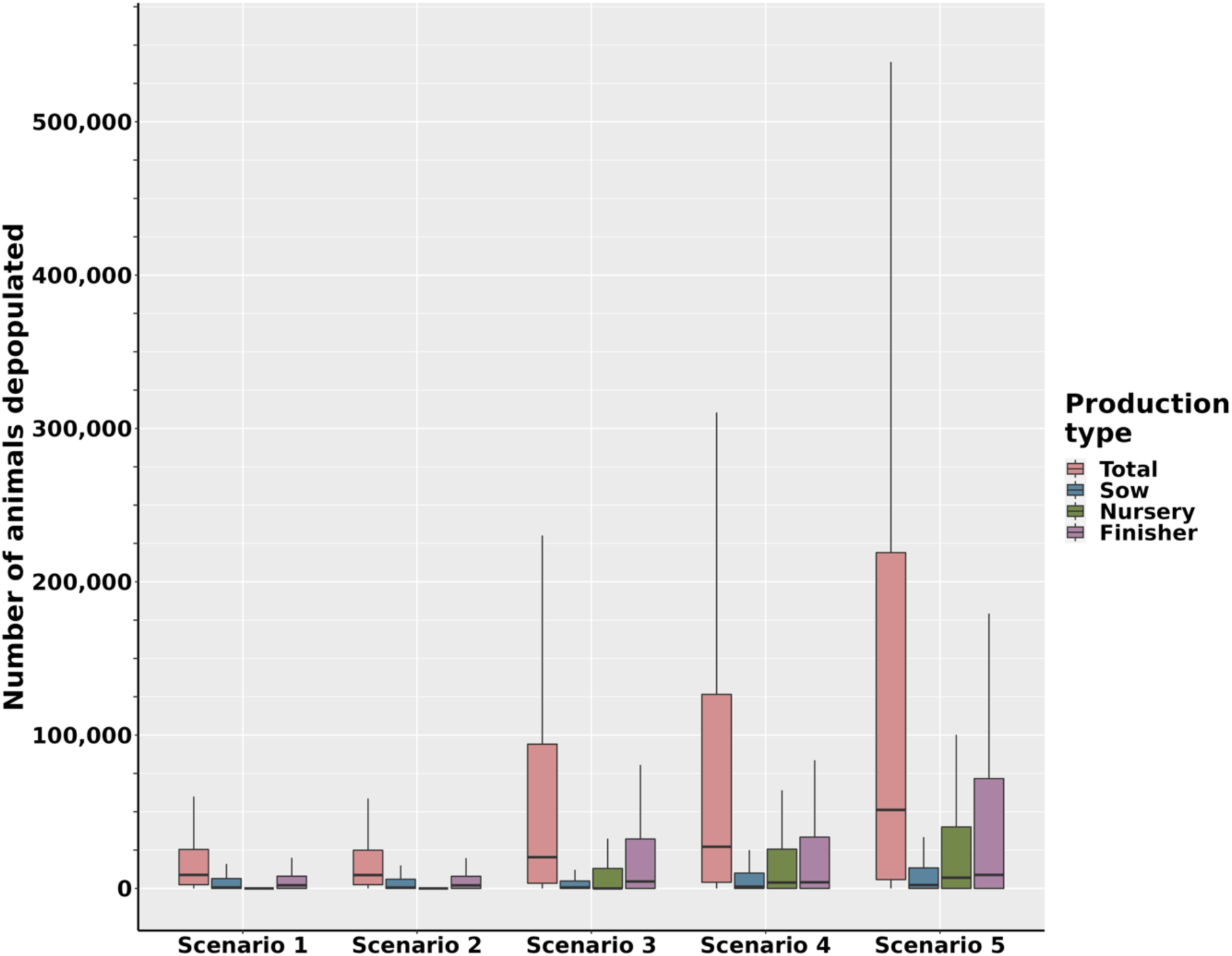
Box and whisker plots showing the number of animals depopulated under the five ASF control and eradication scenarios for each farm type at 60 days post seeded infection.

#### 3.5.2 Depopulation

Across all control scenarios, we investigated the number of depopulated swine considering farm capacity and production type (Figure 5). Scenario one showed a median of 8,800 (IQR: 2,448-25,440) depopulated swine, while scenario two demonstrated the lowest number of depopulated swine with a median of 8,700 (IQR: 2,448-24,929) (Figure 6). In scenarios three to five, the median number of swine depopulated increased substantially with a median of 20,380 (IQR: 3,350-94,100) swine depopulated in scenario three, a median of 27,177 (IQR: 4,000-126,560) swine depopulated in scenario four and a median of 50,208 (IQR: 5,280-217,283) swine depopulated in scenario five, representing the highest number of swine depopulated. Over 140 days of outbreak, the highest number of swine depopulated was under scenario four with an average of 699,887 swine (Supplementary Material Section E, Figure S21). This was followed by scenario three with 526,675 swine depopulated, scenario five with 488,531 swine depopulated, scenario one with 342,923 swine depopulated and scenario two with 332,398 swine depopulated. Additionally, as a result of enhanced surveillance, we observed the depopulation of swine in nursery farms in scenarios three, four, and five, which was not present in scenarios one and two due to the low effectiveness of surveillance assumed in this production type (Figure 5). Estimations on the financial compensation required under each scenario is given in Supplementary Material Section E, Tables S9 and S10.

**Figure 6.**
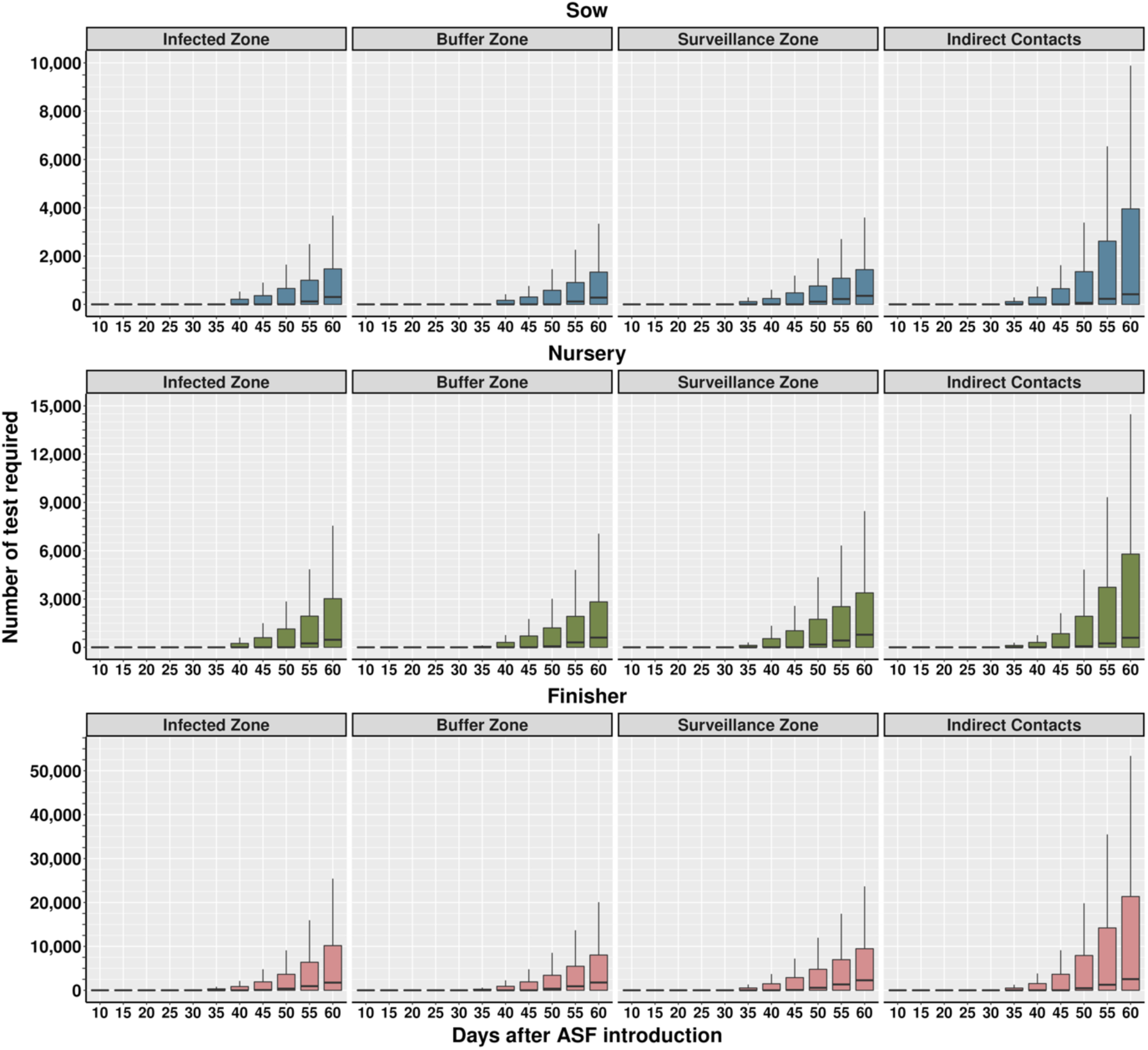
The cumulative number of diagnostic tests required for indirect contacts, infected zones, buffer zones and surveillance zones during control scenario five across 60 days of outbreaks, when samples are pooled in groups of five. Due to the delay in detection implemented in the model, testing was not required earlier than day 13 of the outbreak. Further details are available in section 2.5.

#### 3.5.3 Diagnostic testing

In section 3.5.1 we demonstrated that scenario five was the most effective in decreasing ASF secondary infection. Therefore, we selected only scenario five for further analysis. Over the first 60 days of the outbreak, the median cumulative number of tests required under scenario five, assuming samples were pooled into groups of five, was 11,974 (IQR: 0-68,337). Regardless of farm type, the testing of farms in the IZ required the most tests with a cumulative median of 2,609 (IQR: 0-14,409) (Figure 6) followed by the SZ, which had a cumulative median number of tests of 2,602 (IQR: 0-13,006). The number of tests required for farms in the BZ and indirect contact farms was slightly lower with a cumulative median of 2,473 (IQR: 0-11,626) and 2,513 (IQR: 0-28,074), respectively. Consistently, across the IZ, BZ, SZ and indirect contacts, more tests were required for finisher farms with the median number of tests ranging from 1,512 (IQR: 0-7,665) in the BZ to 1,759 (IQR: 0-19,173) for the indirect contacts. Nursery had lower numbers of the tests required, with median numbers of tests ranging from 364 (IQR: 0-5,188) for indirect contacts to 540 (IQR: 0-2,661) for the BZ. Sow farms had the lowest numbers of tests required, with a median number of tests ranging from 236 (IQR: 0-1,265) in the BZ to 296 (IQR: 0-3,517) for the indirect contacts.

By 140 days of the outbreak, a median of 357,789 (IQR: 24,429-840,126) diagnostic tests were required. Over this time frame, the largest volume of tests was for farms that were identified as indirect contacts, with a median of 115,729 (IQR: 2,798-255,948) tests required. More details regarding the number of tests required for 140 days of outbreaks are presented in Supplementary Material Section E, Figure S22.

#### 3.5.4 Movement permits required

Figure 7 presents the distribution of the required movement permits under scenario five. Daily permits were required for movements to and from farms within IZ and BZ, and movements to and from farms indirectly connected (via the transportation vehicles network) with ASF-positive farms. Over the first 60 days of outbreaks, a median of 1,808 (IQR: 0-9,737) permits was required. The majority of these permits were for movements of feed vehicles which had a median of 1,323 (IQR: 0-6,695) permits. The movement of swine vehicles had the second highest requirement for permits with a median of 240 (IQR: 0-1,641) permits, while the crew vehicles, market vehicles, and the movement of swine required substantially fewer permits with medians of 85 (IQR: 0-759), 43 (IQR: 0-274) and 58 (IQR: 0-291) permits, respectively.

**Figure 7.**
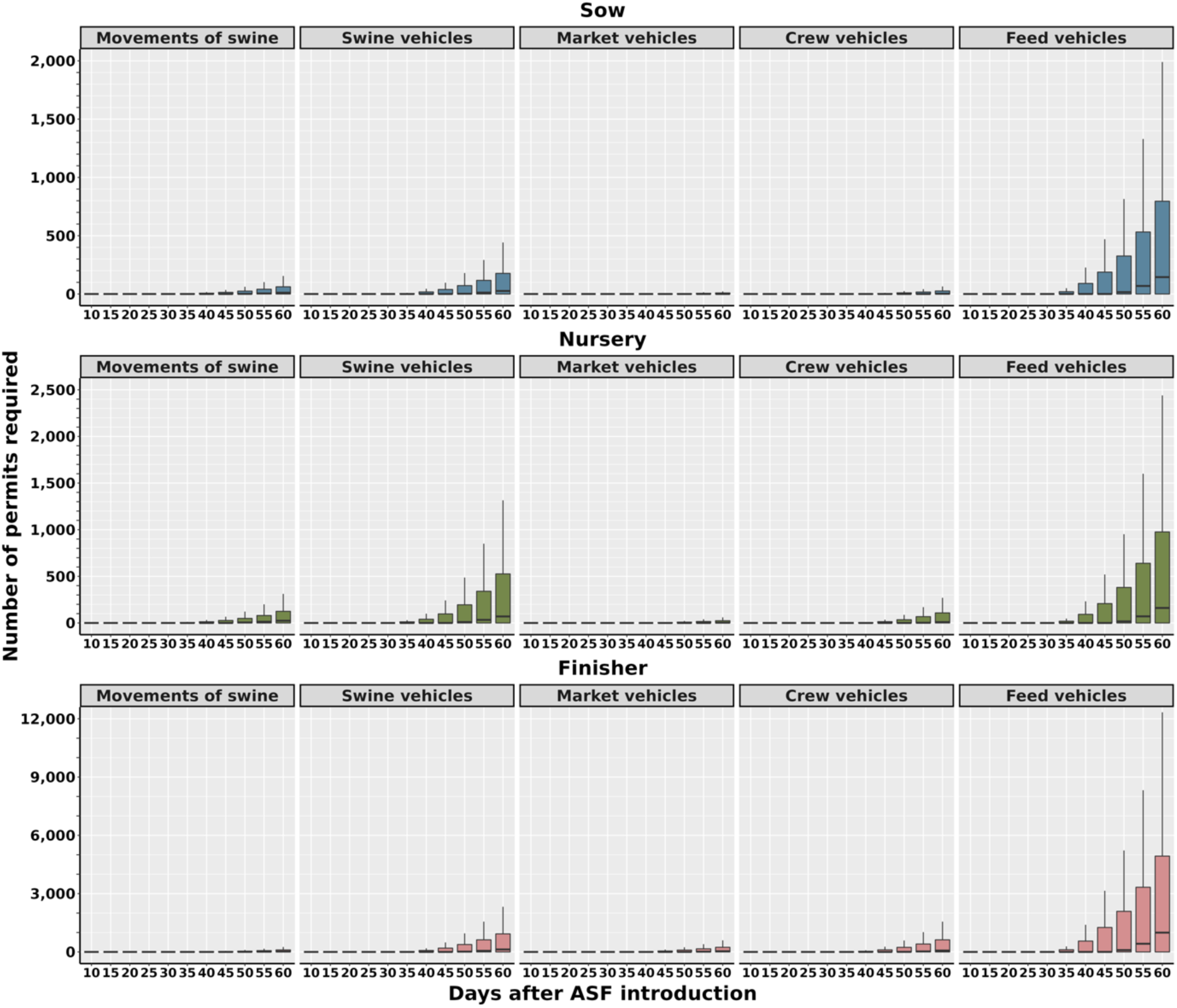
Median daily number of permits required for movements to or from infected zones, buffer zones and indirect contacts, during control scenario five across 60 days of outbreaks. Due to the delay in detection implemented in the model, the earliest date that permits were required was day 13 of the outbreak.

When considering the production type of the movement destination, we observed that movements into finisher farms consistently required more permits ranging from a median of 21 (IQR: 0-103) permits for movements of swine to a median of 994 (IQR: 0-4,933) permits required for movements of feed vehicles over 60 days. For nursery farms, the lowest number of permits was required for the movement of market vehicles with a median of 3 (IQR: 0-24) permits, while the movement of feed vehicles required the most with a median of 161 (IQR:0-976) permits. Lastly, movements to sow farms required the least amount of permits, ranging from a median of 0 (IQR: 0-9) permits for the movement of market vehicles to 145 (IQR: 0-796) permits for the movement of feed vehicles.

The number of permits required over the 140 days of simulated outbreaks was a median of 54,522 (IQR: 4,950-207,912) permits. The highest number of permits required was, once again, for the movement of feed vehicles, with a median of 37,741 (IQR: 3,554-73,339) permits, while the lowest number of permits was required for the movement of market vehicles, with a median of 1,518 (IQR: 154-2,967) permits. The full results of the permit requirements for 140 days of the outbreak are available in Supplementary Material Section E, Figures S23.

## 4. Discussion

In this study, we extended the previously developed farm-level stochastic transmission model *PigSpread* (Galvis, Corzo et al., 2022) to simulate ASF dissemination within the domestic swine population of the southeastern U.S.. This is the first published study to estimate the dissemination of ASF among a commercial swine population in the U.S. using real population and multiple contact networks, incorporating most of the known direct and indirect transmission routes for the virus (Hayes et al., 2021; Lee et al., 2021; Galli et al., 2022). Our data-driven modeling results elucidate the contribution of six transmission routes to ASF dissemination and evaluate the impact of control actions based on the Red Book (USDA, 2020), with a few key differences, while also estimating the numbers of depopulated animals, diagnostic tests, and movement permits required under the most effective combination of control actions.

### 4.1. ASF transmission

From our model simulations, we demonstrated that, on average, 72 farms could become infected in the first 60 days of dissemination, with finisher farms being the most affected, likely due, in part, to the larger number of finisher farms in the study population. We also identified between-farm movements of infected and exposed swine as the main routes of ASF transmission while also confirming the contribution of local spread and vehicle movement, particularly in the infection of finisher farms. Our results also showed long-distance spread between seeded and secondary infections as far as 474 km during the first 60 days and demonstrated ASF dissemination among farms not commercially related, which can be attributed to the spatial proximity between farms.

While the population structure of swine production and the pattern of between-farm contact networks is considerably different across countries, we observed similar epidemic trajectories in our model as seen in ASF transmission models from France and Australia (Bradhurst et al., 2021; Andraud et al., 2022). In the Australian model, 28 days of silent ASF dissemination within moderate to large commercial farms led to, on average, 4.3 to 6.3 infected farms (Bradhurst et al., 2021). This is particularly close to the eight infected farms that we observed over the same time frame. The incidence of infection observed in the French model was slightly lower with a median of 12 farms infected over 40 days (Andraud et al., 2022), whereas our model estimated 20 infections. These discrepancies could be attributed to differences in the population structure or model assumptions between the studies. Indeed, both studies (Bradhurst et al., 2021; Andraud et al., 2022) limited their local spread to 2 km compared to our less restrictive range of one km to 25 km, which coupled with the high density of farms in our study population (Galvis, Corzo, Prada et al., 2021; Jara et al., 2021), led to a more effective spread between farms. Similar ASF spread dynamics have been observed in outbreaks in Vietnam (Nguyen-Thi et al., 2021) and the Dominican Republic (Gonzales et al., 2021) where a substantial number of cases were observed in a short time frame.

The prominent route of ASF transmission in our study was the movement of infected and exposed swine, whose contribution to ASF dissemination varied from a minimum of 59.9% to a maximum of 86.7% over 140 days (Supplementary Material Section E, Figure S13). Recent mathematical simulation studies have demonstrated a similar dominance of between-farm swine movement in ASF dissemination (Halasa, Boklund et al., 2016; Andraud et al., 2019, 2022). An ASF transmission model in Denmark (Halasa, Bøtner et al., 2016), associated 99% of transmission with the movement of swine, while a French transmission model (Andraud et al., 2022) observed contributions between 68% and 85% for the movement of swine between farms. The contrast of the Danish model with the results observed in our model and the French model could be attributed to the structure of the models, the model assumptions and the pig population dynamics. Both our model and the French model, used detailed between-farm movement data including daily movements, while the Danish model approximated the daily frequency of movements from annual data, which may not accurately represent the contribution of swine movements. Additionally, in our study, each farm is under a swine production contract, only moving swine between farms of the same company which reduces the mixing of swine of different health statuses and subsequently lowers the probability of disease transmission (Passafaro et al., 2020; Galvis, Corzo, Prada et al., 2021).

We also observed a significant contribution of local spread and transportation vehicles to ASF dissemination, highlighting the role that such transmission routes may play in an ASF outbreak in the U.S. (Gao et al., 2021; Cheng and Ward, 2022; Galli et al., 2022). Local spread was particularly important in finisher farms where it contributed to nearly 20% of ASF dissemination, likely due to the high farm density in our study area. While previous literature has associated local spread with ASF transmission, there is no consensus on the maximum distance that ASF can spread via this route. To account for both short and long-distance transmission scenarios, such as those observed in ASF outbreaks in Africa, Asia and Europe (Okoth et al., 2013; Vergne et al., 2015; Boklund et al., 2020; Nguyen-Thi et al., 2021), we allowed our local spread to vary between one km and 25km. The contribution of swine vehicles was also important in finisher and nursery farms, most likely due to the vertical structure of the U.S. swine industry resulting in numerous swine vehicle movements to these production types. Similar to local spread, transportation vehicles have previously been recognized for their influence on ASF outbreaks in China and Korea (Yoo et al., 2021; Cheng and Ward, 2022), supporting the necessity to include such transmission routes in future ASF transmission models.

We also demonstrated long distance spread of ASF and dissemination between farms of unrelated production companies. Long distance spread in ASF has been observed before in outbreaks in the Republic of China (Akhmetzhanov et al., 2020) as well as a transmission model from France (Andraud et al., 2019, 2022). Interestingly our results diverge from those of the Danish transmission model (Halasa, Bøtner et al., 2016), where most secondary infections were at short distances (under ten km), which could be attributed to several variables including the geographical distribution of the study populations, model assumptions and the transmission route parameters (Boklund et al., 2009; Nigsch et al., 2013; Halasa, Boklund et al., 2016). In contrast to the long-distance spread of ASF, this study is the first description of between-company dissemination in ASF. As there are no recorded movements of swine or vehicles between companies, our results suggest that local spread was the most likely transmission route between companies. Indeed, 88.9% of farms in our study population were within 25 km (the maximum distance for ASF spread in the model) of a farm of another company. However, we cannot rule out the contribution of other transmission routes, such as rendering vehicles, which are managed by third-party companies and may serve multiple farms of different companies.

### 4.2. ASF control and elimination

When considering the effectiveness of the control actions included in this paper, we observed a reduction of ASF cases in all control scenarios compared to the no control scenario. With the addition of each control action, we demonstrated an increase in the efficacy of ASF control, with scenario five, which implements all the control actions, being the most successful, reducing secondary cases by 79% over 140 days of outbreaks. In particular, we observed substantial reductions in cases when adding control actions which enhanced ASF surveillance, such as contact tracing of direct contacts (contacts from swine movements) and indirect contacts (contacts from vehicle movements), and the implementation of control areas and surveillance zones which included frequent testing of farms. Similar reductions in cases have been observed in the French model (Andraud et al., 2022) and a model of ASF spread in the Rio Grande do Sul, Brazil (Machado et al., 2021), where the detection of ASF was increased by lowering the mortality threshold in farms. However, despite the increased effectiveness of scenario five, it was not capable of successfully eliminating the disease spread in all simulations by day 140, with only 29% of simulations being controlled. Thus, we demonstrated that with the current proposed control actions, an ASF outbreak duration in the southeastern U.S. will go beyond four months. This could be due to a number of factors including delayed detection of ASF, limited depopulation resources, and high connectivity of the swine industry in our study area.

To provide an estimate of the resources needed under scenario five, we estimated the quantity of swine depopulated, the number of diagnostic tests, and the volume of movement permits required Over the first 60 days of outbreaks, we observed a higher number of swine depopulated under scenario five, compared to the other scenarios, equating to a compensation cost of USD $3.8 million. However, when we consider 140 days of simulation, the compensation cost of depopulation in scenario five was lower than the cost estimated for scenarios three and four, with a maximum compensation cost of $79.2 million in scenario five compared to a maximum cost of $115.2 million in scenario four (Supplementary Material Section E, Table S11). Therefore, while the compensation cost of depopulation in scenario five is high during the first two months of the outbreaks, depopulation in the scenario becomes more cost effective as the outbreaks continue.

Over the simulated 140 days of the outbreak, we observed high numbers of diagnostic tests required for farms identified as indirect contacts of detected farms compared to the IZ, BZ or SZ, attributed to the high connectivity of the vehicle networks. Interestingly we noted a large number of tests required for the surveillance zones, which we attribute to the larger geographic area covered by the SZ compared to the CA, given the SZ surrounds the CA. Similarly, the density of farms in the region and their capacity will influence the number of diagnostic tests that would be required.

We also observed a large number of permits required for vehicles transporting feed, with a median of 37,741 permits required over 140 days, compared to the other movements of vehicles and swine. This substantial difference is due to the size of the feed vehicle network, which consisted of a median of 1,258 (IQR: 257-1,877) farms visited per day (Supplementary Material Section F, Table S9). Despite the decrease in cases associated with scenario five, which will reduce the resources required, the compensation cost of depopulation and quantities of movement permits are significant. By considering permits that cover movement for a 30-day time period for select movement types within the model, such as the movement of feed vehicles, between the same farms, we may reduce the resources needed. Nevertheless, the estimation of resources is pivotal to gauge the feasibility of strategies and help animal health officials anticipate the workload needed in the event of an ASF introduction.

## 5. Limitations and further remarks

While our ASF stochastic transmission model is the first to model ASF between-farm dissemination in the southeastern U.S. using real population and multiple contact networks (swine and vehicles), the lack of ASF outbreak data limited our ability to calibrate transmission parameters. As a novel alternative, we used the first waves of the 2013 PEDV epidemic in the study region (Machado et al., 2019), to reflect the dynamics of a novel virus spreading in a completely susceptible population. Yet, we cannot be certain that ASF dissemination would follow the same transmission dynamics as PEDV, as each virus has a different fitness and transmission capability (Pensaert and de Bouck, 1978; Alonso et al., 2014). An example is differences in transmission rates, which are similar but not equal, as the basic reproduction number *R*_0_ of PEDV has been estimated to be 1.87 (95% CI: 1.52-2.34) (Greer et al., 2017) while ASF *R*_0_ in Vietnam has been estimated to be at 1.49 (95%CI: 1.05–2.21) (Mai et al., 2022). Despite the limitations of using PEDV data to calibrate the ASF model discussed above, we believe that they do not outweigh the benefit of using real outbreak data from our study population to capture the parameters of epidemic disease spread within this region. Similarly, we chose to use PEDV over other swine diseases such as porcine reproductive and respiratory syndrome virus (PRRSV), as we do not have epidemic outbreak data for PRRSV in our study region.

Another limiting factor of our model is the paucity of data regarding additional transmission routes, such as the movements of rendering vehicles, the distribution of independent backyard producers, and the density of feral swine (Yoo et al., 2021; Galli et al., 2022; Pepin et al., 2022). As the access to such data becomes available in the future, subsequent iterations of this model should take into consideration the influence that these transmission routes and population dynamics may have on ASF dissemination (Neumann et al., 2021; Cheng and Ward, 2022). In particular, the introduction of ASF into feral swine populations is greatly important, as it has the potential for spillover into domestic populations (Hayes et al., 2021) due to their wide geographic range (Fodor et al., 2015; Iglesias et al., 2016; Boklund et al., 2020). It should also be noted that without the relevant data from the other U.S. states, the results of this model cannot be generalized outside of the southeastern U.S..

Limited data also may have impacted the transmission routes already considered in the model, as we were unable to collect transportation vehicle movement data from 24% of farms in the study. Additionally, farms were identified as a node in the vehicle networks if a vehicle was stationary within 1.5 km of the farm, however, due to the high density of the farms in our study, a vehicle could have been in 1.5 km of multiple farms at the same time, over-estimating the number of vehicle contacts (Galvis, Corzo et al., 2022). In the future, we expect to improve the reconstruction of the vehicle movement networks, to include these missing vehicle movements and incorporate the movement of additional third-party vehicles, such as rendering vehicles and maintenance crews which are documented as having an important role in the dissemination of ASF in Asia (Neumann et al., 2021; Cheng and Ward, 2022). There is also a myriad of factors such as presence of other pathogens within the population which could influence the impact of an ASF outbreak, however, without comprehensive data or evidence we are not able to simulate this in our model.In future iterations of this model, we will include biosecurity practices present on the farms to account for the direct impact they can have on the pathogens’ forces of infection, such as the presence of on-farm cleaning and disinfection stations which has been shown to reduce the risk of pathogen introduction (Bellini et al., 2016; De Lorenzi et al., 2020; Juszkiewicz et al., 2020; Jiang et al., 2021) and the use of open-sided barns which increases the chance of direct nose-to-nose contact between feral swine and domestic pigs (Yoo et al., 2021; N. T. Mai et al., 2022). It may also be pertinent to model the level of compliance regarding the control actions implemented, which has been associated with ASF persistence in Sardinia (Cappai et al., 2018), and incorporate varying effectiveness of cleaning and disinfection practices, which our current model assumes to be 100%. Additionally, outbreaks of ASF within Europe have identified the spread of less virulent strains of ASF, which has been suggested to lengthen the delay in detection of the initial case, increasing the duration of silent dissemination. Currently the results of this model only apply to an outbreak of highly pathogenic ASF.

Along with the above additions to the model, we also plan to explore the implementation of quarantine to premises in the CA and those identified as contacts, as well as removing the depopulation of direct contacts, in an effort to model realistic scenarios included in the USDA’s current ASF response plan. Thus, future modeling efforts will unlock our capacity to understand ASF propagation at multiple levels and test the effectiveness of disease control strategies under different conditions.

## 6. Conclusion

In this study we have demonstrated that an introduction of ASF into the southeastern U.S. could result in 72 infected farms in the first two months of an outbreak, with the potential for between-company dissemination. Transmission of ASF via between-farm movements of swine was identified as the predominant route of ASF dissemination, however, local spread and the movement of vehicles also contributed significantly to ASF spread, resulting in long-distance spread of ASF as far as 474km away from the index case. However, with the implementation of the control actions similar to what is described in the national response plan, secondary cases could be reduced by 79% by day 140. Despite this reduction in cases, the above control strategy was not able to stamp-out the epidemic in all simulations by day 140 and required a substantial amount of diagnostic testing, movement permits, and depopulation resources. Nevertheless, the critical evaluation of ASF response and control activities provided, along with estimations of the resources needed under the most effective scenario, can guide further research and preparations to ensure we can effectively limit a future outbreak.

## Acknowledgments

The authors would like to acknowledge participating companies and veterinarians for their insightful comments and engaging discussions. We also thank members of the United States Department of Agriculture Animal and Plant Health Inspection Service (USDA-APHIS) of the Center for Epidemiology and Animal Health (CEAH) and National Preparedness and Incident Coordination Center (NPIC).

## Conflict of interest

All authors confirm that there are no conflicts of interest to declare

## Ethical statement

The authors confirm the ethical policies of the journal, as noted on the journal’s author guidelines page. Since this work did not involve animal sampling nor questionnaire data collection by the researchers, there was no need for ethics permits.

## Data Availability Statement

The data that support the findings of this study are not publicly available and are protected by confidential agreements, therefore, are not available. A preprint has previously been published Abagael L. Sykes et.al 2022 and code examples can be found at https://github.com/machado-lab/PigSpread-ASF.

## Funding

This project is funded by USDA’s Animal and Plant Health Inspection Service through the National Animal Disease Preparedness and Response Program via a cooperative agreement between the Animal and Plant Health Inspection Service (APHIS) Veterinary Services (VS) and North Carolina State University, USDA-APHIS Award: AP22VSSP0000C004. The Morrison Swine Health Monitoring Project is a Swine Health Information Center (SHIC) funded project.

## Footnotes

1 Sow farms included: farrow, farrow-to-wean and farrow-to-feeder farms.

2 Finisher farms included: wean-to-feeder, wean-to-finish, feeder-to-finish farms.

3 Gilt development and isolation: farms that could either be a finisher or sow farm depending upon the production company.

4 While current USDA plans for a movement standstill will only cover live swine movements and related vehicles, we have modeled stricter controls.

5 We assumed that all direct contacts were considered high risk and therefore were depopulated, however an automatic depopulation of direct contacts without a known ASF disease status is not included in the USDA’s current ASF response. This current plan would also quarantine direct and indirect contacts

## Notes

### Competing Interest Statement

The authors have declared no competing interest.

### Summary of Updates

Updated results

## References

Abagael L. Sykes, Jason A. Galvis, Kathleen C. O’Hara, Cesar Corzo, Gustavo Machado bioRxiv 2022.09.04.506538; doi: https://doi.org/10.1101/2022.09.04.506538.

Adedeji, A.J., R.B. Atai, H.E. Gyang, P. Gambo, M.A. Habib, R. Weka, V.B. Muwanika, C. Masembe, and P.D. Luka, 2022: Live pig markets are hotspots for spread of African swine fever virus in Nigeria. Transboundary and Emerging Diseases n/a, DOI: 10.1111/tbed.14483.

Akhmetzhanov, A.R., S. Jung, H. Lee, N. Linton, Y. Yang, B. Yuan, and H. Nishiura, 2020: Reconstruction and analysis of the transmission network of African swine fever in People’s Republic of China, August 2018–September 2019 (preprint). Ecology.

Alonso, C., D.P. Goede, R.B. Morrison, P.R. Davies, A. Rovira, D.G. Marthaler, and M. Torremorell, 2014: Evidence of infectivity of airborne porcine epidemic diarrhea virus and detection of airborne viral RNA at long distances from infected herds. Veterinary Research 45, 73, DOI: 10.1186/s13567-014-0073-z.

Andraud, M., T. Halasa, A. Boklund, and N. Rose, 2019: Threat to the French Swine Industry of African Swine Fever: Surveillance, Spread, and Control Perspectives. Frontiers in Veterinary Science 6, 248, DOI: 10.3389/fvets.2019.00248.

Andraud, M., P. Hammami, B. H. Hayes, J. A. Galvis, T. Vergne, G. Machado, and N. Rose, 2022: Modelling African swine fever virus spread in pigs using time-respective network data: Scientific support for decision makers. Transbounding Emerging Distbed.14550, DOI: 10.1111/tbed.14550.

Andraud, M., P. Hammami, B.H. Hayes, J.A. Galvis, G. Machado, and N. Rose, 2021: Modelling African swine fever virus spread in pigs using time-respective network data: scientific support for decision-makers. arxiv.

Bellini, S., D. Rutili, and V. Guberti, 2016: Preventive measures aimed at minimizing the risk of African swine fever virus spread in pig farming systems. Acta Veterinaria Scandinavica 58, 82, DOI: 10.1186/s13028-016-0264-x.

Boklund, A., S. Dhollander, T. Chesnoiu Vasile, J.C. Abrahantes, A. Bøtner, A. Gogin, L.C. Gonzalez Villeta, C. Gortázar, S.J. More, A. Papanikolaou, H. Roberts, A. Stegeman, K. Ståhl, H.H. Thulke, A. Viltrop, Y. Van der Stede, and S. Mortensen, 2020: Risk factors for African swine fever incursion in Romanian domestic farms during 2019. Sci Rep 10, 10215, DOI: 10.1038/s41598-020-66381-3.

Boklund, A., N. Toft, L. Alban, and Å. Uttenthal, 2009: Comparing the epidemiological and economic effects of control strategies against classical swine fever in Denmark. Preventive Veterinary Medicine 90, 180–193, DOI: 10.1016/j.prevetmed.2009.04.008.

Bradhurst, R., G. Garner, S. Roche, R. Iglesias, N. Kung, B. Robinson, S. Willis, M. Cozens, K. Richards, B. Cowled, M. Oberin, C. Tharle, S. Firestone, and M. Stevenson, 2021: Modelling the spread and control of African swine fever in domestic and feral pigs (Technical report). Centre of Excellence for Biosecurity Risk Analysis.

Cannon, R.M., 2001: Sense and sensitivity — designing surveys based on an imperfect test. Preventive Veterinary Medicine 49, 141–163, DOI: 10.1016/S0167-5877(01)00184-2.

Carlson, J., M. Fischer, L. Zani, M. Eschbaumer, W. Fuchs, T. Mettenleiter, M. Beer, and S. Blome, 2020: Stability of African Swine Fever Virus in Soil and Options to Mitigate the Potential Transmission Risk. Pathogens 9, 977, DOI: 10.3390/pathogens9110977.

Chenais, E., K. Depner, V. Guberti, K. Dietze, A. Viltrop, and K. Ståhl, 2019: Epidemiological considerations on African swine fever in Europe 2014–2018. Porcine Health Management 5, 6, DOI: 10.1186/s40813-018-0109-2.

Cheng, J., and M.P. Ward, 2022: Risk factors for the spread of African Swine Fever in China: A systematic review of Chinese-language literature. Transboundary and Emerging Diseases1–10, DOI: 10.1111/tbed.14573.

Chuchard, P., D. Prathumwan, K. Trachoo, W. Maiaugree, and I. Chaiya, 2022: The SLI-SC Mathematical Model of African Swine Fever Transmission among Swine Farms: The Effect of Contaminated Human Vector. Axioms 11, 329, DOI: 10.3390/axioms11070329.

Costard, S., B. Wieland, W. de Glanville, F. Jori, R. Rowlands, W. Vosloo, F. Roger, D.U. Pfeiffer, and L.K. Dixon, 2009: African swine fever: how can global spread be prevented? Philosophical Transactions of the Royal Society B: Biological Sciences 364, 2683–2696, DOI: 10.1098/rstb.2009.0098.

de la Torre, A., J. Bosch, J.M. Sánchez-Vizcaíno, S. Ito, C. Muñoz, I. Iglesias, and M. Martínez-Avilés, 2022: African Swine Fever Survey in a European Context. Pathogens 11, 137, DOI: 10.3390/pathogens11020137.

De Lorenzi, G., L. Borella, G.L. Alborali, J. Prodanov-Radulović, M. Štukelj, and S. Bellini, 2020: African swine fever: A review of cleaning and disinfection procedures in commercial pig holdings. Research in Veterinary Science 132, 262–267, DOI: 10.1016/j.rvsc.2020.06.009.

Efsa, D. Desmecht, G. Gerbier, C. Gortázar Schmidt, V. Grigaliuniene, G. Helyes, M. Kantere, D. Korytarova, A. Linden, A. Miteva, I. Neghirla, E. Olsevskis, S. Ostojic, T. Petit, C. Staubach, H.-H. Thulke, A. Viltrop, W. Richard, G. Wozniakowski, J.A. Cortiñas, A. Broglia, S. Dhollander, E. Lima, A. Papanikolaou, Y. Van der Stede, and K. Ståhl, 2021: Epidemiological analysis of African swine fever in the European Union (September 2019 to August 2020). EFSA Journal 19, e06572, DOI: 10.2903/j.efsa.2021.6572.

EFSA Panel on Animal Health and Welfare (AHAW), 2010: Scientific Opinion on African Swine Fever. EFSA Journal 8, 1556, DOI: 10.2903/j.efsa.2010.1556.

EFSA Panel on Animal Health and Welfare (AHAW), 2014: Scientific Opinion on African swine fever. EFSA Journal 12, 3628, DOI: 10.2903/j.efsa.2014.3628.

Ezanno, P., S. Picault, S. Bareille, G. Beaunée, G.J. Boender, E.A. Dankwa, F. Deslandes, C.A. Donnelly, T.J. Hagenaars, S. Hayes, F. Jori, S. Lambert, M. Mancini, F. Munoz, D.R.J. Pleydell, R.N. Thompson, E. Vergu, M. Vignes, and T. Vergne, 2022: The African swine fever modelling challenge: Model comparison and lessons learnt. Epidemics 40, 100615, DOI: 10.1016/j.epidem.2022.100615.

Ferdousi, T., S.A. Moon, A. Self, and C. Scoglio, 2019: Generation of swine movement network and analysis of efficient mitigation strategies for African swine fever virus. PLOS ONE 14, e0225785, DOI: 10.1371/journal.pone.0225785.

Fodor, J.T., F. Jánoska, and A. Farkas, 2015: The comparative analysis of the habitat use of wild boar in different Romanian habitats (partial results). Proceedings of the Biennial International Symposium. Forest and sustainable development, Brașov, Romania, 24-25th October 2014365–370.

Franzo, G., G. Barbierato, P. Pesente, M. Legnardi, C.M. Tucciarone, G. Sandri, and M. Drigo, 2021: Porcine Reproductive and Respiratory Syndrome (PRRS) Epidemiology in an Integrated Pig Company of Northern Italy: A Multilevel Threat Requiring Multilevel Interventions. Viruses 13, 2510, DOI: 10.3390/v13122510.

Gallardo, C., A. Soler, R. Nieto, C. Cano, V. Pelayo, M.A. Sánchez, G. Pridotkas, J. Fernandez-Pinero, V. Briones, and M. Arias, 2017: Experimental Infection of Domestic Pigs with African Swine Fever Virus Lithuania 2014 Genotype II Field Isolate. Transboundary and Emerging Diseases 64, 300– 304, DOI: 10.1111/tbed.12346.

Galli, F., B. Friker, A. Bearth, and S. Dürr, 2022: Direct and indirect pathways for the spread of African swine fever and other porcine infectious diseases: An application of the Mental Models Approach. Transboundary and Emerging Diseases n/a, DOI: 10.1111/tbed.14605.

Galvis, J.A., C. Corzo, and G. Machado, 2021: Modelling porcine reproductive and respiratory syndrome virus dynamics to quantify the contribution of multiple modes of transmission: between-farm animal and vehicle movements, farm-to-farm proximity, feed ingredients, and re-break. bioRxiv2021.07.26.453902, DOI: 10.1101/2021.07.26.453902.

Galvis, J.A., C.A. Corzo, and G. Machado, 2022: Modelling and assessing additional transmission routes for porcine reproductive and respiratory syndrome virus: Vehicle movements and feed ingredients. Transbounding Emerging Distbed.14488, DOI: 10.1111/tbed.14488.

Galvis, J.A., C.A. Corzo, J.M. Prada, and G. Machado, 2021: Modelling the transmission and vaccination strategy for porcine reproductive and respiratory syndrome virus. Transbound Emerg DisDOI: 10.1111/tbed.14007.

Galvis, J.A., C.M. Jones, J.M. Prada, C.A. Corzo, and G. Machado, 2022: The between-farm transmission dynamics of porcine epidemic diarrhoea virus: A short-term forecast modelling comparison and the effectiveness of control strategies. Transboundary and Emerging Diseases 69, 396–412, DOI: 10.1111/tbed.13997.

Gao, L., X. Sun, H. Yang, Q. Xu, J. Li, J. Kang, P. Liu, Y. Zhang, Y. Wang, and B. Huang, 2021: Epidemic situation and control measures of African Swine Fever Outbreaks in China 2018–2020. Transbound. Emerg. Dis. 68, 2676–2686, DOI: 10.1111/tbed.13968.

Gebhardt, J.T., S.S. Dritz, C.G. Elijah, C.K. Jones, C.B. Paulk, and J.C. Woodworth, 2021: Sampling and detection of African swine fever virus within a feed manufacturing and swine production system. Transboundary and Emerging DiseasesDOI: 10.1111/tbed.14335.

Gonzales, W., C. Moreno, U. Duran, N. Henao, M. Bencosme, P. Lora, R. Reyes, R. Núñez, A. De Gracia, and A.M. Perez, 2021: African swine fever in the Dominican Republic. Transboundary and Emerging Diseases 68, 3018–3019, DOI: 10.1111/tbed.14341.

Greer, A.L., K. Spence, and E. Gardner, 2017: Understanding the early dynamics of the 2014 porcine epidemic diarrhea virus (PEDV) outbreak in Ontario using the incidence decay and exponential adjustment (IDEA) model. BMC Veterinary Research 13, 8, DOI: 10.1186/s12917-016-0922-2.

Halasa, T., A. Boklund, A. Bøtner, N. Toft, and H.-H. Thulke, 2016: Simulation of Spread of African Swine Fever, Including the Effects of Residues from Dead Animals. Frontiers in Veterinary Science 3.

Halasa, T., A. Bøtner, S. Mortensen, H. Christensen, N. Toft, and A. Boklund, 2016: Simulating the epidemiological and economic effects of an African swine fever epidemic in industrialized swine populations. Veterinary Microbiology 193, 7–16, DOI: 10.1016/j.vetmic.2016.08.004.

Halasa, T., M.P. Ward, and A. Boklund, 2020: The impact of changing farm structure on foot-and-mouth disease spread and control: A simulation study. Transboundary and Emerging Diseases 67, 1633–1644, DOI: 10.1111/tbed.13500.

Hartig, F., J.M. Calabrese, B. Reineking, T. Wiegand, and A. Huth, 2011: Statistical inference for stochastic simulation models – theory and application. Ecology Letters 14, 816–827, DOI: 10.1111/j.1461-0248.2011.01640.x.

Hayes, B.H., M. Andraud, L.G. Salazar, N. Rose, and T. Vergne, 2021: Mechanistic modelling of African swine fever: A systematic review. Preventive Veterinary Medicine 191, 105358, DOI: 10.1016/j.prevetmed.2021.105358.

Hoar, B., J. Angelos, A. Arens, and J. Humphrey, 2015: Production Cycle of Swine. Western Institute for Food Safety and Security at University of California Davis and the Food and Drug Administration.

Hu, B., J.L. Gonzales, and S. Gubbins, 2017: Bayesian inference of epidemiological parameters from transmission experiments. Sci Rep 7, 16774, DOI: 10.1038/s41598-017-17174-8.

Iglesias, I., M.J. Muñoz, F. Montes, A. Perez, A. Gogin, D. Kolbasov, and A. de la Torre, 2016: Reproductive Ratio for the Local Spread of African Swine Fever in Wild Boars in the Russian Federation. Transboundary and Emerging Diseases 63, e237–e245, DOI: 10.1111/tbed.12337.

Iscaro, C., A. Dondo, L. Ruocco, L. Masoero, M. Giammarioli, S. Zoppi, V. Guberti, and F. Feliziani, 2022: January 2022: Index case of new African Swine Fever incursion in mainland Italy. Transboundary and Emerging DiseasesDOI: 10.1111/tbed.14584.

Jara, M., D.A. Rasmussen, C.A. Corzo, and G. Machado, 2021: Porcine reproductive and respiratory syndrome virus dissemination across pig production systems in the United States. Transboundary and Emerging Diseases 68, 667–683, DOI: 10.1111/tbed.13728.

Jiang, C., Y. Sun, F. Zhang, X. Ai, X. Feng, W. Hu, X. Zhang, D. Zhao, Z. Bu, and X. He, 2021: Viricidal activity of several disinfectants against African swine fever virus. Journal of Integrative Agriculture 20, 3084–3088, DOI: 10.1016/S2095-3119(21)63631-6.

Juszkiewicz, M., M. Walczak, N. Mazur-Panasiuk, and G. Woźniakowski, 2020: Effectiveness of Chemical Compounds Used against African Swine Fever Virus in Commercial Available Disinfectants. Pathogens 9, 878, DOI: 10.3390/pathogens9110878.

Kim, Y.-J., B. Park, and H.-E. Kang, 2021: Control measures to African swine fever outbreak: Active response in South Korea, preparation for the future, and cooperation. J Vet Sci 22, e13, DOI: 10.4142/jvs.2021.22.e13.

Korennoy, F.I., V.M. Gulenkin, J.B. Malone, C.N. Mores, S.A. Dudnikov, and M.A. Stevenson, 2014: Spatio-temporal modeling of the African swine fever epidemic in the Russian Federation, 2007– 2012. Spatial and Spatio-temporal Epidemiology 11, 135–141, DOI: 10.1016/j.sste.2014.04.002.

Lee, H.S., K.K. Thakur, V.N. Bui, T.L. Pham, A.N. Bui, T.D. Dao, V.T. Thanh, and B. Wieland, 2021: A stochastic simulation model of African swine fever transmission in domestic pig farms in the Red River Delta region in Vietnam. Transbound Emerg Dis 68, 1384–1391, DOI: 10.1111/tbed.13802.

Li, X., Z. Hu, M. Fan, W. Wu, W. Gao, L. Bian, W. Liu, X. Tian, X. Jiang, and Z.J. Yan, 2022: Evidence of aerosol transmission of African swine fever virus in piggeries under field conditions: a case report (preprint). In Review.

Li, Y., M. Salman, C. Shen, H. Yang, Y. Wang, Z. Jiang, J. Edwards, and B. Huang, 2020: African Swine Fever in a commercial pig farm: Outbreak investigation and an approach for identifying the source of infection. Transboundary and Emerging Diseases 67, 2564–2578, DOI: 10.1111/tbed.13603.

Machado, G., T.S. Farthing, M. Andraud, F.P.N. Lopes, and C. Lanzas, 2021: Modelling the role of mortality-based response triggers on the effectiveness of African swine fever control strategies. Transboundary and Emerging Diseases n/a, DOI: 10.1111/tbed.14334.

Machado, G., C. Vilalta, M. Recamonde-Mendoza, C. Corzo, M. Torremorell, A. Perez, and K. VanderWaal, 2019: Identifying outbreaks of Porcine Epidemic Diarrhea virus through animal movements and spatial neighborhoods. Sci Rep 9, 457, DOI: 10.1038/s41598-018-36934-8.

Mai, T.N., S. Sekiguchi, T.M.L. Huynh, T.B.P. Cao, V.P. Le, V.H. Dong, V.A. Vu, and A. Wiratsudakul, 2022: Dynamic Models of Within-Herd Transmission and Recommendation for Vaccination Coverage Requirement in the Case of African Swine Fever in Vietnam. Vet Sci 9, 292, DOI: 10.3390/vetsci9060292.

Malladi, S., A. Ssematimba, P.J. Bonney, K.M.S. Charles, T. Boyer, T. Goldsmith, E. Walz, C. Cardona, and M.R. Culhane, 2022: Predicting the Time to Detect Moderately Virulent African Swine Fever Virus in Finisher Swine Herds Using a Stochastic Disease Transmission Model. BMC Veterinary Research (PREPRINT)DOI: 10.21203/rs.3.rs-716595/v1.

Mason-D’Croz, D., J.R. Bogard, M. Herrero, S. Robinson, T.B. Sulser, K. Wiebe, D. Willenbockel, and H.C.J. Godfray, 2020: Modelling the global economic consequences of a major African swine fever outbreak in China. Nat Food 1, 221–228, DOI: 10.1038/s43016-020-0057-2.

Mazur-Panasiuk, N., J. Żmudzki, and G. Woźniakowski, 2019: African Swine Fever Virus – Persistence in Different Environmental Conditions and the Possibility of its Indirect Transmission. J Vet Res 63, 303–310, DOI: 10.2478/jvetres-2019-0058.

Mighell, E., and M.P. Ward, 2021: African Swine Fever spread across Asia, 2018–2019. Transboundary and Emerging Diseases 68, 2722–2732, DOI: 10.1111/tbed.14039.

Miller, R.S., S.J. Sweeney, C. Slootmaker, D.A. Grear, P.A. Di Salvo, D. Kiser, and S.A. Shwiff, 2017: Cross-species transmission potential between wild pigs, livestock, poultry, wildlife, and humans: implications for disease risk management in North America. Sci Rep 7, 7821, DOI: 10.1038/s41598-017-07336-z.

Minter, A., and R. Retkute, 2019: Approximate Bayesian Computation for infectious disease modelling. Epidemics 29, 100368, DOI: 10.1016/j.epidem.2019.100368. MSHMP, 2022:

Mur, L., J.M. Sánchez-Vizcaíno, E. Fernández-Carrión, C. Jurado, S. Rolesu, F. Feliziani, A. Laddomada, and B. Martínez-López, 2018: Understanding African Swine Fever infection dynamics in Sardinia using a spatially explicit transmission model in domestic pig farms. Transboundary and Emerging Diseases 65, 123–134, DOI: 10.1111/tbed.12636.

National Pork Board, 2021: Pork Checkoff, Life Cycle of a Market Pig [Online] Available at https://porkcheckoff.org/pork-branding/facts-statistics/life-cycle-of-a-market-pig/ (accessed August 25, 2022).

Neumann, E., W. Hall, J. Dahl, D. Hamilton, and A. Kurian, 2021: Is transportation a risk factor for African swine fever transmission in Australia: a review. Aust Vet J 99, 459–468, DOI: 10.1111/avj.13106.

Nguyen-Thi, T., L. Pham-Thi-Ngoc, Q. Nguyen-Ngoc, S. Dang-Xuan, H.S. Lee, H. Nguyen-Viet, P. Padungtod, T. Nguyen-Thu, T. Nguyen-Thi, T. Tran-Cong, and K.M. Rich, 2021: An Assessment of the Economic Impacts of the 2019 African Swine Fever Outbreaks in Vietnam. Front Vet Sci 8, 686038, DOI: 10.3389/fvets.2021.686038.

Niemi, J.K., 2020: Impacts of African Swine Fever on Pigmeat Markets in Europe. Front Vet Sci 7, 634, DOI: 10.3389/fvets.2020.00634.

Nigsch, A., S. Costard, B.A. Jones, D.U. Pfeiffer, and B. Wieland, 2013: Stochastic spatio-temporal modelling of African swine fever spread in the European Union during the high risk period. Preventive Veterinary Medicine 108, 262–275, DOI: 10.1016/j.prevetmed.2012.11.003.

OIE, 2021 (1. November): ProMED, African Swine Fever - Americas (07): Dominican Republic, Haiti, Spread, OIE [Online] Available at https://promedmail.org/promed-post/?id=8699382 (accessed January 5, 2022).

Okoth, E., C. Gallardo, J.M. Macharia, A. Omore, V. Pelayo, D.W. Bulimo, M. Arias, P. Kitala, K. Baboon, I. Lekolol, D. Mijele, and R.P. Bishop, 2013: Comparison of African swine fever virus prevalence and risk in two contrasting pig-farming systems in South-west and Central Kenya. Preventive Veterinary Medicine 110, 198–205, DOI: 10.1016/j.prevetmed.2012.11.012.

Olesen, A.S., L. Lohse, A. Boklund, T. Halasa, G.J. Belsham, T.B. Rasmussen, and A. Bøtner, 2018: Short time window for transmissibility of African swine fever virus from a contaminated environment. Transboundary and Emerging Diseases 65, 1024–1032, DOI: 10.1111/tbed.12837.

Olesen, A.S., L. Lohse, A. Boklund, T. Halasa, C. Gallardo, Z. Pejsak, G.J. Belsham, T.B. Rasmussen, and A. Bøtner, 2017: Transmission of African swine fever virus from infected pigs by direct contact and aerosol routes. Veterinary Microbiology 211, 92–102, DOI: 10.1016/j.vetmic.2017.10.004.

Passafaro, T.L., A.F.A. Fernandes, B.D. Valente, N.H. Williams, and G.J.M. Rosa, 2020: Network analysis of swine movements in a multi-site pig production system in Iowa, USA. Preventive Veterinary Medicine 174, 104856, DOI: 10.1016/j.prevetmed.2019.104856.

Penrith, M.L., and F.M. Kivaria, 2022: One hundred years of African swine fever in Africa: Where have we been, where are we now, where are we going? Transbounding Emerging Distbed.14466, DOI: 10.1111/tbed.14466.

Pensaert, M.B., and P. de Bouck, 1978: A new coronavirus-like particle associated with diarrhea in swine. Arch Virol 58, 243–247, DOI: 10.1007/BF01317606.

Pepin, K.M., V.R. Brown, A. Yang, J.C. Beasley, R. Boughton, K.C. VerCauteren, R.S. Miller, and S.N. Bevins, 2022: Optimizing response to an introduction of African swine fever in wild pigs. Transboundary and Emerging DiseasesDOI: 10.1111/tbed.14668.

Pepin, K.M., A. Golnar, and T. Podgórski, 2021: Social structure defines spatial transmission of African swine fever in wild boar. Journal of The Royal Society Interface 18, 20200761, DOI: 10.1098/rsif.2020.0761.

PIC, 2020: Breed gilts at the right time to optimize performance.

Podgórski, T., and K. Śmietanka, 2018: Do wild boar movements drive the spread of African Swine Fever? Transbound Emerg Dis 65, 1588–1596, DOI: 10.1111/tbed.12910.

Sanson, R.L., 1994: The epidemiology of foot-and-mouth disease: Implications for New Zealand. New Zealand Veterinary Journal 42, 41–53, DOI: 10.1080/00480169.1994.35785.

Schulz, K., F.J. Conraths, S. Blome, C. Staubach, and C. Sauter-Louis, 2019: African Swine Fever: Fast and Furious or Slow and Steady? Viruses 11, 866, DOI: 10.3390/v11090866.

Sisson, S.A., Y. Fan, and M.M. Tanaka, 2007: Sequential Monte Carlo without likelihoods. Proceedings of the National Academy of Sciences 104, 1760–1765, DOI: 10.1073/pnas.0607208104.

USDA, 2014: Swine Enteric Coronavirus Disease Testing Summary Report (Summary Report). United States Department of Agriculture.

USDA, 2020: African swine fever response plan: The Red Book.

USDA, 2022a: USDA Indemnity Values for 2022: Commercial Table. United States Department of Agriculture.

USDA, 2022b: Daily Direct Prior Day Sow and Boar Report. United States Department of Agriculture.

Vergne, T., A. Gogin, and D.U. Pfeiffer, 2015: Statistical Exploration of Local Transmission Routes for African Swine Fever in Pigs in the Russian Federation, 2007–2014. Transboundary and Emerging Diseases 64, 504–512, DOI: 10.1111/tbed.12391.

Weaver, T.R.D., and N. Habib, 2020: Evaluating losses assocatied with African swine fever in the People’s Republic of China and neighboring countries (Working Paper No. 27). Asian Development Bank.

Whitney, M.H., and S.K. Baidoo, 2015: Breeding Boar Nutrient Recommendations and Feeding Management. U.S. Pork Center for Excellence.

Wilkinson, P.J., and A.I. Donaldson, 1977: Transmission studies with African swine fever virus: The early distribution of virus in pigs infected by airborne virus. Journal of Comparative Pathology 87, 497–501, DOI: 10.1016/0021-9975(77)90038-X.

Yoo, D.S., Y. Kim, E.S. Lee, J.S. Lim, S.K. Hong, I.S. Lee, C.S. Jung, H.C. Yoon, S.H. Wee, D.U. Pfeiffer, and G. Fournié, 2021: Transmission Dynamics of African Swine Fever Virus, South Korea, 2019. Emerg Infect Dis 27, 1909–1918, DOI: 10.3201/eid2707.204230.

You, S., T. Liu, M. Zhang, X. Zhao, Y. Dong, B. Wu, Y. Wang, J. Li, X. Wei, and B. Shi, 2021: African swine fever outbreaks in China led to gross domestic product and economic losses. Nat Food 2, 802–808, DOI: 10.1038/s43016-021-00362-1.

Zani, L., K. Dietze, Z. Dimova, J.H. Forth, D. Denev, K. Depner, and T. Alexandrov, 2019: African Swine Fever in a Bulgarian Backyard Farm—A Case Report. Veterinary Sciences 6, 94, DOI: 10.3390/vetsci6040094.

