## Supplementary material for "Estimating the effectiveness of control actions on African swine fever transmission in commercial swine populations in the United States": ss

**Running title: The effectiveness of African swine fever control strategies**

[1]Department of Population Health and Pathobiology, College of Veterinary Medicine, North Carolina State University, Raleigh, NC, USA.

[2]U.S. Department of Agriculture, Animal and Plant Health Inspection Service, Veterinary Services, Strategy and Policy, Center for Epidemiology and Animal Health, Fort Collins, CO, USA.

[3]Veterinary Population Medicine Department, College of Veterinary Medicine, University of Minnesota, St. Paul, MN, USA.

[φ]**share first author**

**Section A.**

*Construction of vehicle movement networks*

Briefly, using the vehicle coordinates and speed we identified the farms that each vehicle visited while also capturing the duration of the visit and the chronological order of the movements (e.g., vehicle X visited farm A for 15 minutes, then traveled for 20 minutes to reach farm B where it stayed for 30 minutes). A farm visit was identified when a vehicle registered a speed of zero km/h within 1.5 km of the coordinates of a farm for at least five minutes (Galvis, Corzo et al., 2022). Network edges between the farms visited by the vehicle were weighted by two factors i) the duration of time each vehicle remained at the farm's premises, which increased the probability of vehicle contamination; and ii) the duration of travel between farms, which decreased the probability of ASF transmission by the vehicle (Galvis, Corzo et al., 2022). The rate of decrease of ASF transmission by vehicles was also dependent on the time of year, reflecting the increased survival of the ASF virus in the environment at colder temperatures (Mazur-Panasiuk et al., 2019). The stability of the ASF virus on contaminated vehicles was assumed to decline linearly over 24 hours in the warm season (Olesen et al., 2018), from April 1st, 2020, to September 30th, 2020, and 72 hours in the cold season (Carlson et al., 2020), from October 1st, 2020 to March 31st, 2020. For example, in the warm season, vehicles that visited both farms A and B within 24-hours would be classed as a contact, however, if the length between the visits was longer than 24-hours they were not considered a contact. The GPS data also contained records of the vehicles visiting cleaning stations, identified by a registered speed of zero km/h within 1.5 km of a known cleaning station for at least five minutes (Galvis, Corzo et al., 2022). We assumed 100% effectiveness of the cleaning and disinfection protocols (De

Lorenzi et al., 2020; Juszkiewicz et al., 2020; Gebhardt et al., 2021; Jiang et al., 2021), removing network edges between farms visited prior to disinfection and farms visited after disinfection.

**Section B.**

*Calibration of ASF transmission parameters using historic PEDV data*

The historic PEDV data used for the calibration covered all registered outbreaks from the first introduction of PEDV in the study area on July 7th, 2013, through to October 31st, 2013 (Machado et al., 2019), for all 2,294 farms included in the ASF model. Prior to the ABC calibration procedure, we extracted the summary statistics from the PEDV data detailing i) the total number of PEDV cases; ii) the mean and maximum number of cases per day for each swine production type; and iii) the mean distance between any two new cases during the outbreak (Table S1), hereon referred as the observed summary statistics. These summary statistics were the foundation for the acceptance criteria in the ABC rejection algorithm. Error-values, detailed in Table S1, were calculated for each observed summary statistic to construct an acceptable range. The calibration algorithm was initiated, simulating ASF epidemics using the priors detailed in Table S2 until ten particles were accepted. After the acceptance of ten particles, the priors were updated using the posterior parameter distributions of these particles, collectively. Subsequent accepted particles were used individually to update the prior distribution. Particles were accepted by comparing their summary statistics, referred to as simulated summary statistics, to the observed summary statistics. If the simulated summary statistics fell within an acceptable error range of the observed summary statistics, the particle was accepted (Sisson et al., 2007; Hartig et al., 2011; Minter and Retkute, 2019). These summary results included the total number of ASF cases during the 140-day simulation, the median and maximum daily cases, and the mean distance between cases infected during the simulation.

**Table S1. Summary statistics for PEDV outbreak data**

| Summary statistic | Observed value | ABC rejection error range |
|---|---|---|
| Total cases in sow farms | 90 | 70 - 110 |
| Median daily cases in sow farms | 2.64 | 1.64 - 3.64 |
| Maximum daily cases in sow farms | 10 | 0 - 20 |
| Total cases in nursery farms | 108 | 88 - 128 |
| Median daily cases in nursery farms | 2.7 | 1.7 - 3.7 |
| Maximum daily cases in nursery farms | 13 | 3 - 23 |
| Total cases in finisher farms | 175 | 150 - 200 |
| Median daily cases in finisher farms | 3.89 | 2.89 - 4.89 |
| Maximum daily cases in finisher farms | 18 | 8 - 28 |
| Average distance between cases (km) | 35.73 | 25.73 - 45.73 |

**Table S2. Prior distributions of ASF transmission parameters calibrated using PEDV outbreak data**

| Parameter | Notation | Prior distribution |
|---|---|---|
| Transmission rate of movements of infected/detected swine | $\beta_n$ | PERT (min = 1.4, mode = 1.6, max = 1.8)* |

| | | |
|---|---|---|
| Transmission rate of movements of exposed swine | $\beta_s$ | PERT (1.4, 1.6, 1.8)* |
| Transmission rate of swine vehicles | $\beta_p$ | PERT (0.00035, 0.0006, 0.00085)* |
| Transmission rate of market vehicles | $\beta_m$ | PERT (0.00025, 0.00049, 0.00075)* |
| Transmission rate of feed vehicles | $\beta_f$ | PERT (0.000024, 0.00003, 0.000034)* |
| Transmission rate of crew vehicles | $\beta_c$ | PERT (0.0001, 0.00027, 0.0004)* |
| Transmission rate of local spread | $\beta_l$ | PERT (0.0005, 0.00075, 0.001)* |
| Surveillance effectiveness (sow) | $L_{sow}$ | Uniform (0.85, 0.95) |
| Surveillance effectiveness (nursery) | $L_{nursery}$ | Uniform (0.15, 0.2) |
| Surveillance effectiveness (finisher) | $L_{finisher}$ | Uniform (0.5, 0.6) |
| Surveillance effectiveness (gilt isolation units) | $L_{gilt}$ | Uniform (0, 0.5) |
| Surveillance effectiveness (boar studs) | $L_{boar}$ | Uniform (0, 0.5) |

*PERT distributions were used as an informative prior for the transmission betas, as it provides a more realistic probability distribution.

**Table S3. Transmission model parameter values for ASF between-farm epidemic spread**

| **Parameter** | **Notation** | **Average value** | **Credible interval (95%)** | **Reference** |
|---|---|---|---|---|
| Transmission rate for swine movements (infected/detected) | $\beta_n$ | 1.60168 | (1.59282, 1.61209) | ABC fitting |
| Transmission rate for swine movements (exposed) | $\beta_s$ | 1.60349 | (1.58731, 1.62410) | ABC fitting |

| | | | | |
|---|---|---|---|---|
| Transmission rate for contaminated swine vehicles | $\beta_p$ | 0.00061 | (0.00060, 0.00063) | ABC fitting |
| Transmission rate for contaminated market vehicles | $\beta_m$ | 0.00050 | (0.00047, 0.00053) | ABC fitting |
| Transmission rate for contaminated feed vehicles | $\beta_f$ | 0.00003 | (0.00003, 0.00003) | ABC fitting |
| Transmission rate for contaminated crew vehicles | $\beta_c$ | 0.00027 | (0.00026, 0.00029) | ABC fitting |
| Transmission rate for local spread | $\beta_l$ | 0.00076 | (0.00075, 0.00076) | ABC fitting |
| Latent period (days) | $\sigma$ | 5 | - | (Hu et al., 2017) |
| Average time to reach the midpoint of the logistic curve (days) | $x0$ | 10* | - | (Malladi et al., 2022) |
| Surveillance effectiveness (sow) | $L_{sow}$ | 0.9012 | (0.8535, 0.9483) | ABC fitting |
| Surveillance effectiveness (nursery) | $L_{nursery}$ | 0.1781 | (0.1544, 0.1986) | ABC fitting |
| Surveillance effectiveness (finisher) | $L_{finisher}$ | 0.5327 | (0.5286, 0.5387) | ABC fitting |

* Once the first case of ASF was detected this was reduced to seven, to reflect the increased awareness of ASF in the region

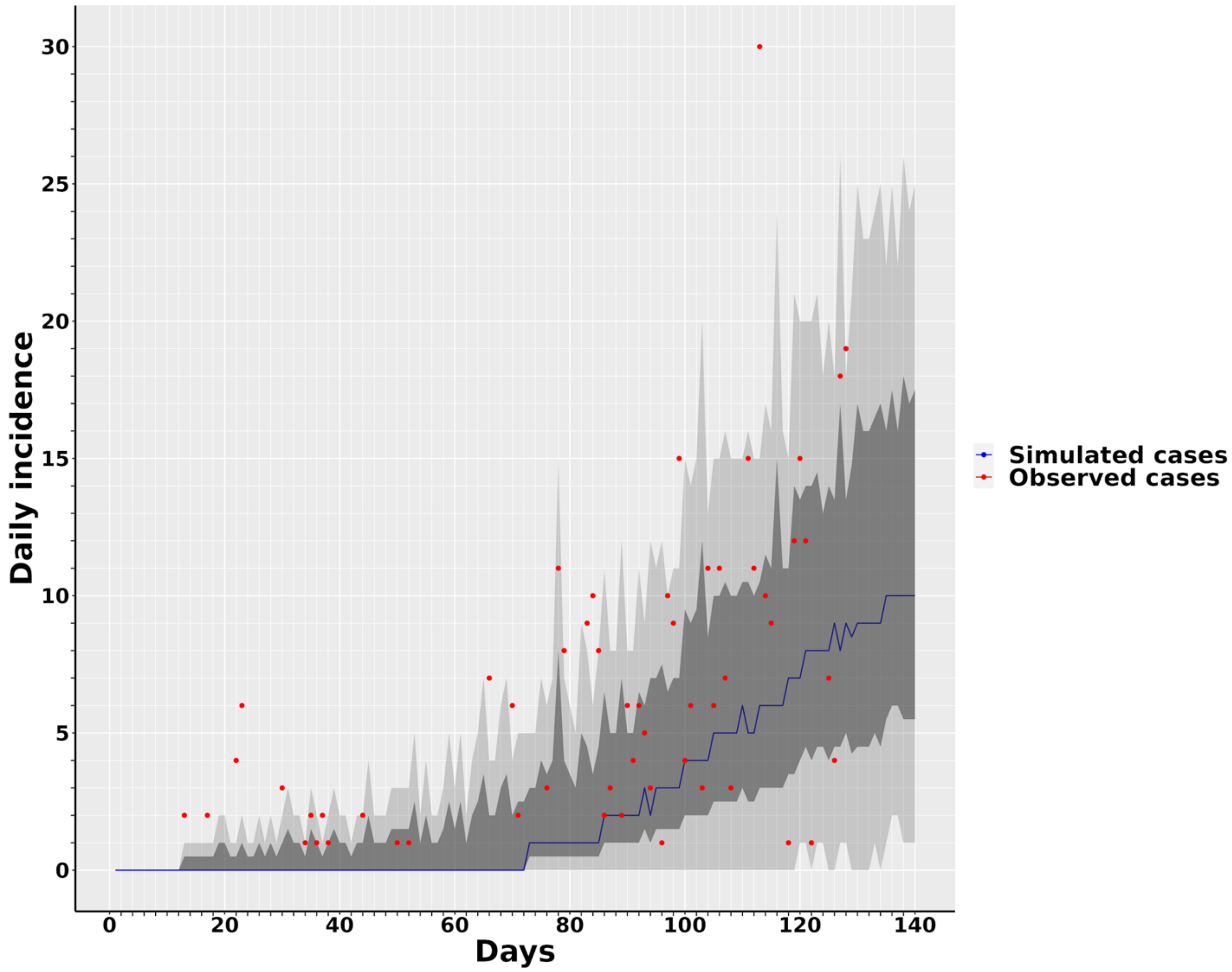


**Figure S1. Comparison of observed PEDV cases and simulated ASF cases over 140 days.** The solid blue line represents the median number of simulated cases, while the dark shade areas represent a 50% interquartile range and the light shade areas maximum and minimum ranges generated by the model. The red dots represent the frequency of outbreaks reported in the observed PEDV data. Uncertainty in the estimated model parameters is reflected by 158 repeated simulations.

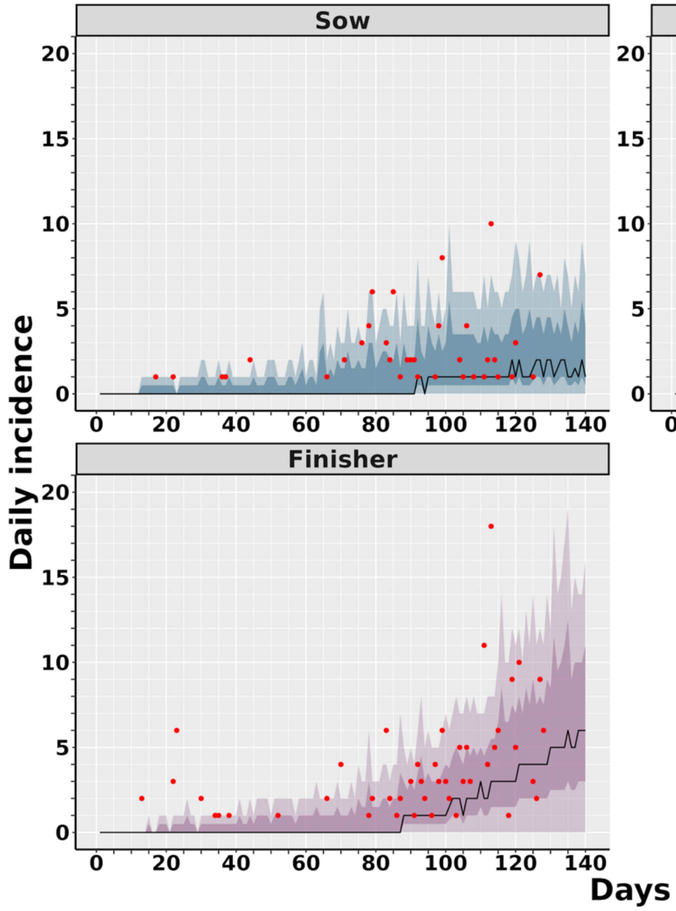


**Figure S2. Comparison of observed PEDV cases and simulated ASF cases from accepted ABC particles, represented by production type.** The solid lines represent the median number of observed cases, while the dark shade areas represent a 50% interquartile range and the light shade areas maximum and minimum ranges generated by the model. The red dots represent the frequency of outbreaks reported in the observed PEDV data. Uncertainty in the estimated model parameters is reflected by 158 repeated simulations.

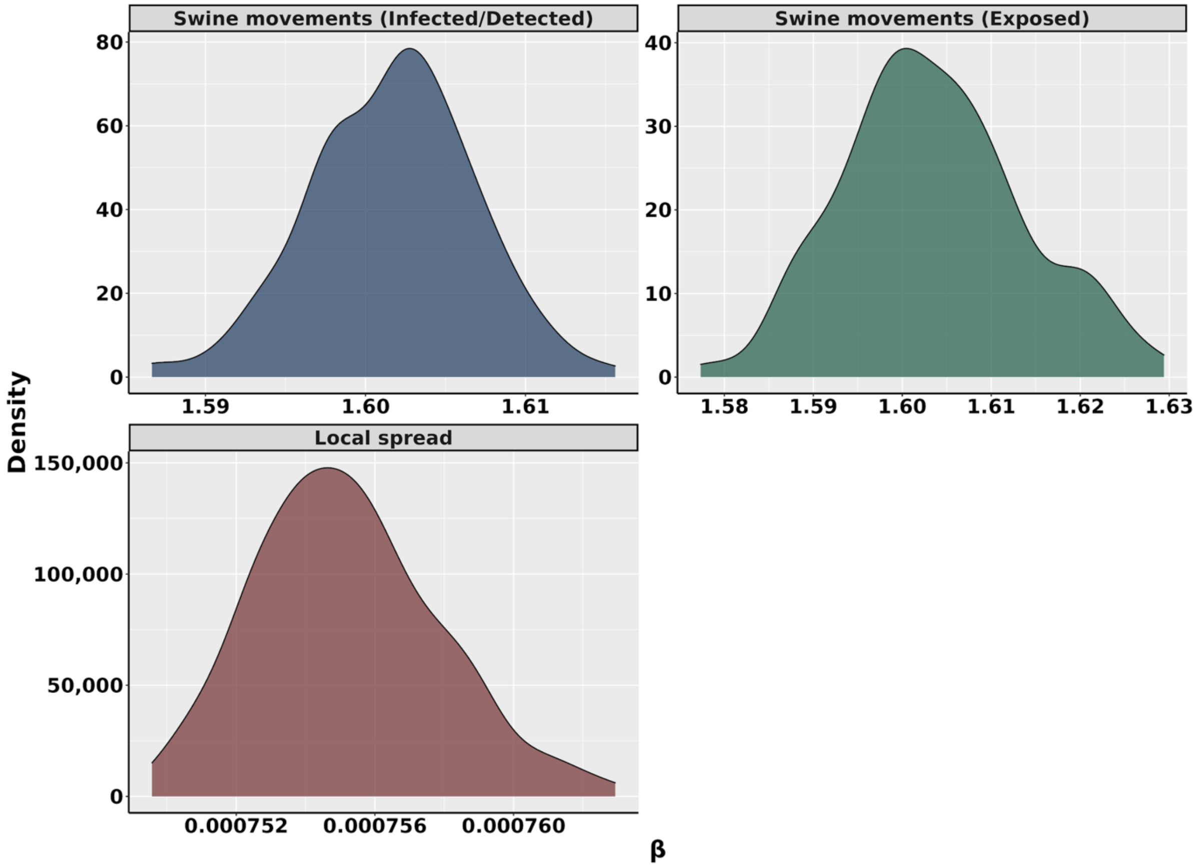


**Figure S3. Posterior distribution of beta parameters for movements of infected and detected swine, movements of exposed swine, and local spread.** Produced from 158 accepted particles from an ABC rejection algorithm

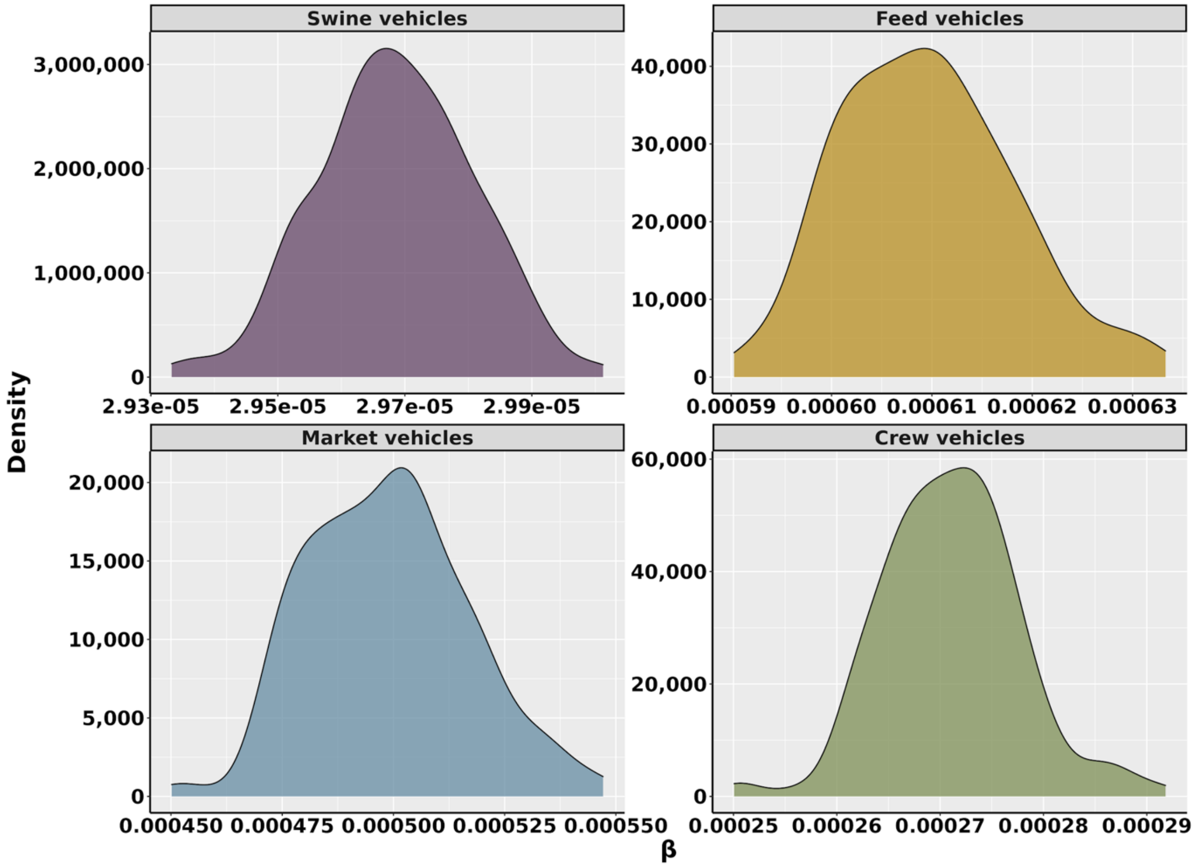


**Figure S4. Posterior distribution of beta parameters for swine vehicles feed delivery vehicles, market vehicles, and crew vehicles.** Produced from 158 accepted particles from an ABC rejection algorithm

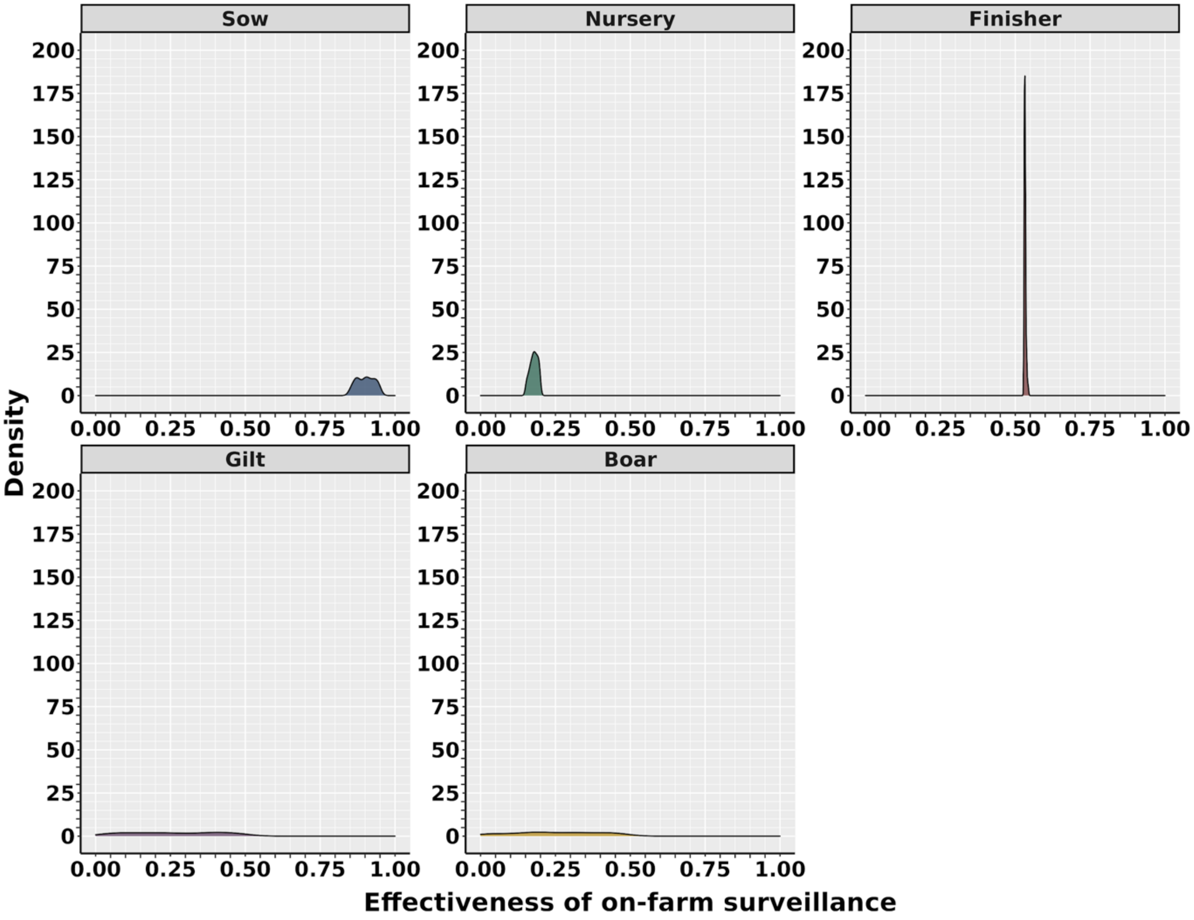


**Figure S5. Posterior distribution of surveillance effectiveness by production type.** Produced from 158 accepted particles from an ABC rejection algorithm

**Section C.**

*Local spread modeling*

Local spread describes short-range transmission from infected farms to nearby susceptible farms that can be explained by geographical proximity (Sanson, 1994). The potential transmission mechanisms that are represented by local spread include mechanical transmission by wildlife, personnel, and the sharing of equipment and tools (Sanson, 1994). Airborne transmission was not represented in the local spread as current evidence suggests that ASF can only spread via aerosols over a one to two-meter distance (Wilkinson and Donaldson, 1977; Olesen et al., 2017; X. Li et al., 2022).

We modeled local spread using a spatial gravity model (Equation 1), where the probability of transmission, $a_{ij}$ , between two farms, $i$ and $j$, was proportional to the product of their swine capacities, $m_i$ and $m_j$, divided by the squared Euclidean distance between the two farms, $d_{ij}^2$.

$$a_{ij} = \frac{m_i m_j}{d_{ij}^2} \qquad \text{Equation 1}$$

Due to the lack of consensus on the distance-dependent spread of ASF (Fodor et al., 2015; Halasa, Boklund et al., 2016; Iglesias et al., 2016; Andraud et al., 2019; Chenais et al., 2019; Boklund et al., 2020; Bradhurst et al., 2021; Pepin et al., 2021), we employed an assortment of cut-off values, ranging from one km to 25 km, to reflect the potential distances of ASF local spread.

*Initial start dates of outbreak*

For simulations that started later than August 13th, 2020, there were less than 140 days left in the study period which would cause the simulation to include calendar dates, such as January 1st,

2021 for which there was no data. To avoid this artifact, once a simulation reached December 31st, 2020, animal and vehicle movement data were used from the beginning of the study period, starting January 1st, 2020 for the remaining days in the simulation (i.e., if there are 14 days left in the simulation after December 31st, 2020, movement from January 1st, 2020 to January 14th, 2020 was used for these 14 days) (Figure S6)

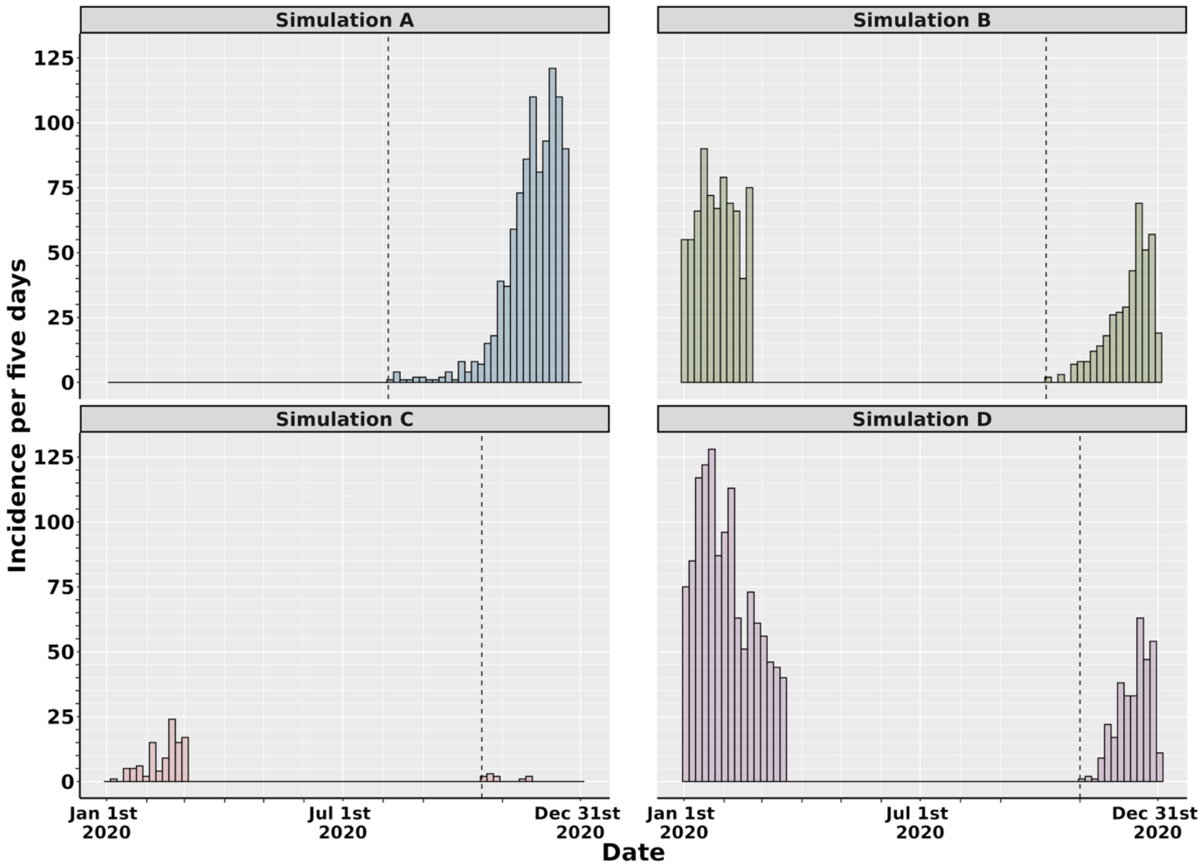


**Figure S6. Example of the simulated period of time used for each initial seed infection.** Each simulation uses 140 days and collects information at each time step along the year 2020. For simulations whose start date (dashed line) was later than August 13th, 2020, animal and vehicle movement data were used from the beginning of the study period (i.e., simulation C and simulation D).

*Base model transmission dynamics*

At the beginning of each simulation (day one), all farms, except the initially infected farm, started in the S compartment of the model (Figure 1). At each time step $t$, starting from day two, the probability of infection (i.e., transition into either the E, I, or D compartment) for each farm, excluding those in the D compartment, was calculated as a function of the force of infection, $\lambda$, for each transmission route (Figure 1). For the between-farm swine movements, the force of infection acting on a susceptible farm $i$ was the product of the transmission rate, represented by $\beta_n$ for movements from farms with exposed animals or $\beta_s$ for movements from farms with infected or detected animals; and the number of farms in each compartment that moved swine to a susceptible farm $i$ on day $t$ ($N_{nit}$ for farms in E, $N_{sit}$ for farms in I, and $N_{ait}$ for farms in D). Farms in the E and I compartments were subjected to a force of infection if they received movements of animals from later compartments, I or D, respectively, which increased the probability of these farms in E and I moving into I or D compartments (Table 1). For example, an exposed farm $i$ that received a movement of swine from a detected farm $j$ is subjected to a force of infection $\lambda_{ait}$ and subsequently has an increased probability, $W_{it}$, of transition into the D compartment. For vehicle movements, the force of infection acting on a susceptible farm $i$, for each vehicle type, was the product of the transmission rate $\beta$; the time elapsed between the vehicle visiting an infected or detected farm $j$, and the same vehicle visiting a susceptible farm $i$, $M_{ijt}$, and the duration of time the vehicle stayed on-farm $i$, $Z_{it}$. Lastly, we calculated the force of infection of local spread on a susceptible farm $i$. This was a product of the transmission rate for local spread $\beta_l$, and the sum of the gravity model values between a farm $i$ and all infectious farms present in the local spread radius. The force of infection of vehicle movements and local spread contributed to the probability of transition from the S compartment to the E compartment only, while swine movements contributed to the probability of transition into the E, I or D compartments, as we assumed that exposed, infected or detected animals would still be exposed, infected or detected in their destination farm, leading the new farm to transition into the respective compartment. The probability of a farm transitioning into another compartment (e.g., E, I, and D) was compared to a transmission threshold which had been sampled from a uniform distribution (0, 1) for each compartment (Figure 1), to determine if the farm would remain in its current compartment or move to the compartment in question.

Farms were also allowed to transition between E and I compartments based on the ASF latent period and I and D compartments based on the rate of detection of ASF infection (Figure 1). The transition of farms from compartment E to compartment I was modulated by the latent period $\sigma$, represented by a positive Poisson distribution, while the transition of farms from compartment I to compartment D was determined by a logistic function $f(x)$ characterizing the rate of clinical disease, an approach used in previous literature (Galvis, Corzo et al., 2022). In our model, $f(x)$ was dependent on the effectiveness of surveillance in each farm, $L$, which varied by production type; the logistic growth rate, $k$; the length of time the farm had spent in the I compartment; and the average time taken to reach the midpoint of the logistic curve using a mortality trigger of 5 per 1000 heads of swine, $x0$ (Malladi et al., 2022) (Table S3 and Figure S7).

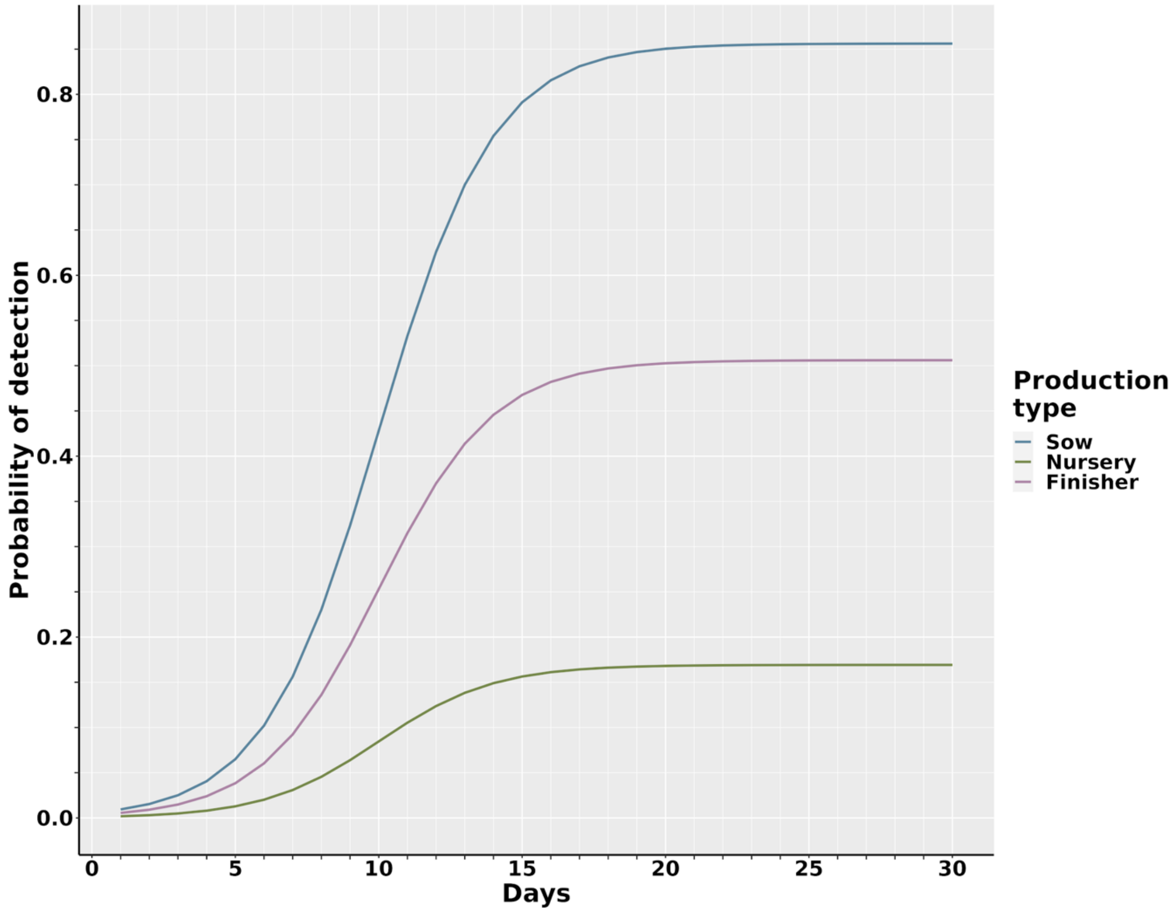


**Figure S7. Probability of detection of ASF-infected farms.** Detection probabilities are based on a logistic distribution, where the maximum detection probability was fixed as the median surveillance effectiveness for each production type (extracted from the respective ABC-MCMC posteriors). For sow farms this was 0.9, for nursery farms this was 0.18 and for finisher farms this was 0.53. The time elapsed to reach the midpoint of the detection curve was ten days for all farm types.

*Sensitivity analysis*

As the baseline model may be sensitive to assumed model parameters, we considered variations in the transmission parameters ($\beta_n$, $\beta_s$, $\beta_p$, $\beta_m$, $\beta_f$, $\beta_c$ and $\beta_l$) and surveillance values ($L_{sow}$, $L_{nursery}$ and $L_{finisher}$) calibrated by the PEDV outbreak data, in addition to variations in the latent period of the virus ($\sigma$) and the average time to reach the midpoint of the detection rate logistic curve ($x0$). For each sensitivity variation, we initiated infection at each farm in the study 50 times, equating to 114,700 simulation repeats. We then used a Kruskal-Wallis test followed by a post-hoc Dunn's test (with a Benjamini-Hochberg correction), or a Mann Whitney U test for parameters with only two values investigated, to compare the number of infections at the end of 140 days for each parameter variation.

**Section D.**

**Table S4. Description of transmission routes which are prevented under the control and eradication actions**

| **Transmission route** | **Quarantine and depopulation** | **72-hour movement standstill** | **Contact tracing†** | **Depopulation of direct contacts** | **Control areas and surveillance zones** |
|---|---|---|---|---|---|
| *Movement of swine (exposed)* | - | ✓ | - | ✓ | - |
| *Movement of swine (infected)* | - | ✓ | - | ✓ | - |
| *Movement of swine (detected)* | ✓ | ✓ | - | ✓ | - |
| *Local spread* | ✓* | - | - | ✓* | - |

| | | | | | |
|---|---|---|---|---|---|
| *Swine vehicles* | ✓ | ✓ | - | ✓ | - |
| *Market vehicles* | ✓ | ✓ | - | ✓ | - |
| *Crew vehicles* | ✓ | ✓ | - | ✓ | - |
| *Feed vehicles* | ✓ | ✓ | - | ✓ | - |

* Local spread is only impacted once the farms have been depopulated. While they are waiting for depopulation, ASF can still be transmitted through this transmission route.

† Contact premises would be quarantine under network-based controls according to the USDA's current ASF response plan.

*Sample size calculation*

Below is the sample size calculation (Equation 2) used for sampling the farms to be tested within the surveillance zone and animals to be tested within all farms. This calculation was taken from (Cannon, 2001).

$$n \cong (1 - (1 - \gamma)^{1/\alpha D})(N - (\alpha D - 1)/2) \qquad \text{Equation 2}$$

Where $n$ is the number of animals that should be sampled, which is estimated in relation to the sensitivity of the testing procedure, α; the prevalence, $D$; and the population size, N, with a level of confidence $\gamma$.

**Section E.**

*ASF dissemination over 140 days*

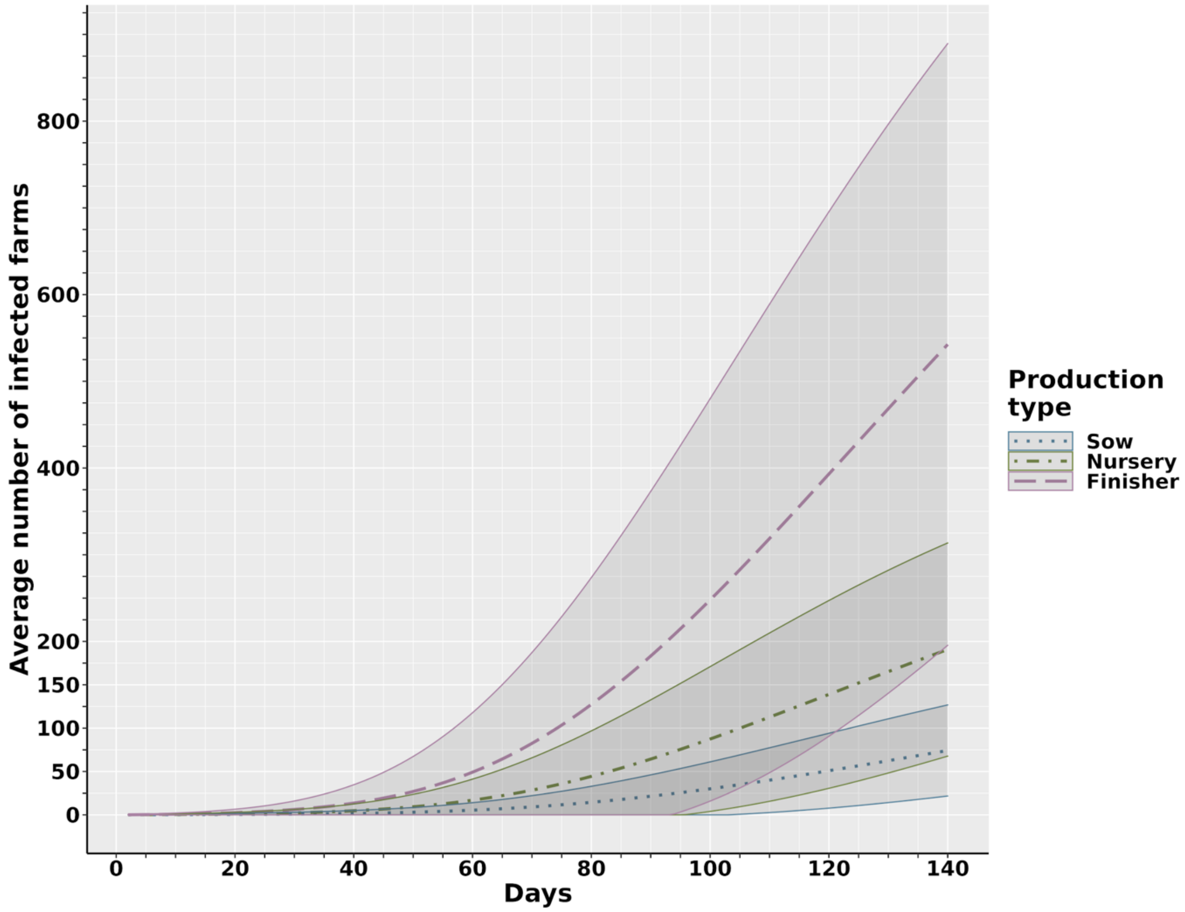


**Figure S8. The average number of secondary ASF cases by farm type accumulated over 140 days of simulated outbreaks.** The dashed and dotted lines represent the mean at time *t* (days) for each production type while the solid lines and shaded areas represent the standard deviation at time *t*.

*Sensitivity results*

The sensitivity analysis highlighted statistically significant sensitivity of the model to several parameters. This included the length of the latent period, average time to detection, the local spread cutoff, the date of initial infection and the transmission rate values. However, it is pertinent to be cautious when interpreting these results, as the substantial number of simulations used to assess the sensitivity could cause very slight differences in secondary cases to be statistically significant. When we consider biological significance instead, we see pronounced changes in secondary cases when the following parameters are changed: i) length of latent period; ii) local spread cutoff; iii) date of initial infection; and iv) the transmission rates for local spread, swine vehicles, crew vehicles and feed vehicles.

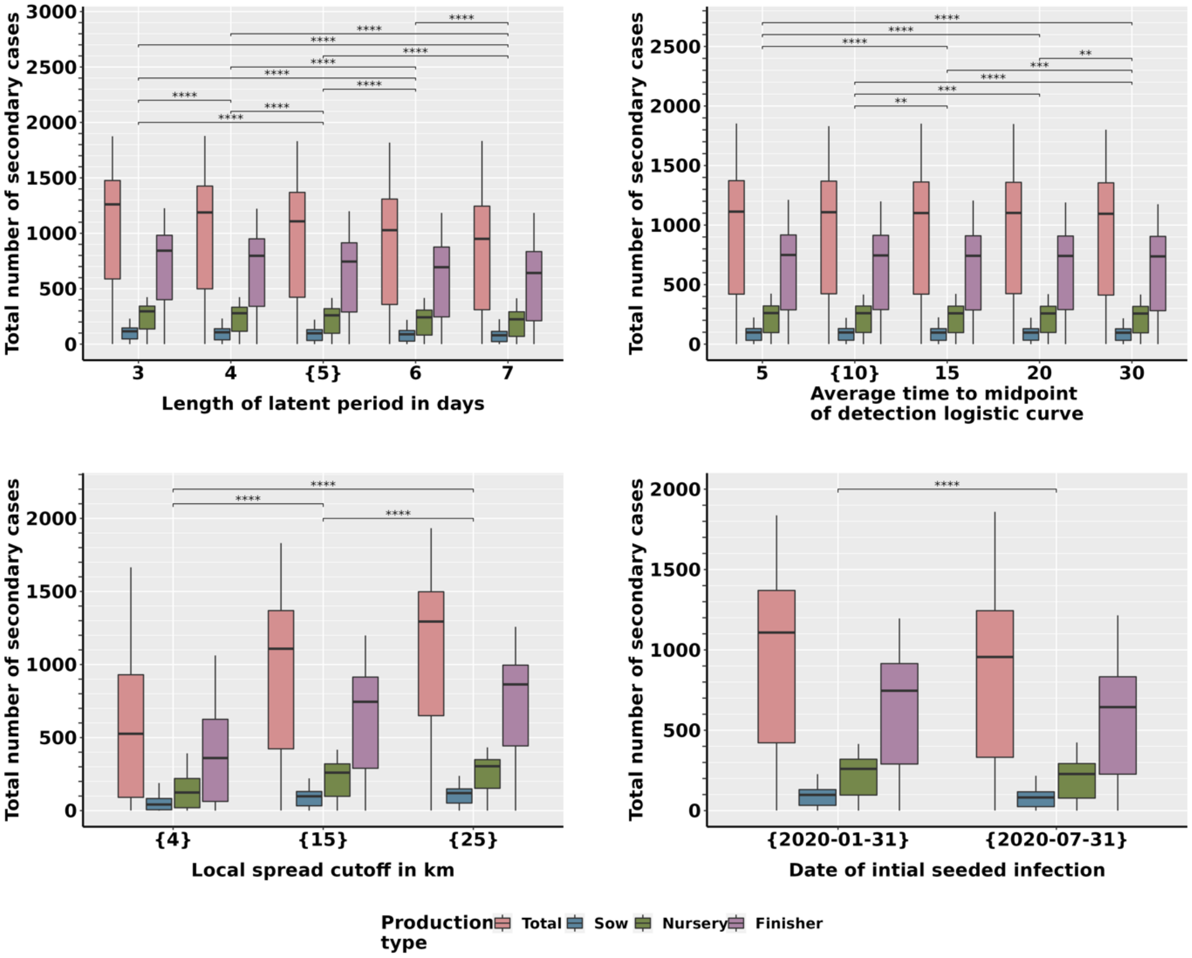


**Figure S9. Effect of changing the length of the latent period, the average time to the midpoint of the detection curve, local spread cutoff and date of initial seeded infection on**

**the total number of secondary ASF cases over 140 days.** The braces (i.e., {}) indicate the values that were used in the baseline model. Statistical analysis using either a Kruskal Wallis test with a post-hoc Dunn's test, or a Mann Whitney test was used to compare the total number of secondary cases between the "Total" production type. The asterisks represent the level of significance as follows: "*" $p \leq 0.05$; "**" $p \leq 0.01$; "***" $p \leq 0.001$; "****" $p \leq 0.0001$.

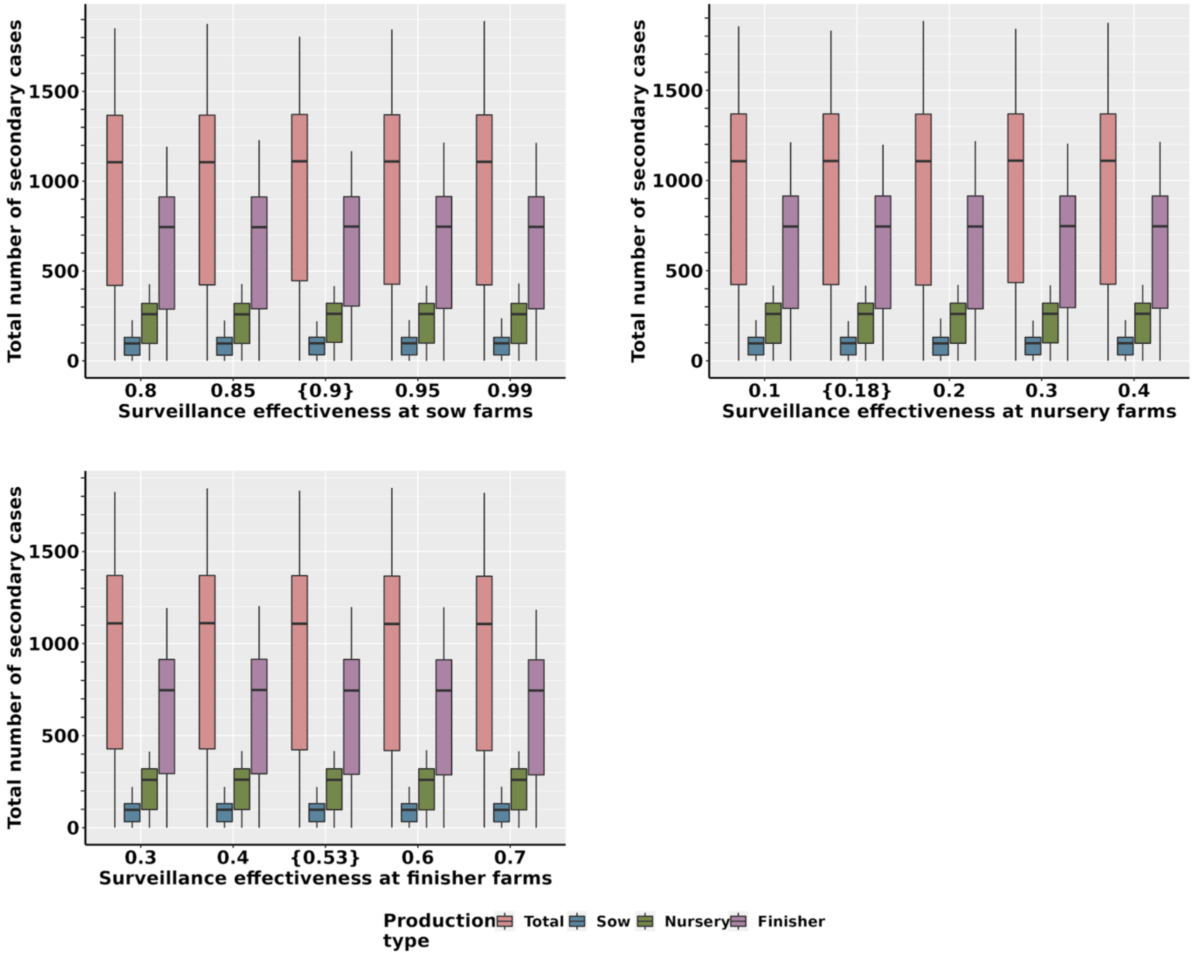


**Figure S10. Effect of changing the surveillance effectiveness at sow farms, nursery farms and finisher farms on the total number of secondary ASF cases over 140 days.** The braces (i.e., {}) indicate the values that were used in the baseline model. Statistical analysis using a Kruskal Wallis test with a post-hoc Dunn's test was used to compare the total number of secondary cases between the "Total" production type. No statistically significant difference was found.

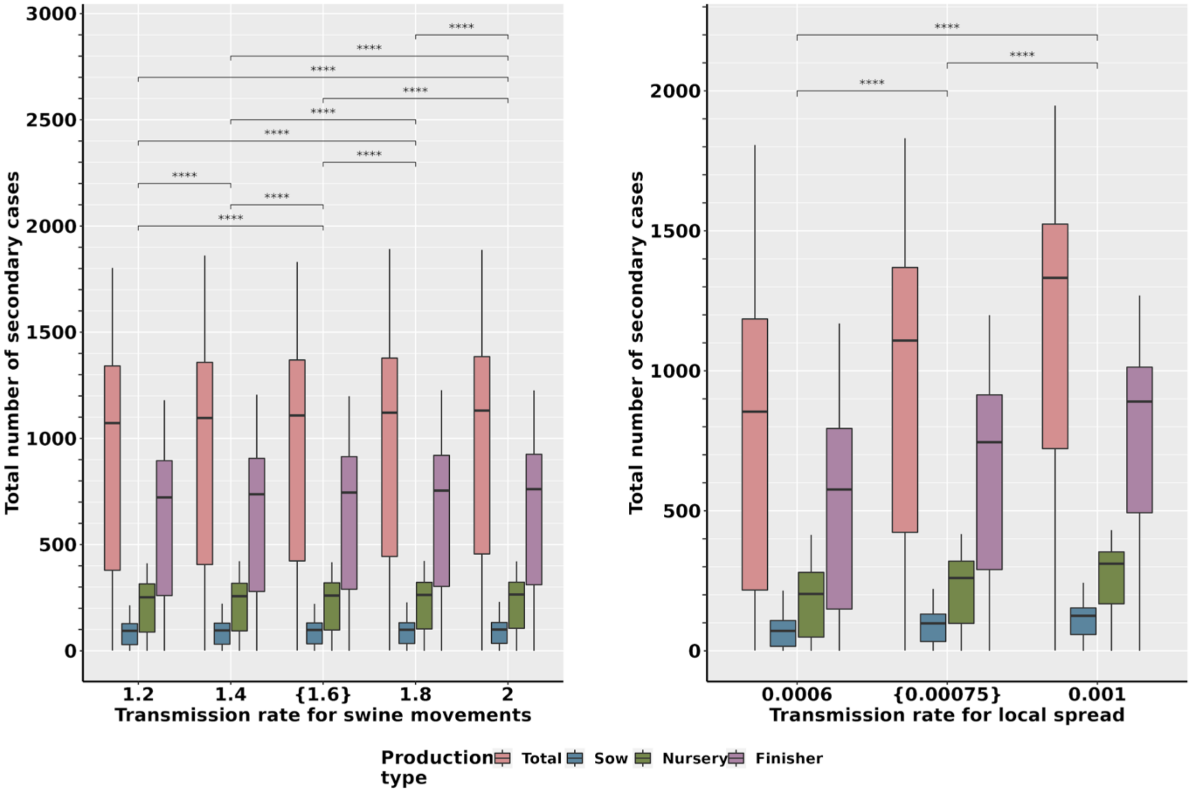


**Figure S11. Effect of changing the transmission rate for swine movements (exposed, infected, and detected swine) and the transmission rate for local spread on the total number of secondary ASF cases over 140 days.** The braces (i.e., {}) indicate the values that were used in the baseline model. Statistical analysis using a Kruskal Wallis test with a post-hoc Dunn's test was used to compare the total number of secondary cases between the "Total" production type. The asterisks represent the level of significance as follows: "*" $p \leq 0.05$; "**" $p \leq 0.01$; "***" $p \leq 0.001$; "****" $p \leq 0.0001$.

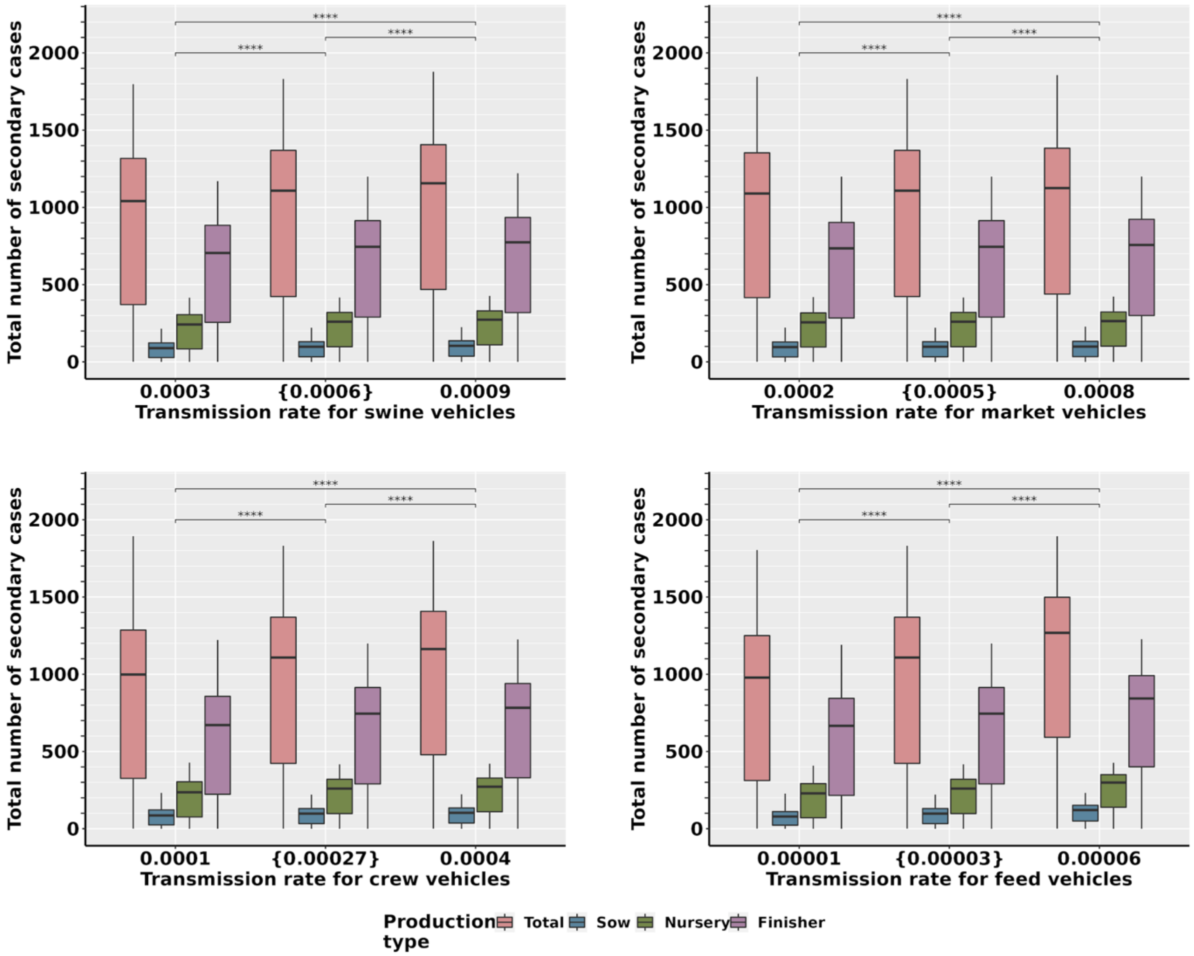


**Figure S12. Effect of changing the transmission rate for swine vehicles, market vehicles, crew vehicles, and feed vehicles on the total number of secondary ASF cases over 140 days.** The braces (i.e., {}) indicate the values that were used in the baseline model. Statistical analysis using a Kruskal Wallis test with a post-hoc Dunn's test was used to compare the total number of secondary cases between the "Total" production type. The asterisks represent the level of significance as follows: "*" $p \leq 0.05$; "**" $p \leq 0.01$; "***" $p \leq 0.001$; "****" $p \leq 0.0001$.

*Contribution of transmission routes to ASF dissemination*

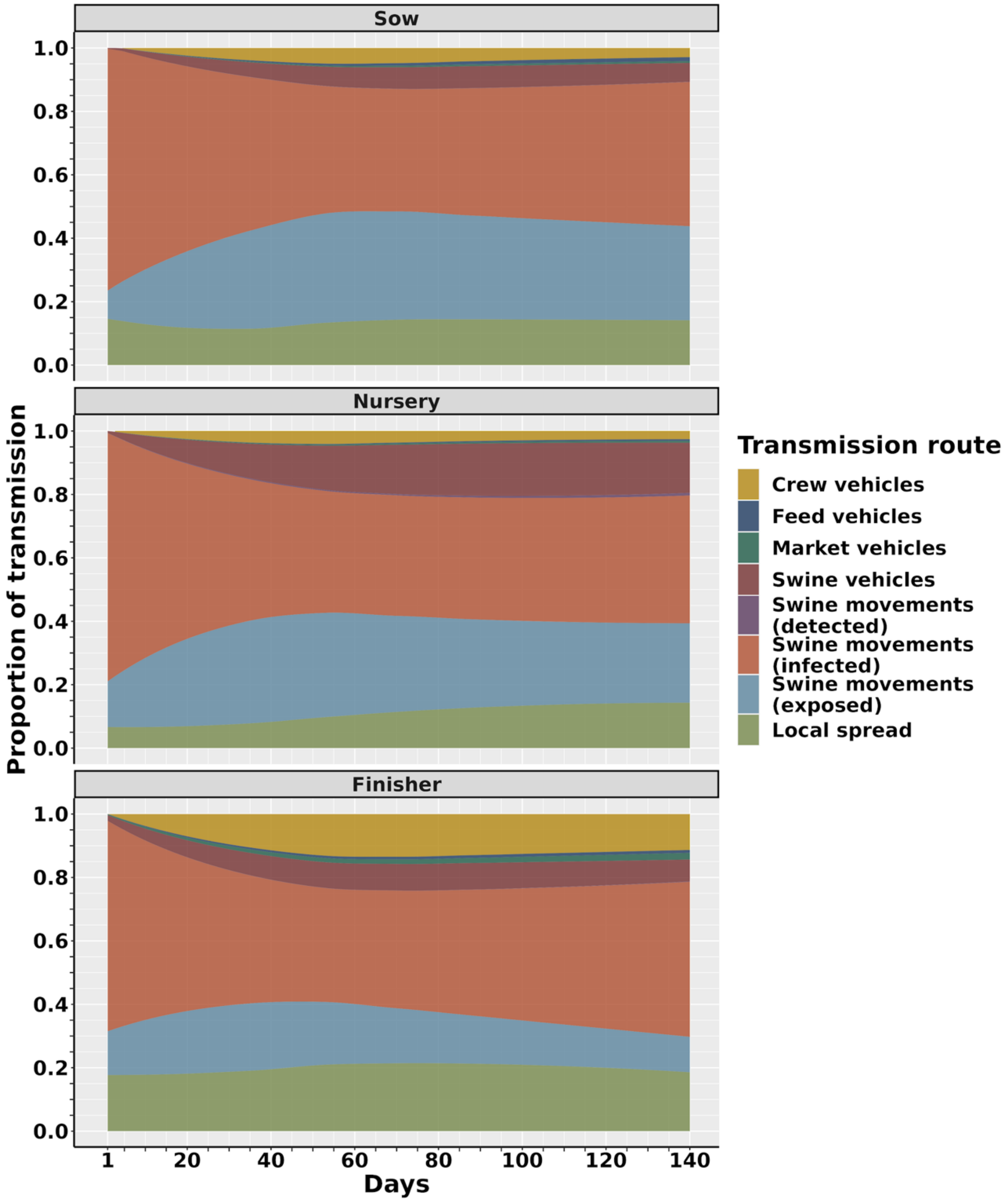


**Figure S13. Contribution of transmission routes to ASF transmission for 140 days of simulated outbreaks, separated by the production type.** The y-axis represents the mean proportion of each transmission route on a given day for each production type. The proportion was then averaged at each time point and smoothed using a generalized linear model.

**Table S5. The average contribution of each transmission route to ASF dissemination over 60 days of ASF outbreaks, as a proportion of the total force of infection for the total population and by production type.**

| Transmission route | Contribution to all infections (SD) | Contribution to sow infections (SD) | Contribution to nursery infections (SD) | Contribution to finisher infections (SD) |
|---|---|---|---|---|
| Swine movements (infected) | 47.7% (±11.2%) | 52.1% (± 13.3%) | 49.3% (±14.0%) | 44.0% (±11.1%) |
| Swine movements (exposed) | 23.3% (±4.6%) | 26.8% (±8.8%) | 28.6% (±7.5%) | 19.4% (±4.3%) |
| Local spread | 14.6% (±3.3%) | 11.9% (±4.1%) | 7.7% (±2.6%) | 18.6% (±4.4%) |
| Swine vehicles | 6.9% (±2.7%) | 3.9% (±1.9%) | 9.2% (±4.3%) | 6.1% (±2.1%) |
| Crew vehicles | 6.4% (±3.1%) | 3.1% (±1.7%) | 2.9% (±1.4%) | 8.7% (±4.2%) |
| Market vehicles | 0.7% (±0.3%) | 0.2% (±0.1%) | 0.2% (±0.1%) | 1.0% (±0.4%) |
| Feed vehicles | 0.4% (±0.2%) | 0.3% (±0.2%) | 0.1% (±0.1%) | 0.5% (±0.2%) |
| Swine movements (detected) | 0.1% (±0.1) | 0.03% (±0.03%) | 0.3% (±0.1%) | 0.03% (±0.02%) |

**Table S6. The average contribution of each transmission route to ASF dissemination over 140 days of ASF outbreaks, as a proportion of the total force of infection for the total population and by production type.**

| Transmission route | Contribution to all infections (SD) | Contribution to sow infections (SD) | Contribution to nursery infections (SD) | Contribution to finisher infections (SD) |
|---|---|---|---|---|
| Swine movements (infected) | 44.7% (±8.2%) | 46.1% (±10.2%) | 43.4% (±10.5%) | 42.9% (±7.8%) |
| Swine movements (exposed) | 21.2% (±3.9%) | 29.7% (±6.4%) | 27.8% (±5.2%) | 16.4% (±4.1%) |
| Local spread | 16.4% (±2.7%) | 13.3% (±2.9%) | 10.8% (±3.3%) | 19.7% (±3.1%) |
| Swine vehicles | 8.7% (±2.4%) | 5.4% (±1.8%) | 13.2% (±4.5%) | 7.2% (±1.2%) |
| Crew vehicles | 7.9% (±2.4%) | 3.5% (±1.3%) | 3.0% (±0.9%) | 10.9% (±3.4%) |
| Market vehicles | 1.0% (±0.4%) | 0.4% (±0.2%) | 0.4% (±0.2%) | 1.4% (±0.5%) |
| Feed vehicles | 0.6% (±0.2%) | 0.7% (±0.4%) | 0.3% (0.2%) | 0.7% (±0.2%) |
| Swine movements (detected) | 0.2% (±0.1%) | 0.04% (±0.03%) | 0.5% (0.2%) | 0.05% (±0.03%) |

*Distance between seeded and secondary infections*

Throughout the first 60 days of the outbreaks, the median distance between seeded infection and secondary cases was 27.6 km (IQR: 13.4-49.1 km) but we also observed long distance spread up to a maximum of 474.0 km (Figure S14). Infections seeded in sow farms showed the farthest spread with a median distance of 40.3 km (IQR: 20.5-70.2 km), while infections seeded in nursery farms had a median distance of 26.7 km (IQR: 13.8-45.6 km) and infections seeded in finisher farms had a median distance of 25.0 km (IQR: 11.9-44.3 km) (Figure S14). The maximum distance between a seeded infection and secondary infection was 474.0 km for infections seeded in finisher farms, while infections seeded in nursery and sow farms generated maximum distances of 373.8 km and 387.3 km, respectively. Over 140 days of outbreaks, the median distance between seeded and secondary infection was 45.2 km (IQR: 27.5-71.8 km) with a maximum of 510.9 km (Figure S15).

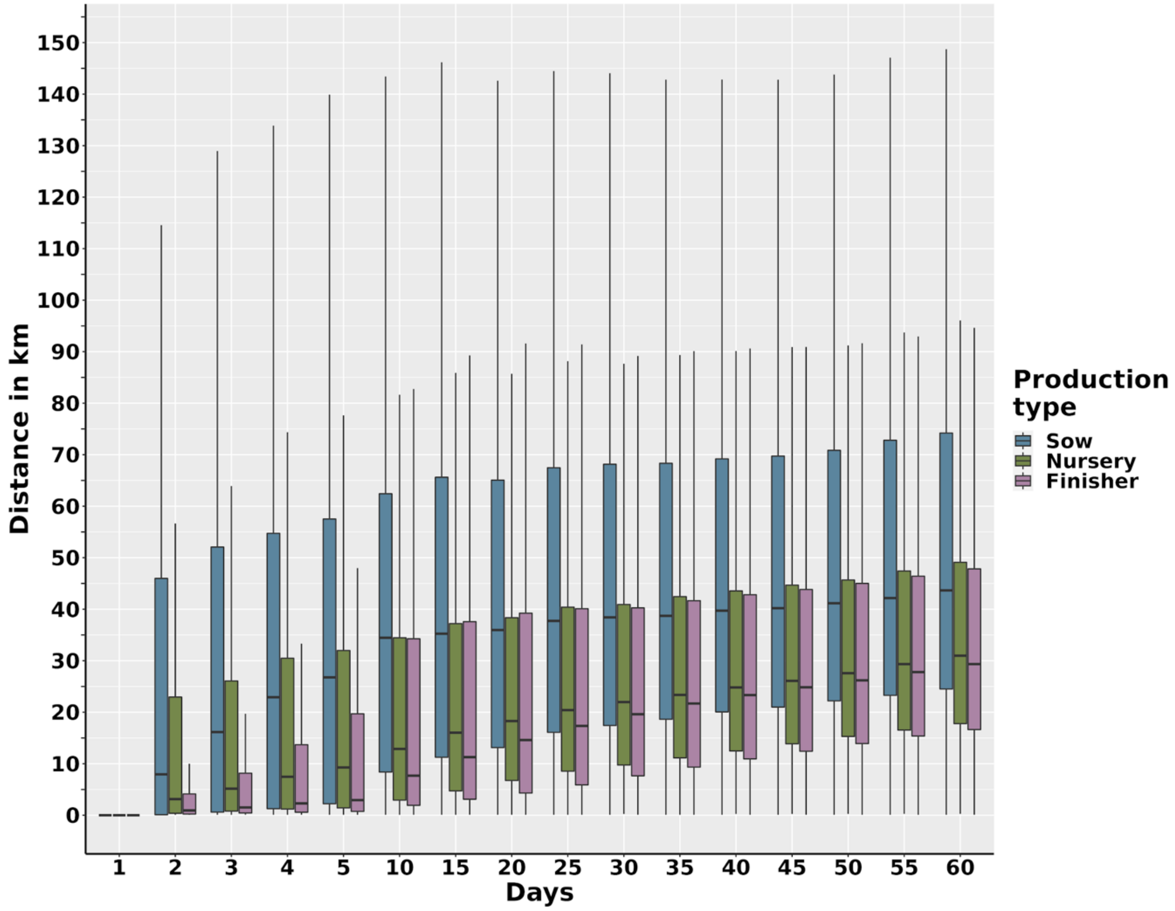


**Figure S14. Average distance between the initial seeded infection and the secondary infections, by production type for the first 60 days of ASF dissemination.** For each simulation, the Euclidean distance between the initial infection and its secondary cases was calculated and averaged at each time shown. The boxplots present the range, interquartile range, and median. Outliers were retained in the dataset but are not shown.

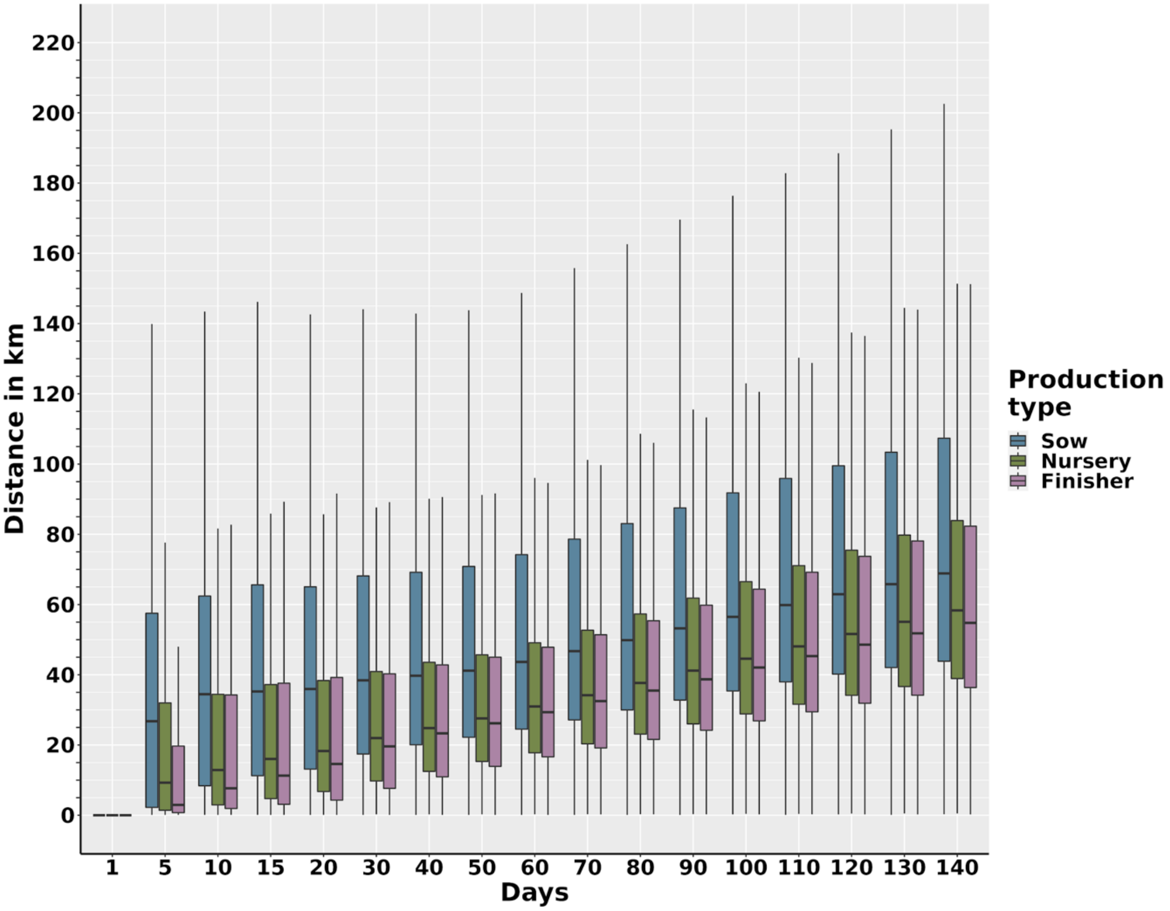


**Figure S15. Average distance between the initial seeded infection and the secondary infections, by production type for the first 140 days of ASF dissemination.** For each simulation, the Euclidean distance between the initial infection and its secondary cases was calculated and averaged for each time shown. The boxplots present the range, interquartile range, and median. Outliers were retained in the dataset but are not shown.

*Between production company transmission*

We also demonstrated that regardless of which company ASF was seeded in, companies not commercially associated would exhibit secondary infections (Figure S16). In particular, the dissemination of ASF among companies resulted in greater numbers of infections in finisher farms, followed by nursery farms and sow farms (Figure S16). At the end of 60 days of outbreaks, company A was the most affected, with a median of 0.7% (IQR: 0.2%-2.4%) of farms infected as a result of a seeded infection in company C (Figure S16). Company B was the least affected with a median of 0.0% (IQR: 0.0%-0.9%) of farms infected as a result of a seeded infection in company C. At the end of 140 days of outbreaks company C was the most affected company when the infection was seeded in company A with a median of 36.8% (IQR: 10.3%-57.6%) of farms infected, however, they were also the least affected company when infection was seeded in company B with a median of 10.6% (IQR: 0.6%-35.2%) of farms infected.

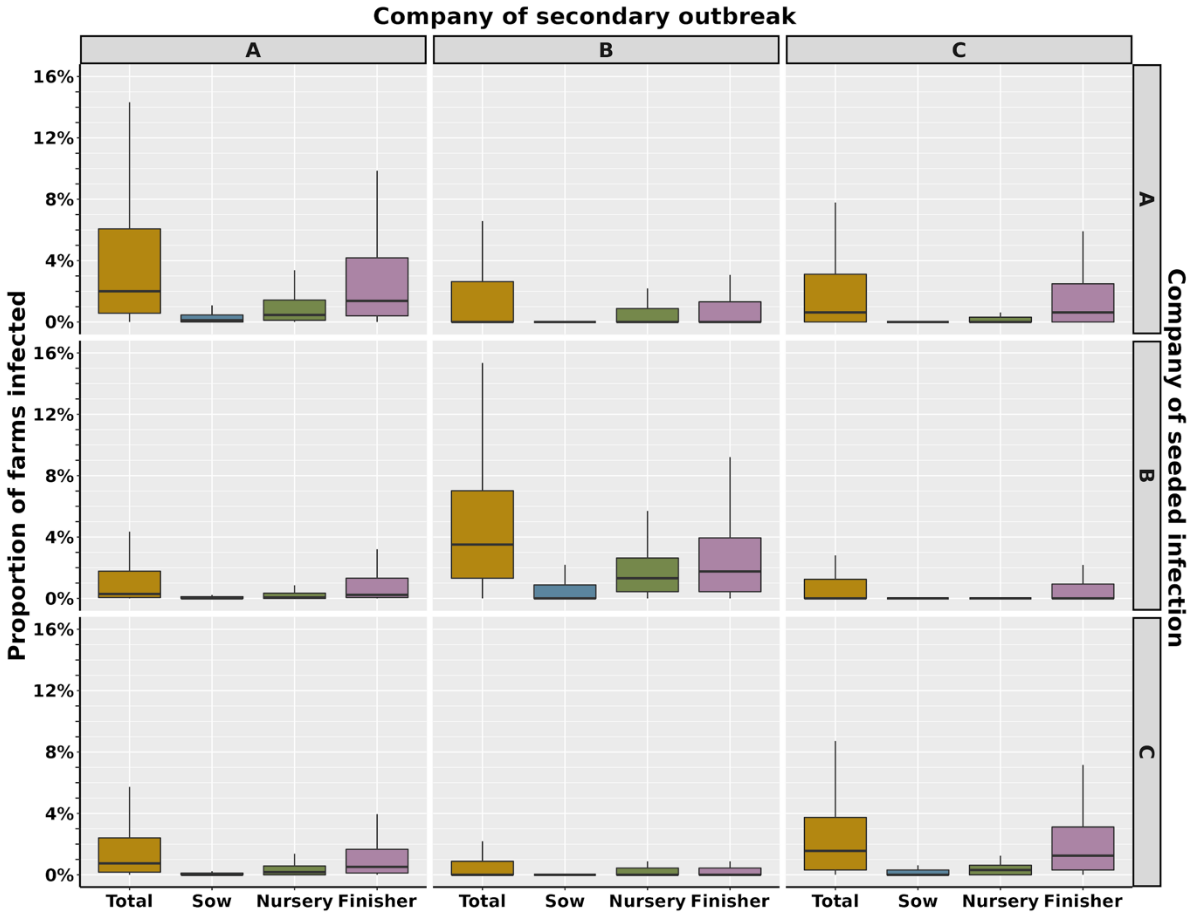


**Figure S16. The average proportion of secondary infections in each company over 60 days of ASF dissemination, relative to the company in which the ASF outbreak was seeded.** Labels on the right vertical axis represent the company in which the initial outbreak was seeded. Labels on the top horizontal axis represent the company in which the cases are being measured.

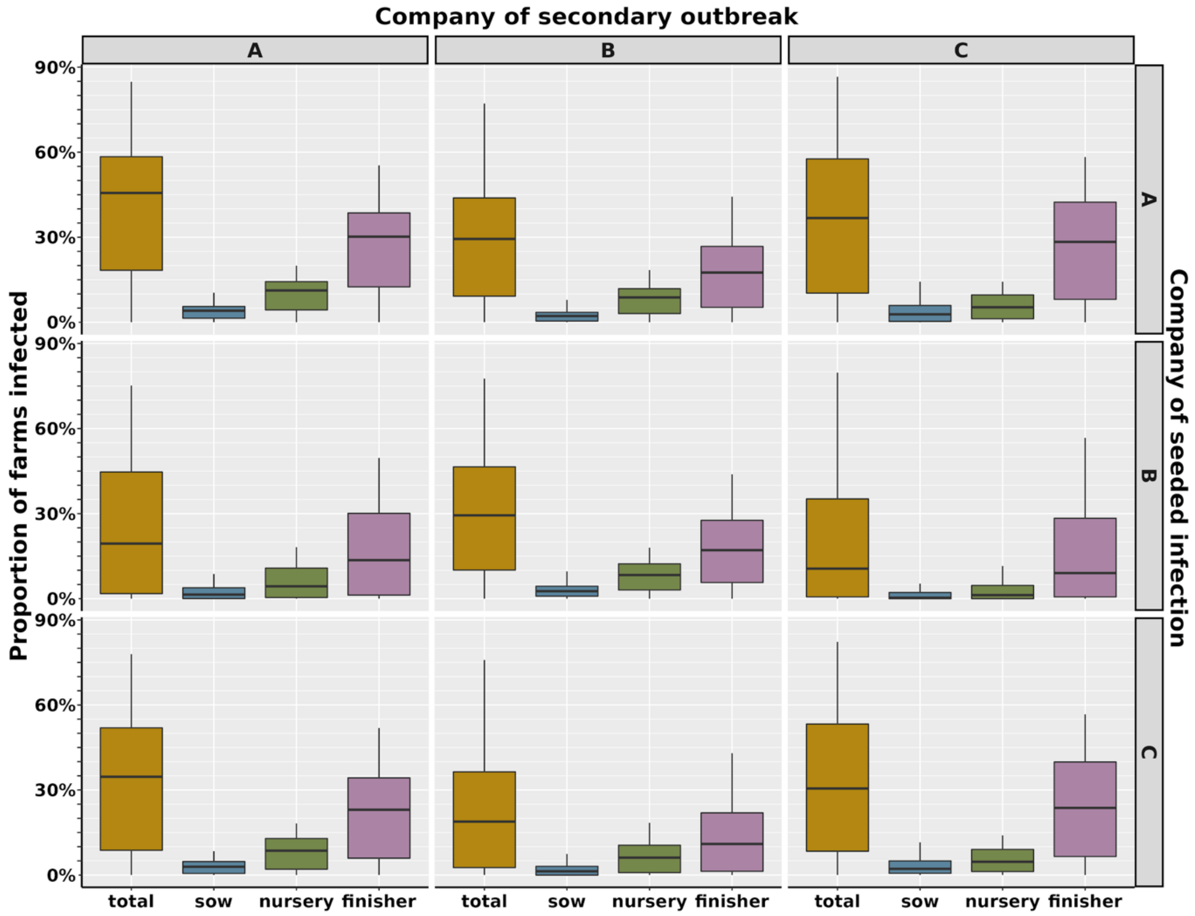


**Figure S17. The average proportion of secondary infections in each company over 140 days of ASF dissemination, relative to the company in which the ASF outbreak was seeded.** Labels on the right vertical axis represent the company in which the initial outbreak was seeded. Labels on the top horizontal axis represent the company in which the cases are being measured.

*Effectiveness of control actions on the number of secondary cases*

**Table S7. Reduction in secondary cases under different control scenarios**

| Scenario | Days | Total reduction in infections (IQR) | Reduction in sow infections (IQR) | Reduction in nursery infections (IQR) | Reduction in finisher infections (IQR) |
|---|---|---|---|---|---|
| 1 | 60 | 11.1% (-69.5%, 56.1%) * | 20.0% (-44.4%, 57.1%) | 16.1% (-54.4%, 56.4%) | 10.0% (-66.8%, 54.8%) |
| | 140 | 7.8% (14.1%, 32.9%) | 12.0% (-14.4%, 37.4%) | 8.5% (-11.3%, 30.7%) | 6.5% (-14.7%, 30.5%) |
| 2 | 60 | 14.5% (-64.4%, 57.4%) | 23.5% (-38.5%, 60.0%) | 20.0% (-50.0%, 58.3%) | 13.6% (-63.1%, 56.4%) |
| | 140 | 8.7% (-12.9%, 34.5%) | 13.1% (-13.2%, 38.6%) | 9.3% (-10.2%, 32.1%) | 7.4% (-13.6%, 32.0%) |
| 3 | 60 | 34.4% (-34.6%, 71.4%) | 43.2% (0.0%, 68.4%) | 39.4% (-20.6%, 70.7%) | 33.3% (-35.3%, 69.2%) |
| | 140 | 30.5% (9.0%, 67.8%) | 36.7% (12.9%, 65.5%) | 29.5% (9.6%, 62.8%) | 27.8% (7.4%, 62.6%) |
| 4 | 60 | 33.5% (-35.5%, 70.2%) | 43.8% (0.0%, 67.4%) | 40% (-20.0%, 70.0%) | 32.1% (-37.5%, 67.6%) |
| | 140 | 35.9% (16.2%, 68.4%) | 47.0% (27.1%, 68.5%) | 39.0% (21.9%, 65.6%) | 31.4% (11.9%, 62.6%) |
| 5 | 60 | 47.4% (-13.2%, 77.6%) | 50.0% (9.1%, 75.0%) | 50.0% (0.0%, 76.8%) | 45.1% (-15.6%, 75.4%) |
| | 140 | 79.0% (56.6%, 93.7%) | 82.1% (66.7%, 91.8%) | 79.3% (60.6%, 91.4%) | 76.6% (52.4%, 91.0%) |

*We attribute these negative values observed in the interquartile range to the stochastic nature of the model which may change from simulation to simulation, including the latent period of the virus and the thresholds for transmission and detection.

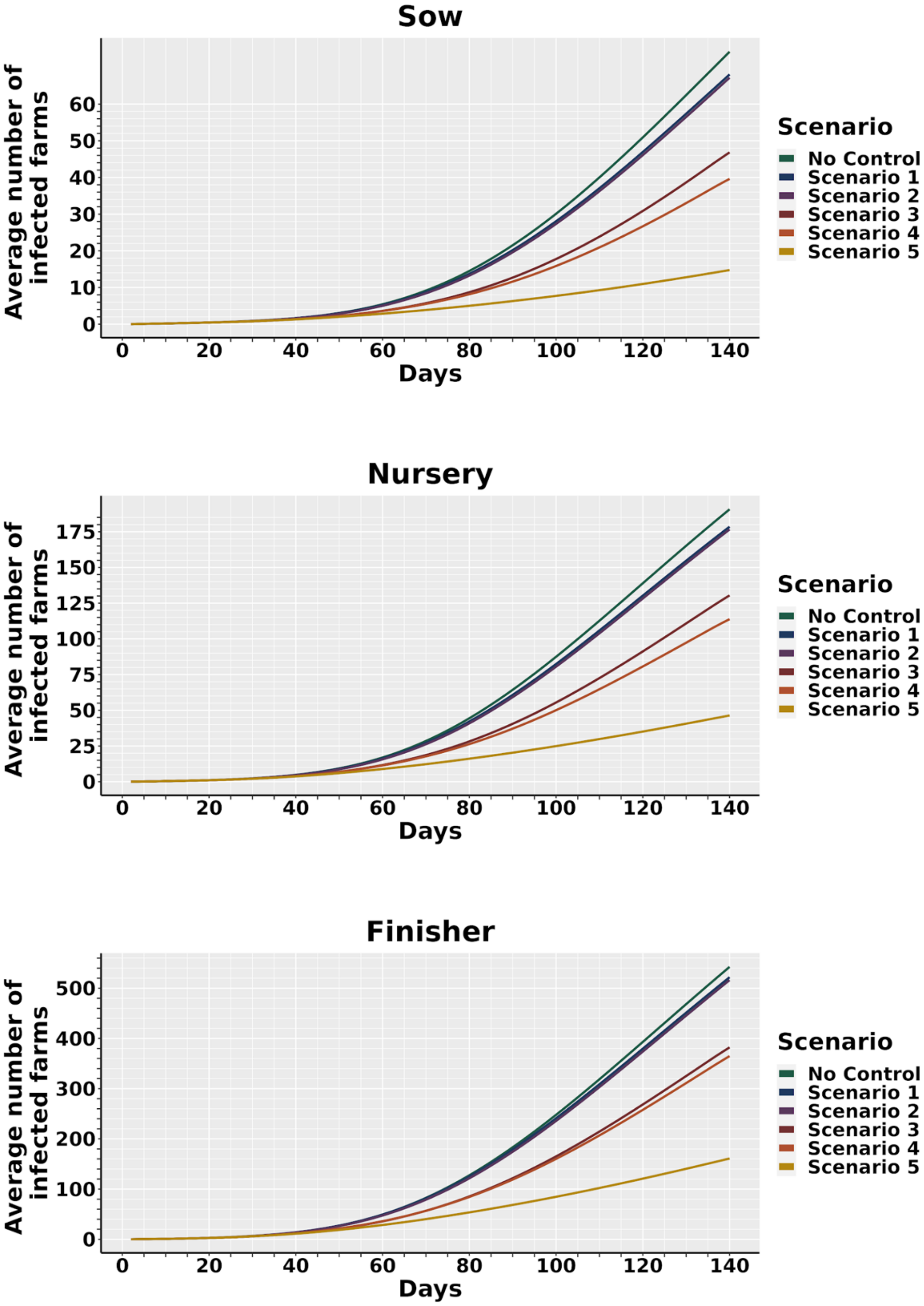

**Figure S18. Average number of secondary cases accumulated over 140 days of the outbreak, under five different control scenarios.** The colored lines represent the mean prevalence after *t* days, under each control scenario. Control scenarios were implemented upon detection of the first ASF case. Control scenarios are described in detail in Table 1.

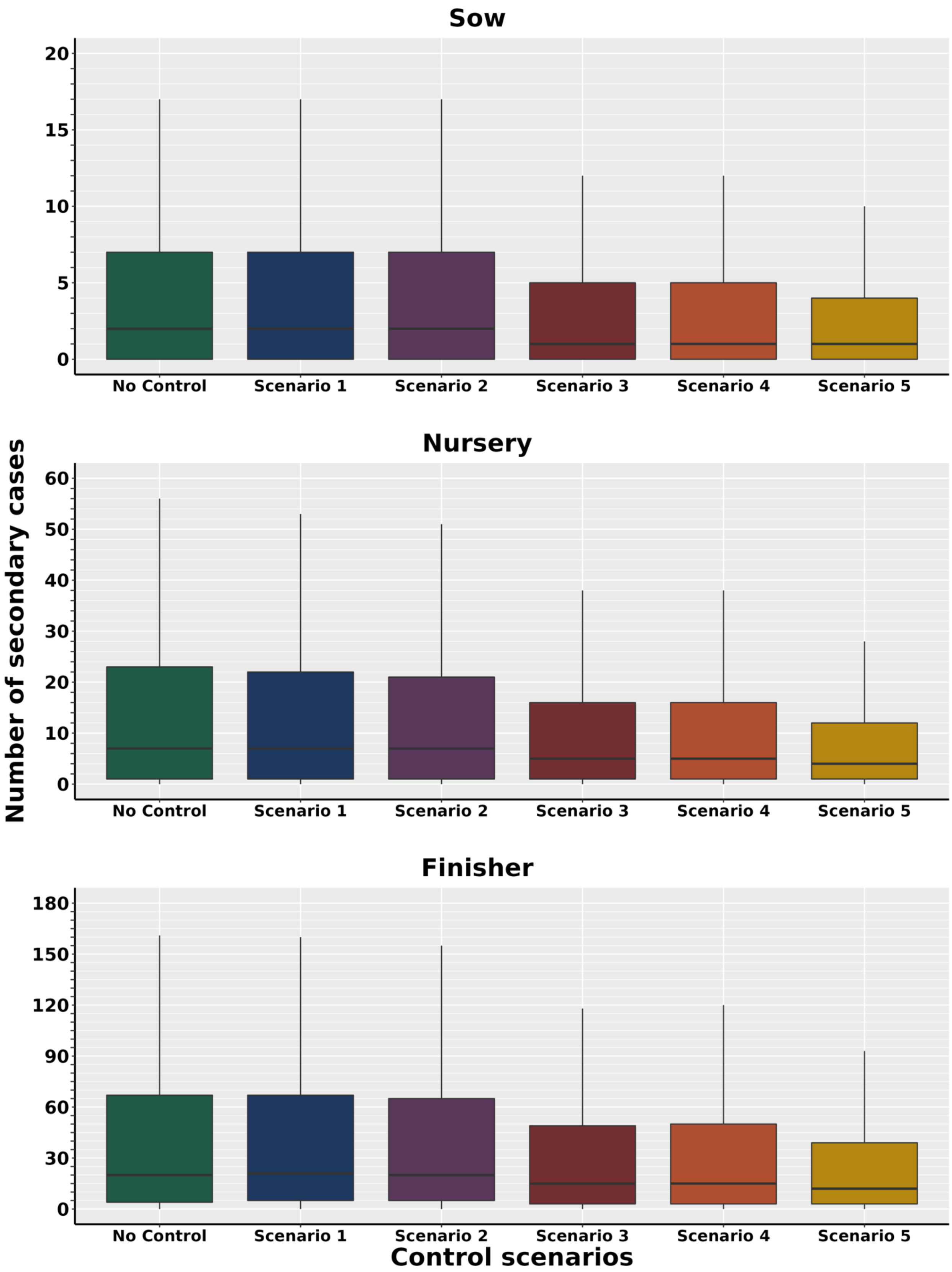
Sow
20
15
10
5
0
No Control
Scenario 1
Scenario 2
Scenario 3
Scenario 4
Scenario 5
Nursery
60
50
40
30
20
10
0
No Control
Scenario 1
Scenario 2
Scenario 3
Scenario 4
Scenario 5
Finisher
180
150
120
90
60
30
0
No Control
Scenario 1
Scenario 2
Scenario 3
Scenario 4
Scenario 5
Number of secondary cases
Control scenarios

373

**Figure S19. Total secondary cases at day 60 of the epidemic across the model simulations for each control scenario.** The boxplots present the range, interquartile range, and median. Outliers were retained in the dataset but are not shown. Only model simulations which resulted in secondary cases were used to create this figure.

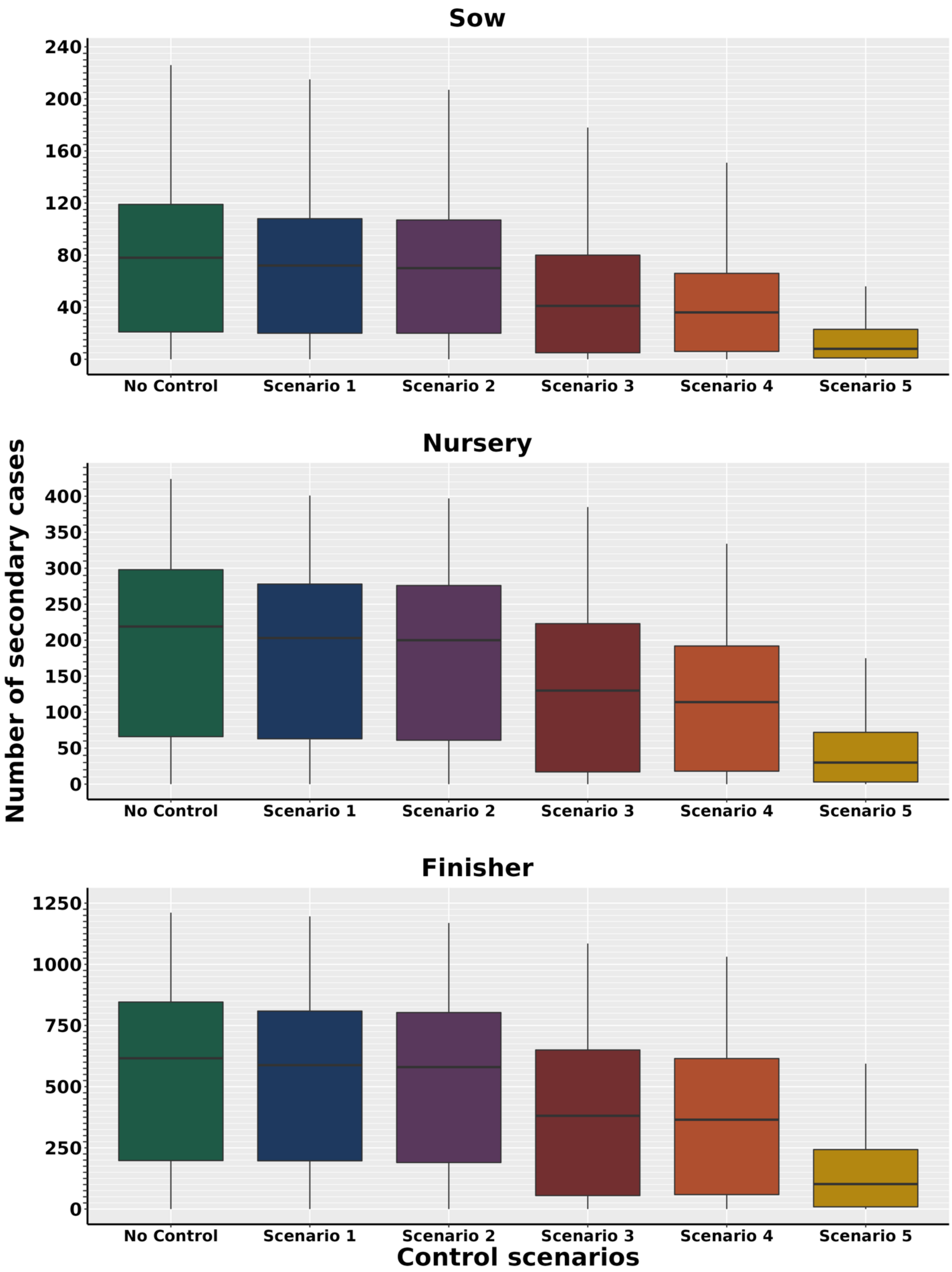
Sow
Nursery
Finisher
Number of secondary cases
Control scenarios
No Control
Scenario 1
Scenario 2
Scenario 3
Scenario 4
Scenario 5
0
40
80
120
160
200
240
50
100
150
250
300
350
400
500
750
1000
1250

**Figure S20. Total secondary cases by day 140 of the epidemic across the model simulations for each control scenario.** The boxplots present the range, interquartile range, and median. Outliers were retained in the dataset but are not shown. Only model simulations which resulted in secondary cases were used to create this figure.

*Depopulation*

We also estimated the direct cost of depopulation (i.e., the cost of compensation of producers) in each simulated control scenario, by using USDA indemnity tables (Supplementary Material Section E, Table S8). Over 60 days of outbreak, the lowest direct cost was observed in scenario two with an estimated cost of USD $400,702 while the maximum was observed in scenario five with a direct cost of USD $2,869,936. A detailed description of the direct costs is presented in Supplementary Material Section E, Table S9. Over 140 days the median number of depopulated animals in scenario five was 495,619 (IQR: 28,670-1,410,459) equating to a cost of USD $79.2 million (Supplementary Material Section E, Table S10).

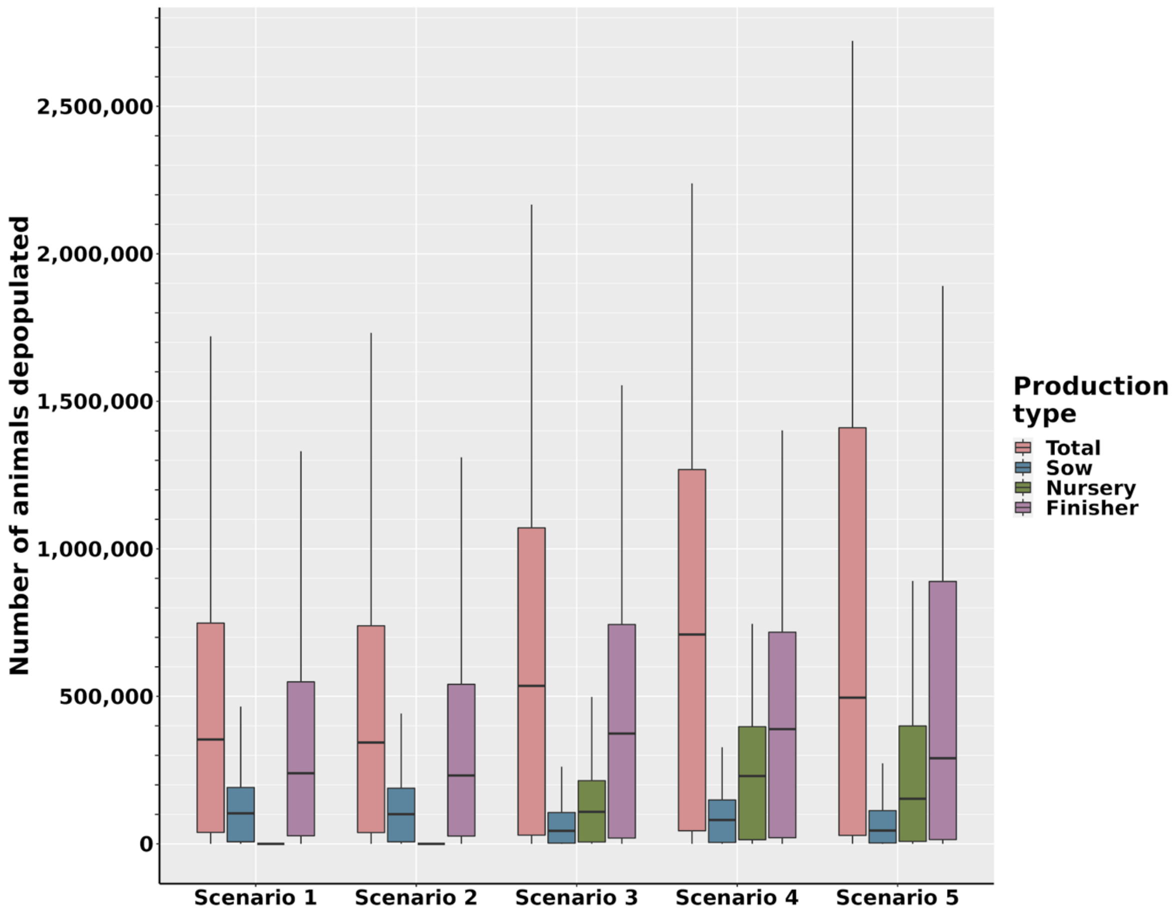


**Figure S21. Boxplots showing the number of animals depopulated under the five control and eradication scenarios for each farm type at 140 days post seeded infection.**

**Table S8. Indemnity prices of different production types per animal**

| **Production Type** | **Price (USD)[a]** | **Source** |
|---|---|---|
| Sow[b] | 270.83 - 352.10 | (USDA, 2022a) |
| Nursery[c] | 75.22 | (USDA, 2022a) |
| Finisher[d] | 128.31 - 178.27 | (USDA, 2022a) |
| Boar[e] | 270.83 - 352.10 | (USDA, 2022a) |

| | | |
|---|---|---|
| Gilt[f] | 178.27 - 270.83 | (USDA, 2022a) |

a. Price based on the weight (lbs) of the animal.

b. Weight of sows extracted from (USDA, 2022b)

c. Weight of nursery swine extracted from (USDA, 2022a)

d. Weight of finisher swine extracted from (National Pork Board, 2021; USDA, 2022a)

e. Weight of boar studs extracted from (Whitney and Baidoo, 2015)

f. Weight of replacement gilts extracted from (Hoar et al., 2015; PIC, 2020)

**Table S9. The estimated direct cost of expected depopulated animals after 60 days of outbreak in U.S. dollars.**

| **Production type** | **Scenario 1** | **Scenario 2** | **Scenario 3** | **Scenario 4** | **Scenario 5** |
|---|---|---|---|---|---|
| Sow | $162,768.83 - $211,612.100 | $144,081.56 - $187,317.20 | $162,498 - $211,260 | $324,996 - $422,520 | $595,826 - $774,620 |
| Nursery | $0 | $0 | $0 | $288,844.80 | $526,540 |
| Finisher | $264,318.60 - $367,236.20 | $256,620 - $356,540 | $579,704.58 - $805,423.86 | $513,240 - $713,080 | $1,129,128 - $1,568,776 |
| Boar Stud | $0 | $0 | $0 | $0 | $0 |
| Gilt | $0 | $0 | $0 | $0 | $0 |
| Total | $427,087.43 - $578,848.30 | $400,701.56 - $543,857.20 | $742,202.58 - $1,016,683.86 | $1,127,080.80 - $1,424,444.80 | $2,251,494 - $2,869,936 |

**Table S10. The estimated direct cost of expected depopulated animals after 140 days of outbreak in U.S. dollars**

| Production type | Scenario 1 | Scenario 2 | Scenario 3 | Scenario 4 | Scenario 5 |
|---|---|---|---|---|---|
| Sow | $28,027,925.87 - $36,438,476.90 | $27,259,039.50 - $35,438,865 | $11,950,102.92 - $15,536,060.92 | $21,940,209.13 - $28,523,973.10 | $12,241,516 - $15,914,920 |
| Nursery | $0 | $0 | $8,167,011.50 | $17,294,507.18 | $11,507,381.26 |
| Finisher | $30,721,776.54 - $42,683,899.20 | $29,735,585.88 - $41,313,715.96 | $47,984,860.56 - $66,668,701.50 | $49,907,072.67 - $69,339,364.39 | $37,254,551.88 - $51,760,337.96 |
| Boar Stud | $0 | $0 | $0 | $0 | $0 |
| Gilt | $0 | $0 | $0 | $0 | $0 |
| Total | $58,749,702.41 - $79,122,376.10 | $56,994,625.38 - $76,752,580.96 | $68,101,974.98 - $90,371,773.40 | $89,141,788.98 - $115,157,844.70 | $61,003,449.14 - $79,182,639.22 |

*Diagnostic testing*

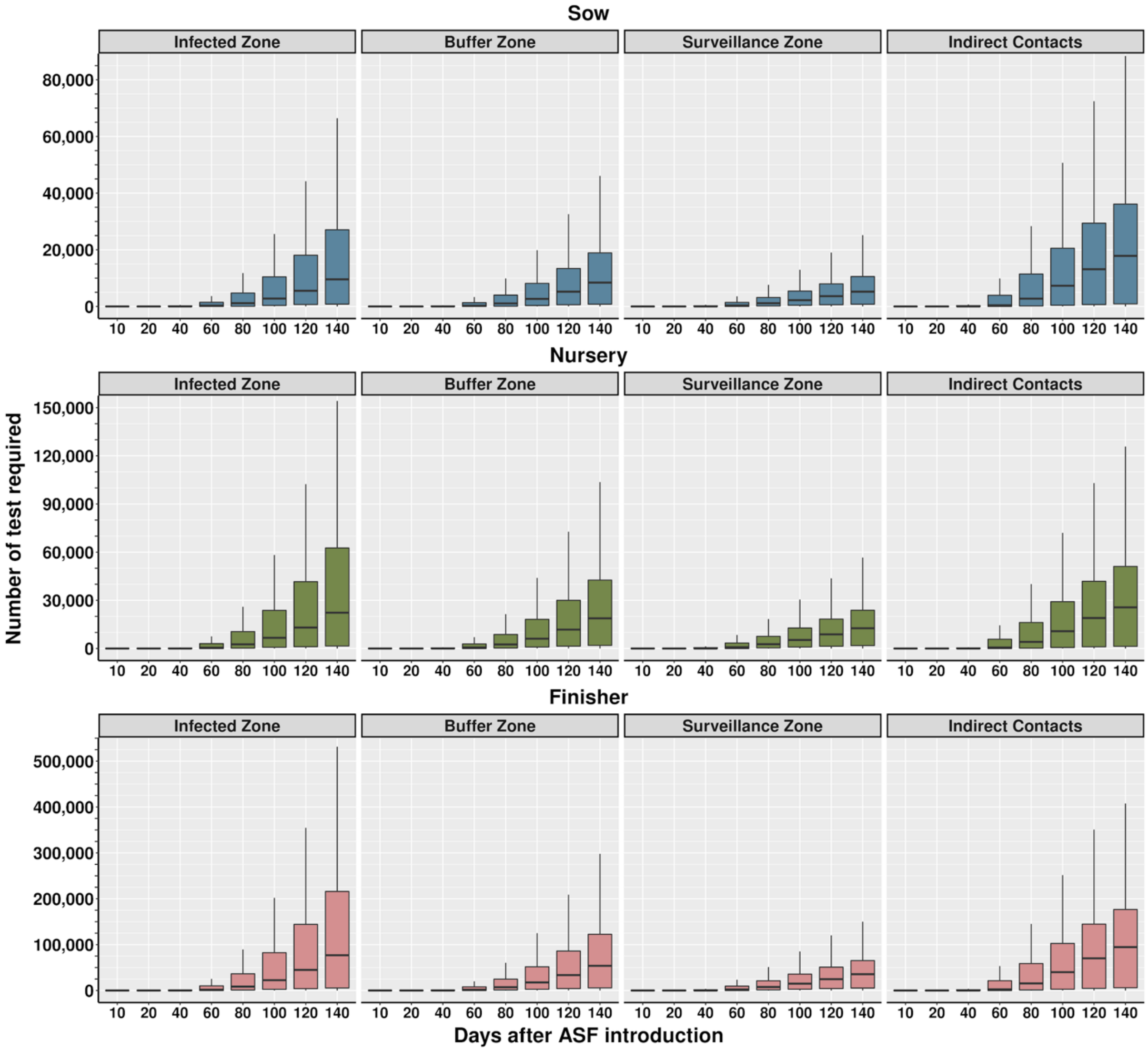


**Figure S22. The cumulative number of diagnostic tests required for indirect contacts, infected zones, buffer zones and surveillance zones during control scenario five across 140 days of outbreaks, when samples are pooled in groups of five.** Due to the delay in detection implemented in the model, testing was not required earlier than day 13 of the outbreak. Further details are available in section 2.5.

*Movement permits required over 140 days*

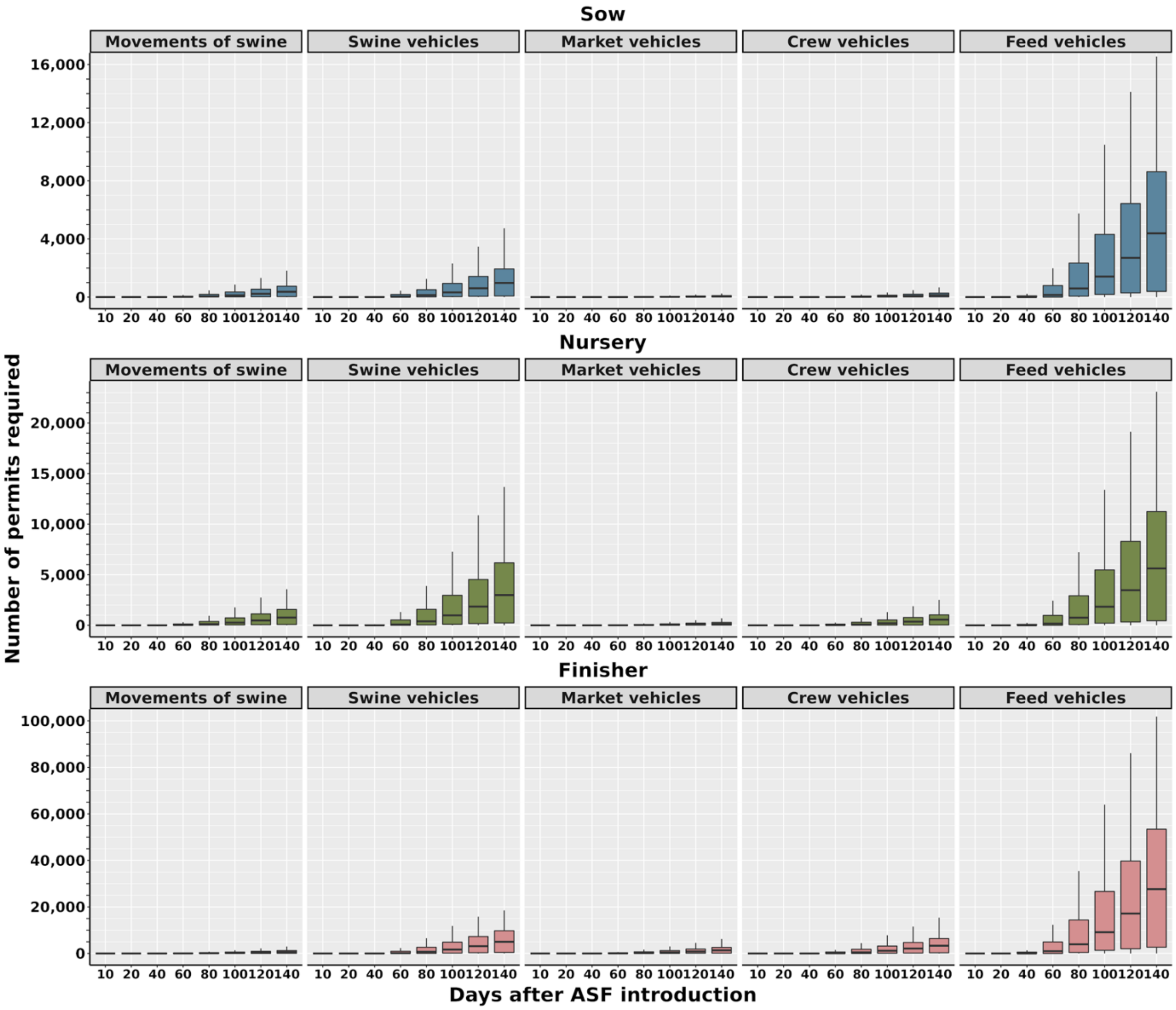


**Figure S23. Median daily number of permits required for movements to or from infected zones, buffer zones and indirect contacts, during control scenario five across 140 days of**

**outbreaks.** Due to the delay in detection implemented in the model, the earliest date that permits were required was day 13 of the outbreak, see the methods section 2.5 for further details.

**Section F.**

*Additional data*

**Table S9. Characterization of animal and vehicle movements**

| **Movement type** | **Total number of movements over calendar year** | **Median daily movement** | **Daily movement interquartile range** |
|---|---|---|---|
| Swine movements | 37,501 | 105.0 | 86.00 – 142.00 |
| Swine vehicle movements | 118,509 | 29.5 | 0.00 – 659.25 |
| Market vehicle movements | 20,948 | 40.0 | 7.00 – 92.00 |
| Crew vehicle movements | 44,296 | 93.5 | 23.00 – 198.75 |
| Feed vehicle movements | 425,464 | 1258.0 | 257.00 – 1876.50 |

**References**

1. Andraud, M., T. Halasa, A. Boklund, and N. Rose, 2019: Threat to the French Swine Industry of African Swine Fever: Surveillance, Spread, and Control Perspectives. *Frontiers in Veterinary Science* 6, 248, DOI: 10.3389/fvets.2019.00248.
2. Boklund, A., S. Dhollander, T. Chesnoiu Vasile, J.C. Abrahantes, A. Bøtner, A. Gogin, L.C. Gonzalez Villeta, C. Gortázar, S.J. More, A. Papanikolaou, H. Roberts, A. Stegeman, K. Ståhl, H.H. Thulke, A. Viltrop, Y. Van der Stede, and S. Mortensen, 2020: Risk factors for

African swine fever incursion in Romanian domestic farms during 2019. *Sci Rep* 10, 10215, DOI: 10.1038/s41598-020-66381-3.

3. Bradhurst, R., G. Garner, S. Roche, R. Iglesias, N. Kung, B. Robinson, S. Willis, M. Cozens, K. Richards, B. Cowled, M. Oberin, C. Tharle, S. Firestone, and M. Stevenson, 2021: Modelling the spread and control of African swine fever in domestic and feral pigs (Technical report). Centre of Excellence for Biosecurity Risk Analysis.
4. Cannon, R.M., 2001: Sense and sensitivity — designing surveys based on an imperfect test. *Preventive Veterinary Medicine* 49, 141–163, DOI: 10.1016/S0167-5877(01)00184-2.
5. Carlson, J., M. Fischer, L. Zani, M. Eschbaumer, W. Fuchs, T. Mettenleiter, M. Beer, and S. Blome, 2020: Stability of African Swine Fever Virus in Soil and Options to Mitigate the Potential Transmission Risk. *Pathogens* 9, 977, DOI: 10.3390/pathogens9110977.
6. Chenais, E., K. Depner, V. Guberti, K. Dietze, A. Viltrop, and K. Ståhl, 2019: Epidemiological considerations on African swine fever in Europe 2014–2018. *Porcine Health Management* 5, 6, DOI: 10.1186/s40813-018-0109-2.
7. De Lorenzi, G., L. Borella, G.L. Alborali, J. Prodanov-Radulović, M. Štukelj, and S. Bellini, 2020: African swine fever: A review of cleaning and disinfection procedures in commercial pig holdings. *Research in Veterinary Science* 132, 262–267, DOI: 10.1016/j.rvsc.2020.06.009.
8. Fodor, J.T., F. Jánoska, and A. Farkas, 2015: The comparative analysis of the habitat use of wild boar in different Romanian habitats (partial results). *Proceedings of the Biennial International Symposium. Forest and sustainable development, Brașov, Romania, 24-25th October 2014*365–370.
9. Galvis, J.A., C.A. Corzo, and G. Machado, 2022: Modelling and assessing additional transmission routes for porcine reproductive and respiratory syndrome virus: Vehicle movements and feed ingredients. *Transbounding Emerging Dis*tbed.14488, DOI: 10.1111/tbed.14488.
10. Gebhardt, J.T., S.S. Dritz, C.G. Elijah, C.K. Jones, C.B. Paulk, and J.C. Woodworth, 2021: Sampling and detection of African swine fever virus within a feed manufacturing and swine production system. *Transboundary and Emerging Diseases*DOI: 10.1111/tbed.14335.
11. Halasa, T., A. Bøtner, S. Mortensen, H. Christensen, N. Toft, and A. Boklund, 2016: Simulating the epidemiological and economic effects of an African swine fever epidemic in industrialized swine populations. *Veterinary Microbiology* 193, 7–16, DOI: 10.1016/j.vetmic.2016.08.004.
12. Hartig, F., J.M. Calabrese, B. Reineking, T. Wiegand, and A. Huth, 2011: Statistical inference for stochastic simulation models – theory and application. *Ecology Letters* 14, 816–827, DOI: 10.1111/j.1461-0248.2011.01640.x.

13. Hu, B., J.L. Gonzales, and S. Gubbins, 2017: Bayesian inference of epidemiological parameters from transmission experiments. *Sci Rep* 7, 16774, DOI: 10.1038/s41598-017-17174-8.

14. Iglesias, I., M.J. Muñoz, F. Montes, A. Perez, A. Gogin, D. Kolbasov, and A. de la Torre, 2016: Reproductive Ratio for the Local Spread of African Swine Fever in Wild Boars in the Russian Federation. *Transboundary and Emerging Diseases* 63, e237–e245, DOI: 10.1111/tbed.12337.

15. Jiang, C., Y. Sun, F. Zhang, X. Ai, X. Feng, W. Hu, X. Zhang, D. Zhao, Z. Bu, and X. He, 2021: Viricidal activity of several disinfectants against African swine fever virus. *Journal of Integrative Agriculture* 20, 3084–3088, DOI: 10.1016/S2095-3119(21)63631-6.

16. Juszkiewicz, M., M. Walczak, N. Mazur-Panasiuk, and G. Woźniakowski, 2020: Effectiveness of Chemical Compounds Used against African Swine Fever Virus in Commercial Available Disinfectants. *Pathogens* 9, 878, DOI: 10.3390/pathogens9110878.

17. Li, X., Z. Hu, M. Fan, W. Wu, W. Gao, L. Bian, W. Liu, X. Tian, X. Jiang, and Z.J. Yan, 2022: Evidence of aerosol transmission of African swine fever virus in piggeries under field conditions: a case report (preprint). In Review.

18. Machado, G., C. Vilalta, M. Recamonde-Mendoza, C. Corzo, M. Torremorell, A. Perez, and K. VanderWaal, 2019: Identifying outbreaks of Porcine Epidemic Diarrhea virus through animal movements and spatial neighborhoods. *Sci Rep* 9, 457, DOI: 10.1038/s41598-018-36934-8.

19. Malladi, S., A. Ssematimba, P.J. Bonney, K.M.S. Charles, T. Boyer, T. Goldsmith, E. Walz, C. Cardona, and M.R. Culhane, 2022: Predicting the Time to Detect Moderately Virulent African Swine Fever Virus in Finisher Swine Herds Using a Stochastic Disease Transmission Model. *BMC Veterinary Research (PREPRINT)*DOI: 10.21203/rs.3.rs-716595/v1.

20. Mazur-Panasiuk, N., J. Żmudzki, and G. Woźniakowski, 2019: African Swine Fever Virus – Persistence in Different Environmental Conditions and the Possibility of its Indirect Transmission. *J Vet Res* 63, 303–310, DOI: 10.2478/jvetres-2019-0058.

21. Minter, A., and R. Retkute, 2019: Approximate Bayesian Computation for infectious disease modelling. *Epidemics* 29, 100368, DOI: 10.1016/j.epidem.2019.100368.

22. Olesen, A.S., L. Lohse, A. Boklund, T. Halasa, G.J. Belsham, T.B. Rasmussen, and A. Bøtner, 2018: Short time window for transmissibility of African swine fever virus from a contaminated environment. *Transboundary and Emerging Diseases* 65, 1024–1032, DOI: 10.1111/tbed.12837.

23. Olesen, A.S., L. Lohse, A. Boklund, T. Halasa, C. Gallardo, Z. Pejsak, G.J. Belsham, T.B. Rasmussen, and A. Bøtner, 2017: Transmission of African swine fever virus from infected

pigs by direct contact and aerosol routes. *Veterinary Microbiology* 211, 92–102, DOI: 10.1016/j.vetmic.2017.10.004.

24. Pepin, K.M., A. Golnar, and T. Podgórski, 2021: Social structure defines spatial transmission of African swine fever in wild boar. *Journal of The Royal Society Interface* 18, 20200761, DOI: 10.1098/rsif.2020.0761.
25. Sanson, R.L., 1994: The epidemiology of foot-and-mouth disease: Implications for New Zealand. *New Zealand Veterinary Journal* 42, 41–53, DOI: 10.1080/00480169.1994.35785.
26. Sisson, S.A., Y. Fan, and M.M. Tanaka, 2007: Sequential Monte Carlo without likelihoods. *Proceedings of the National Academy of Sciences* 104, 1760–1765, DOI: 10.1073/pnas.0607208104.
27. USDA, S. reports, 2022 (6. June): Swine Summary Reports, Agricultural Marketing Service [Online] Available at https://www.ams.usda.gov/market-news/swine-reports (accessed June 6, 2022).
28. Wilkinson, P.J., and A.I. Donaldson, 1977: Transmission studies with African swine fever virus: The early distribution of virus in pigs infected by airborne virus. *Journal of Comparative Pathology* 87, 497–501, DOI: 10.1016/0021-9975(77)90038-X.
